# A conserved cell-pole determinant organizes proper polar flagellum formation

**DOI:** 10.1101/2023.09.20.558563

**Authors:** Erick Eligio Arroyo-Pérez, John C. Hook, Alejandra Alvarado, Stephan Wimmi, Timo Glatter, Kai M. Thormann, Simon Ringgaard

## Abstract

The coordination of cell cycle progression and flagellar synthesis is a complex process in motile bacteria. In γ-proteobacteria, the localization of the flagellum to the cell pole is mediated by the SRP-type GTPase FlhF. However, the mechanism of action of FlhF, and its relationship with the cell pole landmark protein HubP remain unclear. In this study, we discovered a novel protein called FipA that is required for normal FlhF activity and function in polar flagellar synthesis. We demonstrated that membrane-localized FipA interacts with FlhF and is required for normal flagellar synthesis in *Vibrio parahaemolyticus*, *Pseudomonas putida*, and *Shewanella putrefaciens*, and it does so independently of the polar localization mediated by HubP. FipA exhibits a dynamic localization pattern and is present at the designated pole before flagellar synthesis begins, suggesting its role in licensing flagellar formation. This discovery provides insight into a new pathway for regulating flagellum synthesis and coordinating cellular organization in bacteria that rely on polar flagellation and FlhF-dependent localization.

## Introduction

Many cellular processes depend on a specific spatiotemporal organization of its components. The DNA replication machinery, cell division proteins and motility structures are some examples of elements that need to be positioned at a particular site during the cell cycle in a coordinated manner to ensure adequate cell division or proper function of the structures. To carry out this task, many cellular components of bacteria have a polar organization, *i. e.* they are asymmetrically positioned inside the cell, which is particularly apparent in monopolarly flagellated bacteria. In the marine bacteria *Vibrio*, which constitutively express a single polar flagellum (McCarter 1995; Kim and McCarter 2000), it is crucial to coordinate the localization and timing of flagellar components, in order to guarantee that newly born cells only produce a flagellum in timing with cell division. This process is coordinated with the chromosome replication and the chemotaxis clusters through a supramolecular hub that is tethered at the old cell pole (Takekawa et al. 2016; Yamaichi et al. 2012). This organization depends on ATPases of the ParA/MinD family, which regulate the migration of the hub components to the new cell pole as the cell cycle progresses. In this way, the new cell has a copy of the chromosome and a set of chemotaxis clusters positioned next to what will be the site for the new flagellum. Central to this process is the protein HubP, which recruits to the pole the three ATPases responsible for the localization of these organelles: ParA1 for the chromosome (Fogel and Waldor 2006), ParC for the chemotaxis clusters (Ringgaard et al. 2011) and FlhG for the flagellum (Correa, Peng, and Klose 2005; Arroyo-Pérez and Ringgaard 2021). Homologs of HubP occur in several species, e.g., *Pseudomonas* (where it is called FimV), *Shewanella* or *Legionella*, where they similarly act as organizers (hubs) of the cell pole (Wehbi et al. 2011; Rossmann et al. 2015; Coil and Anné 2010).

The bacterial flagellum is a highly intricate and complex subcellular structure. It is composed of around 25 different proteins, which have to be assembled in different stoichiometries in a spatiotemporally coordinated manner (Macnab 2003; Chevance and Hughes, 2008). Although there may be significant differences among species, flagella in general are composed of a motor attached to the cytoplasmic membrane, a rod connecting it to the extracellular space, a hook and a filament. The basal body is composed of a cytoplasmic C-ring or switch complex (Francis et al. 1994), which receives the signal from the chemotaxis proteins via binding to phosphorylated CheY. The C-ring is connected to the MS-ring, which is embedded in the cytoplasmic membrane (Homma et al. 1987; Ueno, Oosawa, and Aizawa 1992). Attached to it, on the cytoplasmic side, is the export apparatus, which allows secretion of rod, hook and filament components (Minamino 2014). Flagella can also be sheathed, if an extension of the outer membrane covers the filament, as is the case in many *Vibrio* species (Chen et al. 2017).

The positioning of the flagella and control of flagellar numbers per cell are mediated in many bacteria by the interplay between two flagellar regulators, FlhF and FlhG. Studies in a wide variety of proteobacteria indicate that FlhG is a negative regulator that restricts the number of flagella that are synthesized. In organisms such as strains of *Vibrio*, *Shewanella putrefaciens* and *Pseudomonas aeruginosa*, the absence of MinD-like ATPase FlhG results in hyper-flagellated cells (Kusumoto et al. 2006; Arroyo-Pérez and Ringgaard 2021; Schuhmacher et al. 2015; Blagotinsek et al., 2020; Campos-García et al. 2000; Murray and Kazmierczak 2006). In contrast, in polarly flagellated species such as *Campylobacter jejuni, Vibrio cholerae, Vibrio parahaemolyticus*, *P. aeruginosa, S. putrefaciens* and *Shewanella oneidensis*, the SRP-type GTPase FlhF is a positive regulator of flagellum synthesis and necessary for proper localization. A deletion of *flhF* results in reduced number and mis-localization of flagella (Hendrixson and Dirita 2003; Correa, Peng, and Klose 2005; Arroyo-Pérez and Ringgaard 2021; Pandza et al. 2000; Rossmann et al. 2015; Gao et al. 2015, Navarrete et al, 2019).

The current model predicts that GTP-bound dimeric FlhF (Bange et al. 2007; Kondo et al. 2018) localizes to the cell pole to where it recruits the initial flagellar building blocks (Green et al., 2009). A long-standing question is how FlhF recognizes and localizes to the designated cell pole. In *P. aeruginosa* and *V. alginolyticus* it was demonstrated that FlhF localized polarly upon ectopic production in the absence of the flagellar master regulator FlrA (Green et al. 2009; Kondo, Homma, and Kojima 2017). It was therefore speculated that FlhF assumes its polar localization in the absence of other flagellar proteins or even without any further additional protein factors. The underlying mechanism, however, remains enigmatic, given that neither in its monomeric nor its dimeric form, FlhF possesses any regions that would indicate a membrane association. In a recent study it was shown that in the polarly flagellated gammaproteobacterium *Shewanella putrefaciens* FlhF binds the C-ring protein FliG via a specific region at the very N-terminus of FlhF. The FlhF-FliG complex is then recruited to the designated cell pole by HubP, where FlhF-bound FliG captures the transmembrane protein FliF and promotes formation of the MS-ring. This forms the scaffold from where further flagellar synthesis can occur (Dornes et al, 2024). However, in the absence of HubP, the majority of cells still forms normal polar flagella, indicating that HubP is not the only polarity factor in this process (Rossmann et al, 2015).

Here, we provide evidence demonstrating that FlhF does not localize independently. Instead, we show that FipA, a small integral membrane protein with a domain of unknown function, facilitates the recruitment of FlhF to the membrane at the cell pole and, at least in some species, it acts in concert with the polar landmark protein HubP/FimV. Using *V. parahaemolyticus* and two additional polarly flagellated γ-proteobacteria, the monopolarly flagellated *S. putrefaciens* and the lophotrichous *P. putida*, we show that FipA universally mediates recruitment of FlhF to the designated cell pole. The spatiotemporal localization behavior of FipA as well as its relationship to FlhF, support its role as a licensing protein that enables flagellum synthesis.

## Results

### Identification of an FlhF protein interaction partner: FipA

In order to identify potential factors required for FlhF function and its recruitment to the cell pole, we performed affinity purification of a superfolder green fluorescent protein (sfGFP)-tagged FlhF (FlhF-sfGFP) ectopically expressed in wild-type *V. parahaemolyticus* cells followed by shotgun proteomics using liquid chromatography-tandem mass spectrometry analysis (LC-MS/MS). Among the proteins that were significantly enriched in FlhF-sfGFP purifications, eight were structural components of the flagellum (**Fig. 1A**; **Supplementary Table S1**), but also a non-flagellar protein, VP2224, was significantly co-purified with FlhF (**Fig. 1A; Supplementary Table S1**). The homologue of VP2224 in *V. cholerae*, there named FlrD, was previously shown to exert the same activating role on flagellar gene regulation as FlhF (Moisi et al. 2009), suggesting a function related to FlhF. Notably, FlhF was also significantly co-purified in the reciprocal co-IP-MS/MS experiment using VP2224-sfGFP as bait (**Fig. 1B**), suggesting a direct or indirect interaction of VP2224 and FlhF. To further investigate the co-purification data, bacterial two-hybrid (BACTH) assays in the heterologous host *E. coli* were carried out and suggested a direct interaction between FlhF and VP2224. FlhF and VP2224 were also found to self-interact (**Fig. 1E**). Additionally, these data indicated that FlhF and VP2224 form an interaction complex in both the native and a heterologous host organism. Thus, we identified VP2224 as a novel interaction partner of FlhF and named it FipA for Flh**<u>F I</u>**nteraction **<u>P</u>**artner **<u>A</u>**.

**Figure 1.**
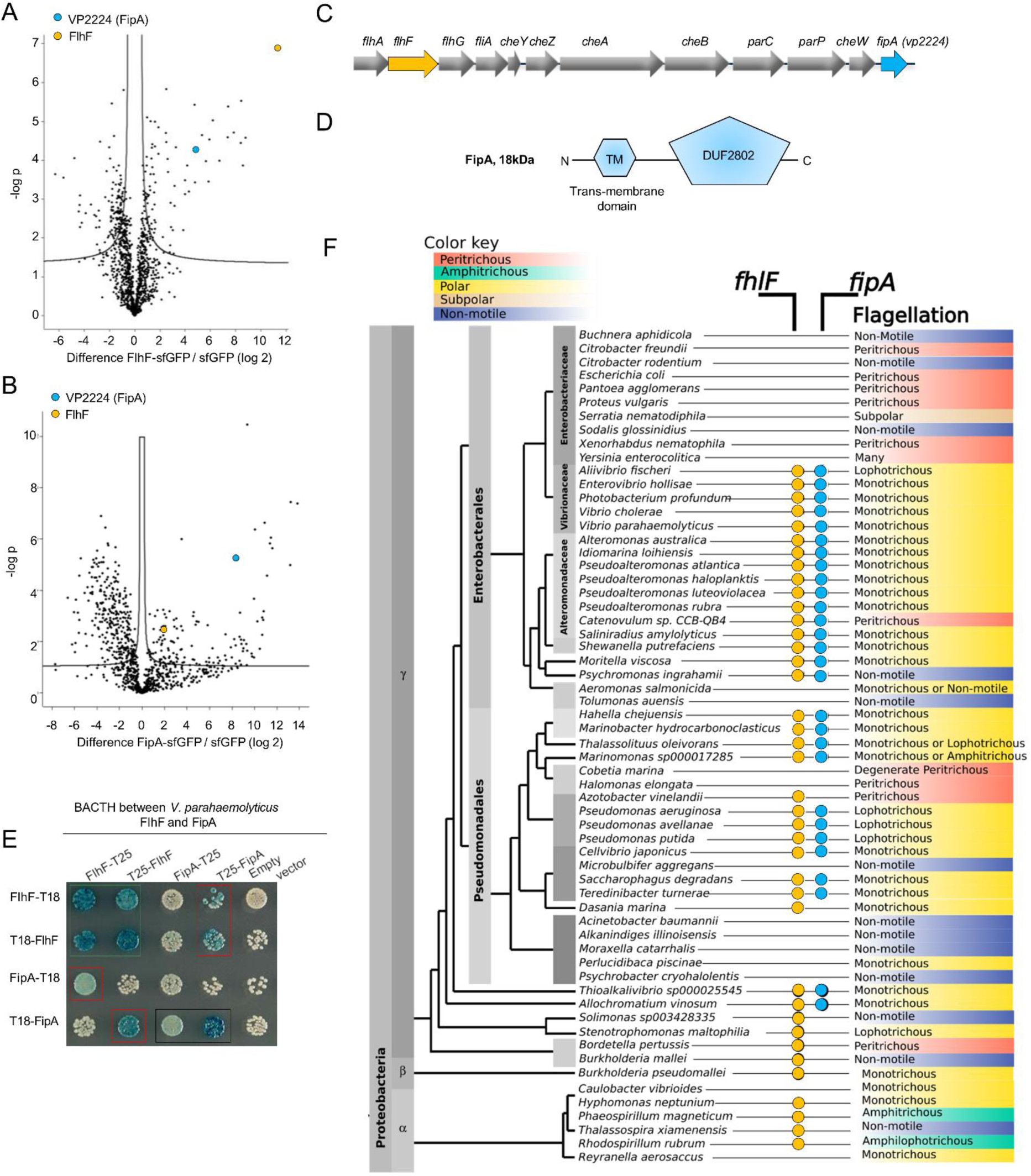
FipA constitutes a new family of FlhF interaction partners. Volcanoplots representing Log-ratios versus significance values of proteins enriched in **(A)** FlhF-sfGFP or **(B)** FipA-sfGFP purifications using shotgun proteomics and liquid chromatography-mass spectrometry; sfGFP was used as control. The full list of pulled-down proteins can be found in the Supplementary Tables S1 and S2. **C)** Organization of the flagellar/chemotaxis gene region encoding FlhF and FipA in *V. parahaemolyticus*. **D)** Domain organization of FipA (TM, transmembrane region; DUF, domain of unknown function). **E)** Bacterial two-hybrid confirming the interaction between FipA and FlhF from *V. parahaemolyticus*. The indicated proteins (FipA, FlhF) were fused N- or C-terminally to the T18- or T25-fragment of the *Bordetella pertussis* adenylate cyclase. In vivo interaction of the fusion proteins in *Escherichia coli* is indicated by blue color. The corresponding assay in *P. putida* and *S. putrefaciens* is displayed in **Supplementary Figure 2**. **F)** Dendrogram of γ-proteobacteria, indicating the presence of FlhF or FipA homologues and the corresponding flagellation pattern. An extended version and sources are available in **Supplementary Table 2**.

### FipA constitutes a new family of FlhF interaction partners

The gene encoding FipA is located immediately downstream of the flagellar operon that encodes FlhA, FlhF, FlhG, FliA and the chemotaxis proteins (**Fig. 1C**). In *V. parahaemolyticus*, FipA consists of 163 amino acids with a molecular mass of 18.4 kDa. *In silico* analysis and membrane topology mapping predicted that FipA consists of a short periplasmic N-terminal part, consisting of amino acids 1-5, followed by a transmembrane region (6-28), a cytoplasmic part harboring a coiled region (amino acids 31-58) and a domain of unknown function, DUF2802, positioned in the C-terminal half of the protein (from amino acids 68-135) (**Fig. 1D; Supplementary Fig. S1**).

InterPro database analyzes found that FipA homologues (*i. e.* membrane proteins consisting of a single cytoplasmic DUF2802 repeat) are widespread among γ-proteobacteria. Exceptions are the *Enterobacteriaceae*, which do not possess any copies of either *fipA* nor of *flhF* and *flhG*. Actually, FipA is only present in genomes that also encode FlhF and FlhG (**Fig. 1F**), which prompted the question of whether FipA is involved in regulating the flagellation pattern in concert with the FlhF-FlhG system. By including in our analysis the flagellation pattern reported in the literature, we found that the species that encode FipA are all polar flagellates, either monotrichous or lophotrichous (**Fig. 1F**). FipA homologues are absent from bacteria that use the FlhF-FlhG system to produce different flagellation patterns, like the peritrichous *Bacillus*, the amphitrichous ε-Proteobacteria or Spirochetes (**Supplementary Table S2**). In the α-Proteobacteria, where FlhF homologues are only present in a few species, FipA is absent as well (**Fig. 1F**). Based on these analyzes, we hypothesized that FipA represents a new family of FlhF interaction partners important for the γ-proteobacteria. To test this hypothesis, we decided to analyze FipA-FlhF interaction in addition in the distantly related and lophotrichously flagellated *Pseudomonas putida* and the monotrichous *Shewanella putrefaciens*. BACTH analysis showed that the FipA orthologue from both *P. putida* (*Pp*FipA, PP_4331) and *S. putrefaciens* (*Sp*FipA, SputCN32_2550) interact with FlhF of their respective species and that they self-interact (**Supplementary Fig. S2**) – a result similar to that of FipA from *V. parahaemolyticus* (*Vp*FipA). This supported our hypothesis and indicated that FipA has a general function as an FlhF interaction partner, thus constituting a new class of FlhF interaction partners.

### FipA is required for proper swimming motility and flagellum formation

To explore the role of FipA with respect to flagellation, we generated mutant strains with individual deletions of the *fipA* and *flhF* genes. Strikingly, absence of FipA in *V. parahaemolyticus* completely abolished swimming motility in soft-agar medium to the same degree as cells lacking FlhF (**Fig. 2A**). Furthermore, single-cell tracking of planktonic *V. parahaemolyticus* cells confirmed that cells lacking FipA were completely non-motile and behaved identical to cells lacking FlhF (**Fig. 2B**). Importantly, ectopic expression of FipA in the *ΔfipA* strain restored the strain’s swimming ability to wild-type levels (**Fig. 2B**), further supporting that it is the deletion of *fipA* that results in the phenotype and not polar effects resulting from the *fipA* deletion. The C-terminal FipA-sfGFP fusion used throughout this paper also restored the phenotype (**Fig. 2B**), indicating that the fusion protein is fully functional. The swimming phenotypes in the absence of FipA could result from defects in either flagellum assembly or in the flagellar motor performance. To differentiate between these possibilities, we examined *V. parahaemolyticus* by transmission electron microscopy (TEM) (**Fig. 2D, E**). Planktonic cells lacking either FlhF or FipA, showed a complete absence of flagella on the bacterial surface, while a single polar flagellum was observed in ∼50 % of wild-type cells (**Fig. 2D, E**).

**Figure 2.**
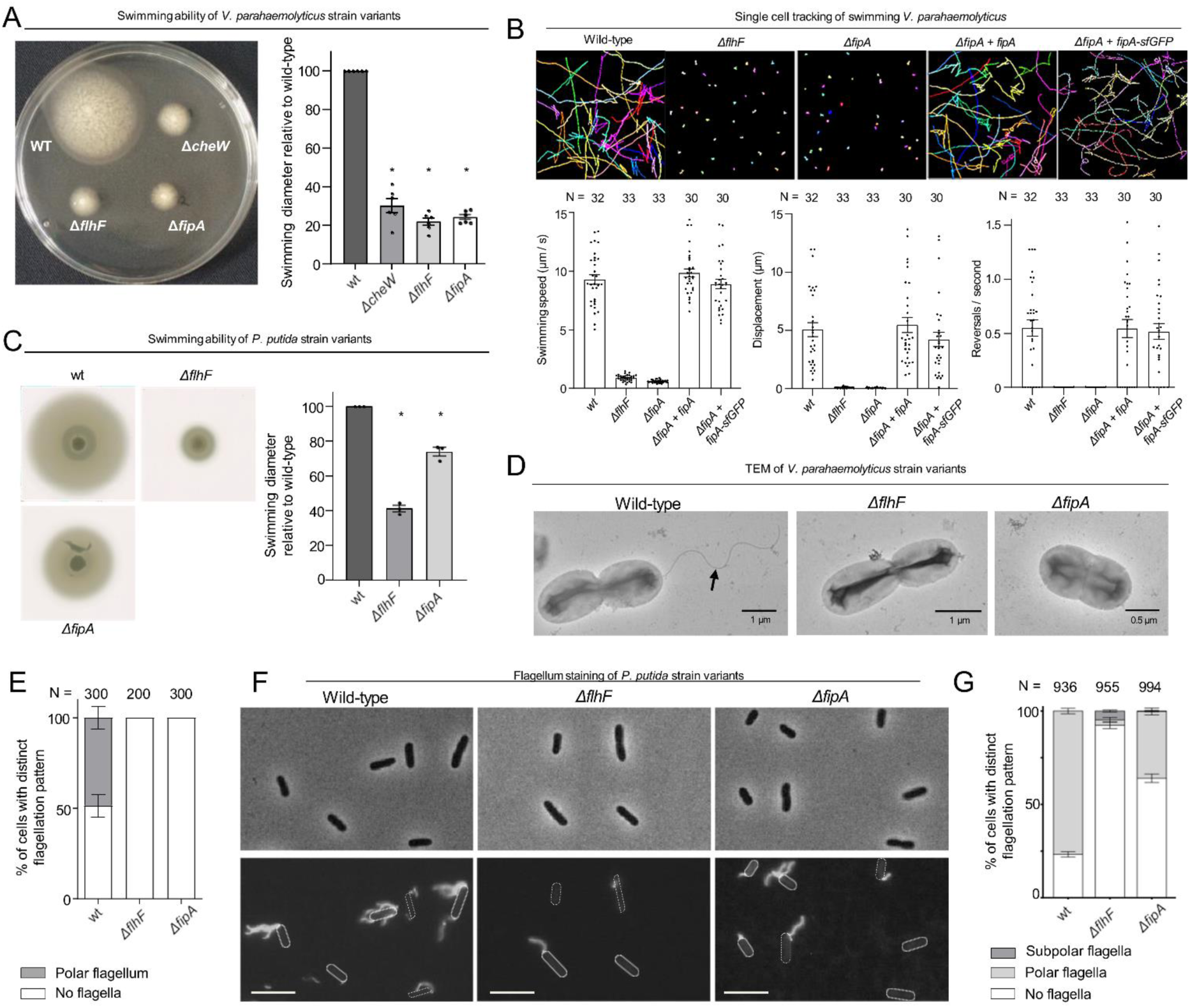
FipA is required for correct flagellum formation. **(A, C)** Representative soft-agar swimming assay of *V. parahaemolyticus* (A) or *P. putida* (C) strains (left panels) and the corresponding quantification (right panels). For the latter, the halo diameter measurements were normalized to the halo of the wild type on each plate. Data presented are from six (A) or three (C) independent replicates, asterisks represent a p-value <0.05 (according to ANOVA + Tukey tests). **(B)** Single-cell tracking of *V. parahaemolyticus*. Shown are representative swimming trajectories and quantification of swimming speed, total displacement and reversal rate. N indicates number of cells tracked among 3 biological replicates (ANOVA + Tukey test). **(D)** Representative electron micrographs of the indicated *V. parahaemolyticus* strains stained with uranyl acetate. **(E)** Quantification of flagellation pattern in the populations of the indicated *V. parahaemolyticus* strains. **(F)** Flagellum stain of indicated *P. putida* strains with Alexa Fluor 488-C5-maleimide and **(G)** quantification of the corresponding flagellation in the population. N indicates the number of cells counted among 3 biological replicates. For *S. putrefaciens*, see **Supplementary Figure 3**.

Based on this set of experiments, we concluded that FipA and FlhF are essential for normal formation of polar flagella.

### FipA and HubP ensure proper localization of FlhF to the cell pole

As our previous results showed an interaction between FipA and FlhF, we analyzed if the intracellular localization of FlhF was influenced by FipA. In this regard, we used functional translational fusions of FlhF to fluorescent proteins expressed from its native site on the chromosome in *V. parahaemolyticus* (**Supplementary Figs. S4 and S5**; Arroyo-Pérez and Ringgaard 2021).

We observed that FlhF localized either diffusely in the cytoplasm or to the cell pole as previously reported (Arroyo-Pérez and Ringgaard 2021). However, a significant delocalization of FlhF from the cell pole occurred in the absence of FipA (**Fig. 3A-C; Supplementary Fig. 6**). Particularly, FlhF was diffusely localized in ∼37% of cells or localized to the cell pole in a uni- and bi-polar manner in ∼45% and ∼19%, respectively, compared to wild-type cells (**Fig. 3B**). Absence of FipA significantly reduced localization of FlhF to the cell pole with a concomitant increase in diffusely localized FlhF (70%; **Fig. 3A-C, Supplementary Fig. 6**). Furthermore, the foci of FlhF at the cell pole in the absence of FipA were significantly dimmer compared to wild-type FlhF foci (**Fig. 3D**), indicating that the amount of FlhF localized to the cell pole is lower in the absence of FipA. Importantly, even though FlhF was still able to localize to the cell pole in a certain number of cells lacking FipA, no flagellum was formed in this background (**Fig. 2D-E**), thus indicating that FipA not only is required for proper polar recruitment of FlhF, but also for the stimulation of flagellum formation.

**Figure 3.**
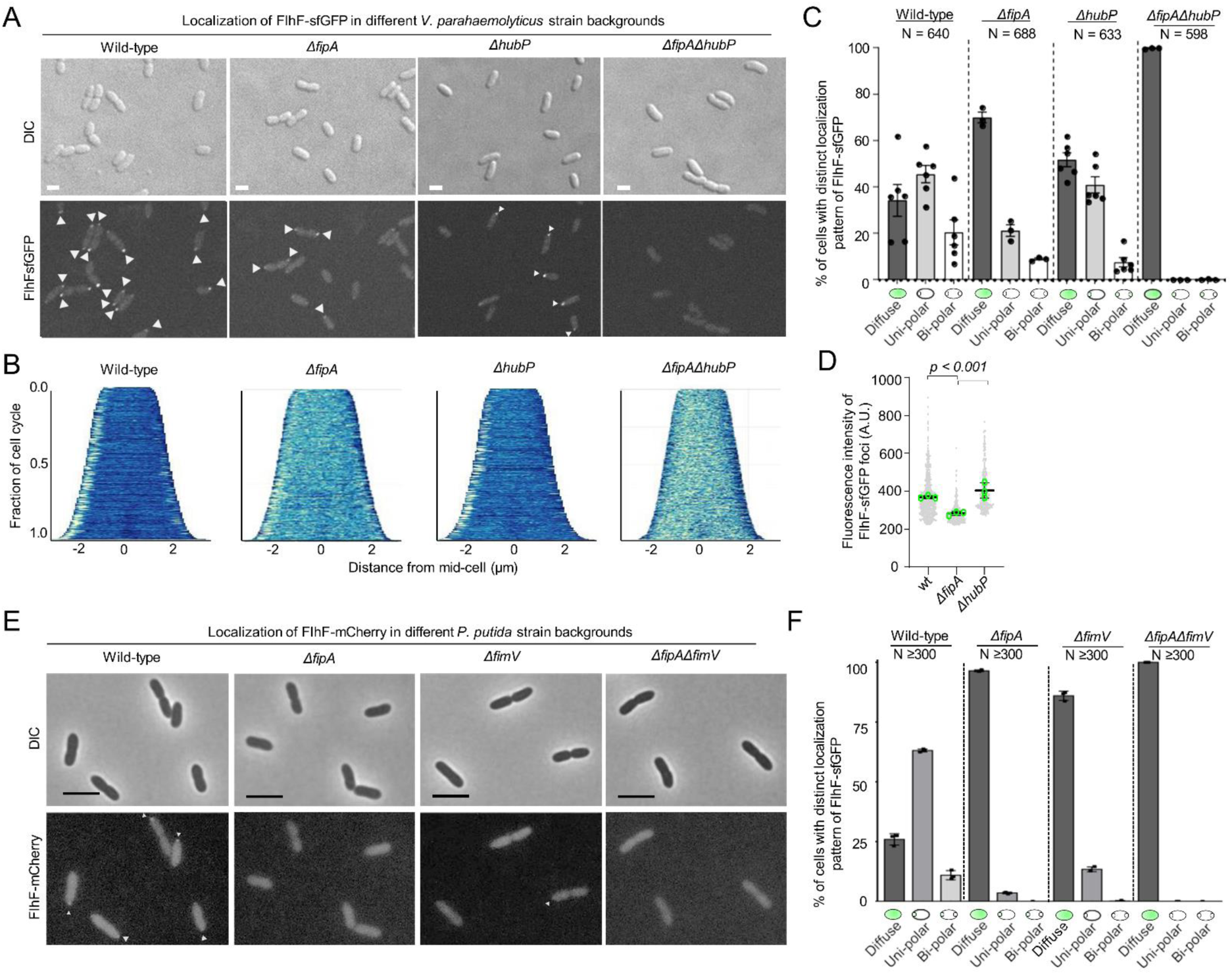
Localization of FlhF depends on FipA and HubP. **(A)** Representative micrographs of the indicated strains of *V. parahaemoloyticus* expressing FlhF-sfGFP from its native promoter. Upper panel shows the DIC image, the lower panel the corresponding fluorescence image. Fluorescent foci are highlighted by white arrows. Scale bar = 2 μm. For an enlargment see **Supplementary Fig. 6. (B)** Demographs displaying FlhF-sfGFP fluorescence intensity along the cell length within the experiments shown in (A). **(C)** Quantification of localization patterns and **(D)** foci fluorescence intensity of the fluorescence microscopy experiment presented in (A). The data was combined from the given number (N) of cells combined from three biological replicates. **(E)** Representative micrographs of the indicated *P. putida* strains expressing FlhF-sfGFP from its native promoter. The upper panels show the DIC and the lower panels the corresponding fluorescence images. Fluorescent foci are marked by small white arrows. The scale bar equals 5 µm. The low intensity of the foci did not allow a quantitative analysis of foci intensities or the generation of demographs. For an enlargement of the micrographs see **Supplementary Fig. 7. (F)** Quantification of FlhF-sfGFP localization patterns in the corresponding strains of *P. putida* from the experiments shown in **(E)**. Corresponding data on the localization of FlhF in *S. putrefaciens* is displayed in **Supplementary Fig. 8**.

Given the role of HubP in cell pole organization and its reported interaction with FlhF (Yamaichi et al, 2012), we also analyzed the localization of FlhF in the absence of HubP. As for FipA mutants, recruitment of FlhF to the cell pole was reduced in the absence of HubP (**Fig. 3A-C, Supplementary Fig. 6**). Strikingly, in the double deletion strain *ΔfipA ΔhubP*, FlhF did not localize as foci at the cell pole at all but was instead localized diffusely in the cytoplasm in 100% of cells (**Fig. 3A-C, Supplementary Fig. 6**). Immunoblot analysis showed that the difference in localization of FlhF-sfGFP was not due to differences in expression levels or protein stability (**Supplementary Fig. S5**).

Altogether, these data suggest that both FipA and HubP work together to promote normal polar localization of FlhF.

### The transmembrane and conserved cytoplasmic domains of FipA is required for its function in regulating flagellum formation and proper FlhF localization

In order to identify the regions of FipA mediating its role in flagellum regulation, we next analyzed the N-terminal transmembrane domain for FipA membrane anchoring and its function in mediating proper flagellum production. When expressed in *E. coli*, a C-terminally GFP-tagged version of FipA (*Vp*FipA-sfGFP) showed a clear membrane localization, while a FipA variant deleted for the predicted transmembrane (TM) domain between residues 7-27 (FipAΔTM), was diffusely localized in the cytoplasm (**Fig. 4A**). These observations show that FipA is indeed a membrane protein anchored by the predicted transmembrane domain. To then study the function of FipA membrane localization, the native *fipA* locus was replaced by a gene encoding a FipA variant lacking the N-terminal TM domain, and the resulting strain (*fipA ΔTM*) was analyzed for flagellum production. The *V. parahaemolyticus* strain carrying this mutation did not produce any flagella at all (**Fig. 4B**), as did the Δ*fipA* strain (**Fig. 2D**).

**Figure 4.**
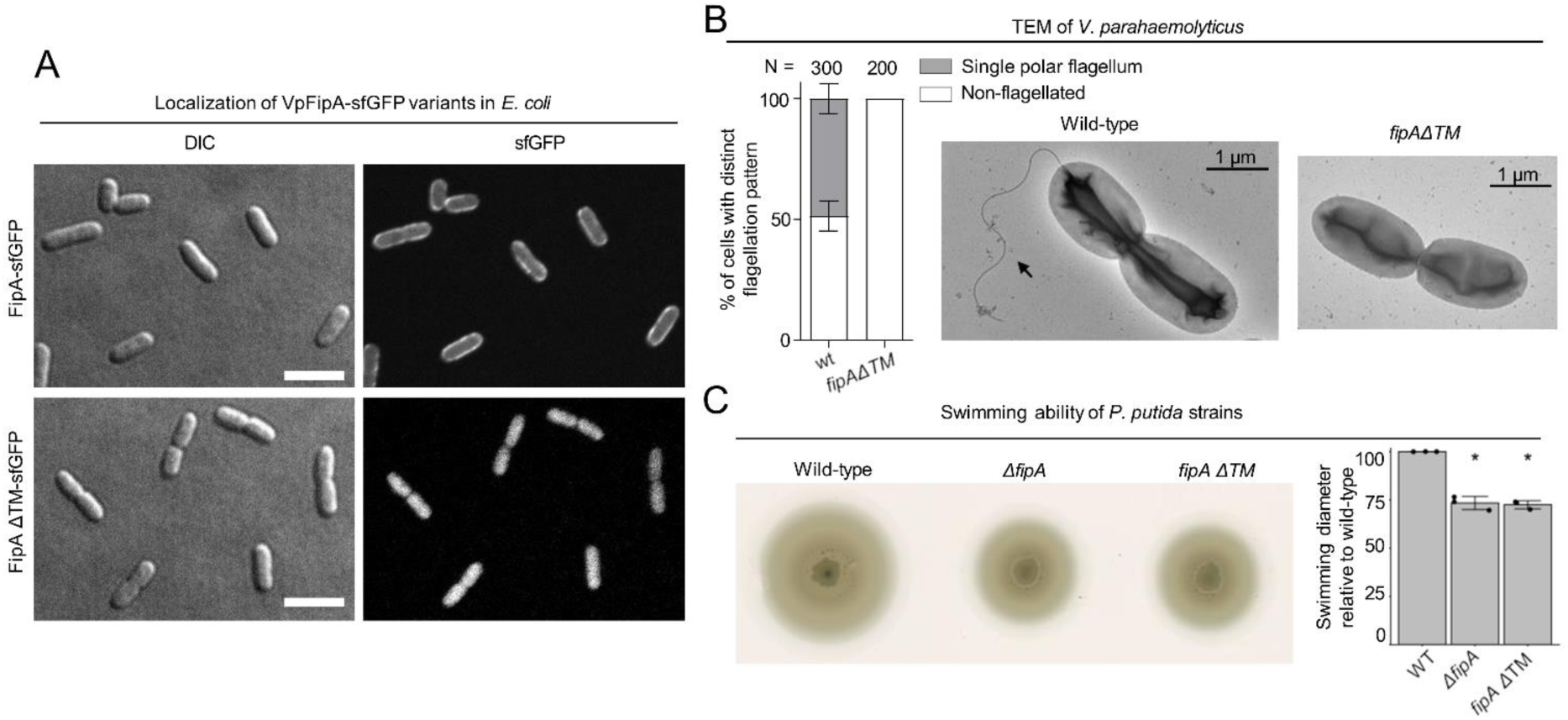
Activity of FipA depends on FlhF and on its transmembrane domain. **(A)** Micrographs of *E. coli* cells expressing FipA-sfGFP from *V. parahaemolyticus*, and a truncated version lacking the transmembrane domain (ΔTM). The left panels display the DIC and the right panels the corresponding fluorescence images. The scale bar equals 5 μm. **(B)** Electron micrographs of *V. parahaemolyticus* wild-type and mutant cells lacking the transmembrane domain of *fipA*, respectively. The corresponding quantification of the flagellation pattern is shown to the left of the micrographs. Note that the data for the wild-type cells is the same as in Fig. 2D, E. **(C)** Spreading behavior of the indicated *P. putida* strains (left) with the corresponding quantification (right). Loss of the FipA TM region phenocopies a complete *fipA* deletion.

After validating an essential role for membrane anchoring of FipA, we analyzed in more detail the cytoplasmic DUF2802 domain. By aligning various FipA homologs, we identified a motif of conserved amino acids, and three of them (G110, E126 and L129 of *Vp*FipA) were 100% conserved among FipA homologues (**Fig. 5A**). Therefore, we chose these amino acids for mutagenesis. Alanine substitutions of residues G110 and L129 abolished the interaction between *Vp*FipA and *Vp*FlhF in BACTH analysis, while the E126A substitution did not affect the interaction (**Fig. 5B**). All these variants were, however, still able to self-interact with wild-type *Vp*FipA (**Fig. 5B**). Altogether, these results suggest that conserved residues in the DUF2802 domain support the interaction between FipA and FlhF, and that the self-interaction is mediated by a different region.

**Figure 5.**
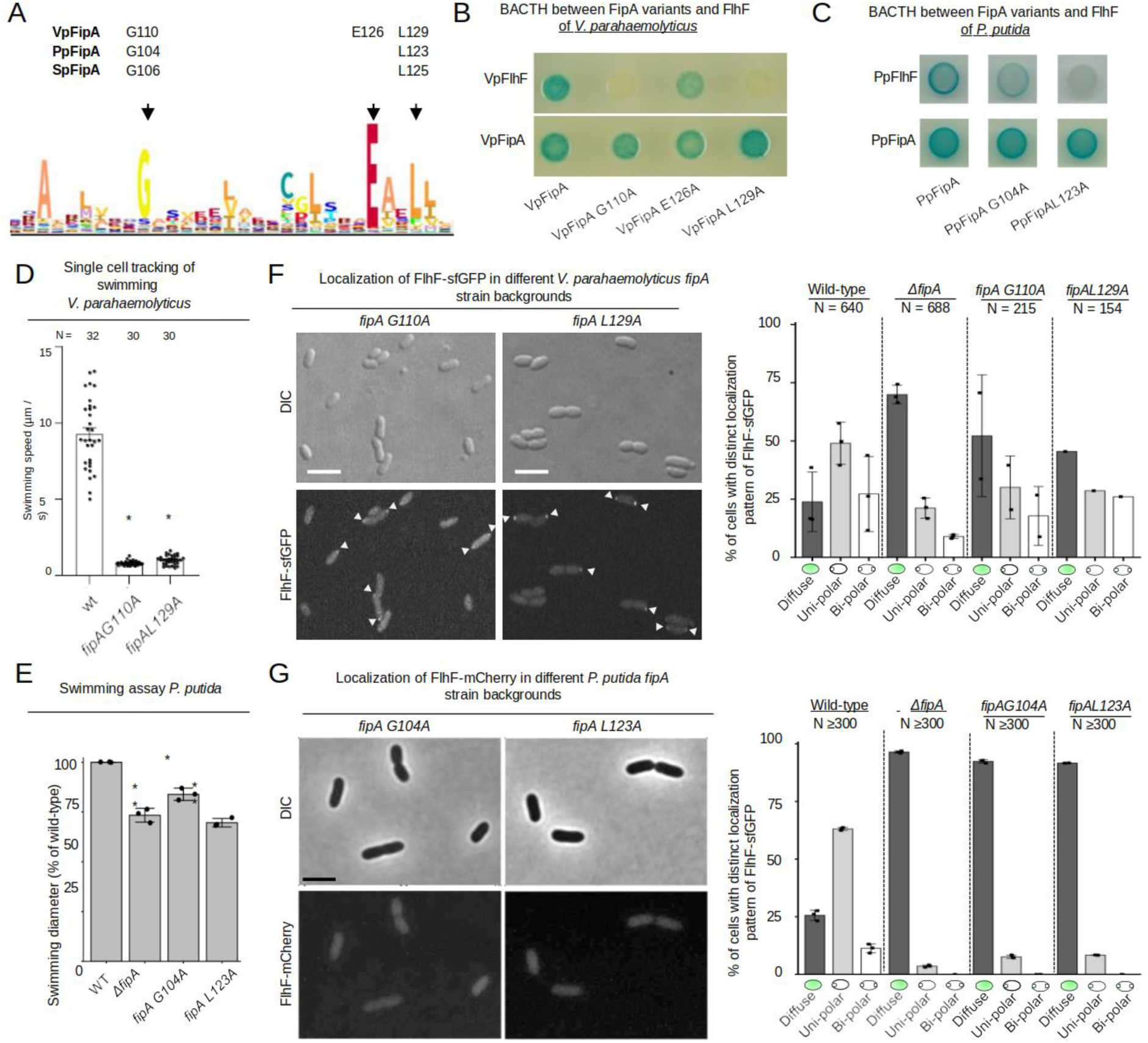
Conserved residues in the domain of unknown function of FipA are essential for interaction with FlhF. **(A)** Weight-based consensus sequence of the conserved region of DUF2802 as obtained from 481 species. The residues targeted in the FipA orthologs of *V. parahaemolyticus*, *P. putida* and *S. putrefaciens* are indicated along with their appropriate residue position. **(B, C)** Bacterial two-hybrid assay of FipA variants of *V. parahaemolyticus* **(B)** or *P. putida* **(C)** with an alanine substitution in the conserved residues indicated in (A). The constructs were tested for self-interaction and interaction with FlhF. *In vivo* interaction of the fusions in *E. coli* is indicated by blue coloration of the colonies. **(D)** Quantification from single-cell tracking of swimming *V. parahaemolyticus* cells expressing FipA bearing the indicated substitution in the DUF2802 domain (see Figure 2B for wild-type behavior). Asterisks indicate a p-value <0.05 (ANOVA + Tukey test) **(F)** Localization of *Vp*FlhF in the absence of FipA or in cells with substitutions in the DUF2802 domain. Left: Micrographs showing the localization of FipA-sfGFP in the indicated strains; the upper panels display the DIC and the lower panels the corresponding fluorescence images (for an enlargement see **Supplementary Fig. 9A**). The scale bar equals 5 µm. Right: the corresponding quantification of the FlhF-sfGFP patterns in the indicated strains. **(E)** Soft-agar spreading assays of *P. putida* wild-type and indicated mutant strains, asterisks display a p-value of 0.05 (*) or 0.01 (**) (ANOVA). **(G)** Localization of *P. putida* FlhF in strains bearing substitutions in the DUF2802 interaction site. Left: micrographs displaying the localization of FlhF-mCherry in the indicated strains. Upper panels show the DIC and lower panels the corresponding fluorescence images. The scale bar equals 5 µm (for an enlargement see **Supplementary Fig. 9B**). Right: Corresponding quantification of the FlhF localization pattern in the indicated strains. Data for *S. putrefaciens* is displayed in **Supplementary Fig. 10**.

### Interaction between FipA and FlhF is required for FipA function on regulating flagellum formation and FlhF localization

We proceeded to evaluate the role of these residues *in vivo*. After introducing the mutations in the *fipA* gene in its native loci, the effects on motility were almost indistinguishable from a Δ*fipA* mutation. In *V. parahaemolyticus,* this resulted in completely non-motile cells in planktonic cultures (**Fig. 5D**).

The localization of FlhF was also affected by the residue substitutions in *fipA.* Cells of *V. parahaemolyticus* expressing FlhF-sfGFP natively had reduced polar localization when FipA was substituted with either G110A or L129A (**Fig. 5F; Supplementary Fig. 9**), with almost 50% of cells presenting only diffuse FlhF-sfGFP signal. Thus, it seems that the DUF2802 domain of FipA is responsible for the effect of FipA on motility, and that its effects occur primarily through the interaction with FlhF. Furthermore, the interruption of FipA-FlhF impedes the recruitment of FlhF to the cell pole.

### Cell cycle-dependent polar localization of FipA

Given FipA’s function in regulating correct recruitment of FlhF to the cell pole, we next analyzed the intracellular localization of FipA. To this end, a hybrid gene bearing a fusion of *fipA* to *sfgfp* was used to replace native *fipA*, resulting in stable and functional production of FipA C-terminally tagged with sfGFP (**Supplementary Fig. 11**). In wild-type strains, FipA localization remained diffuse in half of the population (**Fig. 6A, Supplementary Fig. 12**). In the other half of the population, FipA formed distinct foci at the cell pole, either uni- or bi-polarly (**Figs. 6A, Supplementary Fig. 12**). No delocalized foci were ever observed. The FipA localization behavior was explored further by following the protein through the cell cycle. We observed that FipA foci disappear frequently (**Fig. 6C; 14’**). A new focus appears at the opposite pole, sometimes before cell division (**Fig. 6C; 56’**), sometimes after (**Fig. 6C; 42’**). On rare cases, the first focus persists until after the second focus appears, resulting in bipolar foci (**Fig. 6C; 70’**). These results are consistent with a protein that is recruited to the cell pole right before the start of the flagellum assembly after cell division, albeit the timing may vary between different species.

**Figure 6.**
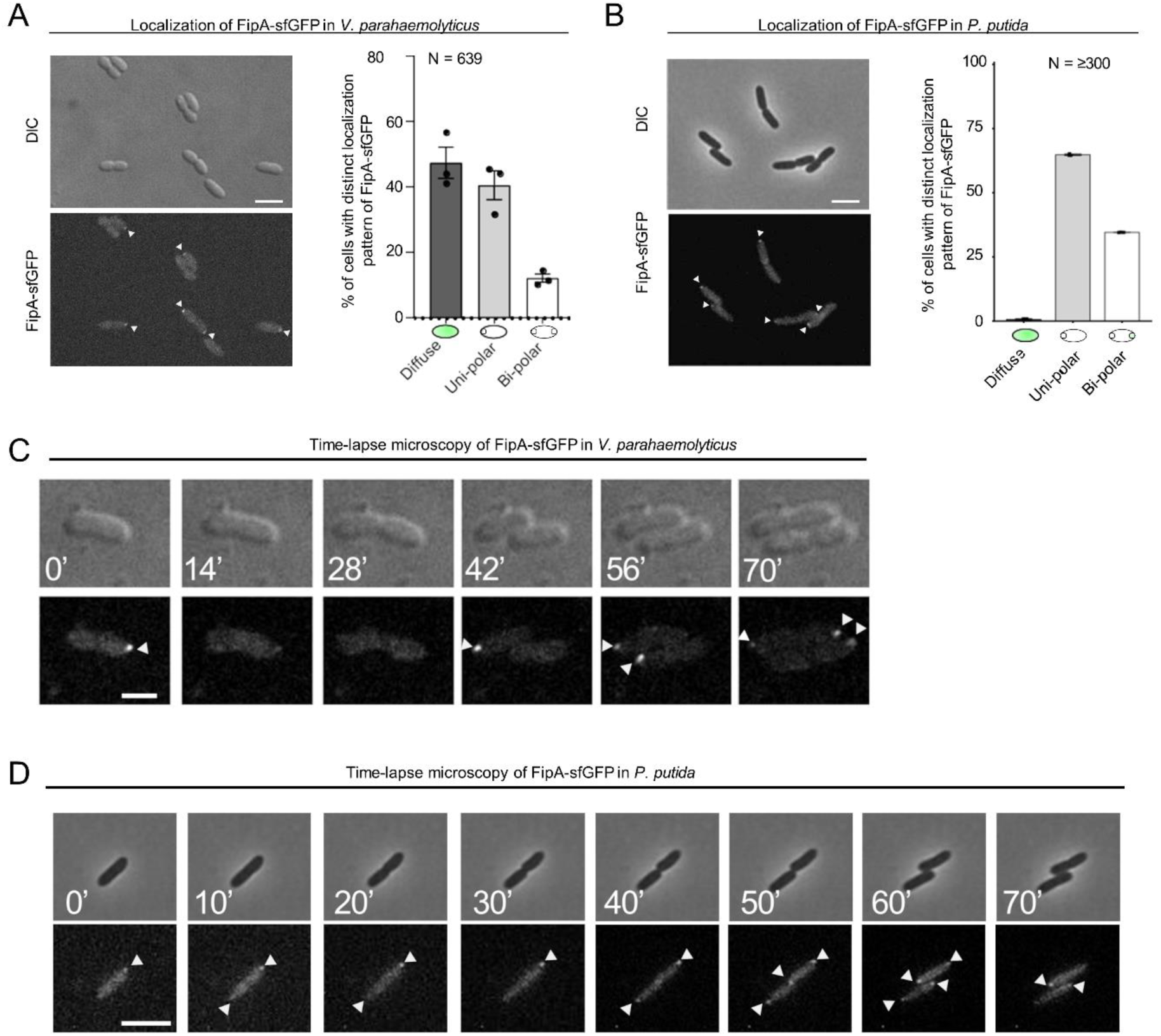
The localization pattern of FipA. **(A, B)** Localization pattern of fluorescently labeled FipA in *V. parahaemolyticus* and *P. putida*. **(A)** Representative micrographs of *V. parahaemolyticus* expressing FipA-sfGFP from its native promoter. Scale bar = 2 μm. The upper panel shows the DIC and the lower panel the corresponding fluorescence channel. To the right the localization was quantified accordingly. **(B)** The same analysis for *P. putida*. **(C, D)** Time lapse analysis of FipA-sfGFP localization over a cell cycle in *V. parahaemolyticus* **(C)** and *P. putida* **(D)**. The numbers in the upper DIC micrographs show the minutes after start of the experiment. The scale bars equal 1 µm (C) and 5 µm (D).

### Polar localization of FipA depends on the FipA-FlhF interaction

Finally, we determined whether FlhF plays a role in FipA localization. To this end, FlhF was deleted in the strains that are expressing FipA-sfGFP from its native promoter. Almost no FipA foci were detected in this background for both *V. parahaemolyticus* (**Fig. 7A-C, Supplementary Fig. 13**) even though the protein levels of FipA were comparable to that in the wild type (**Supplementary Fig. 11**). Furthermore, the FipA variants that do not interact with FlhF (**Fig. 5B**) were also labeled in the native *fipA* locus with sfGFP. Both *Vp*FipA G110A and *Vp*FipA L129A were incapable of forming polar foci (**Fig. 7B, C; Supplementary Fig. 13**).

**Figure 7:**
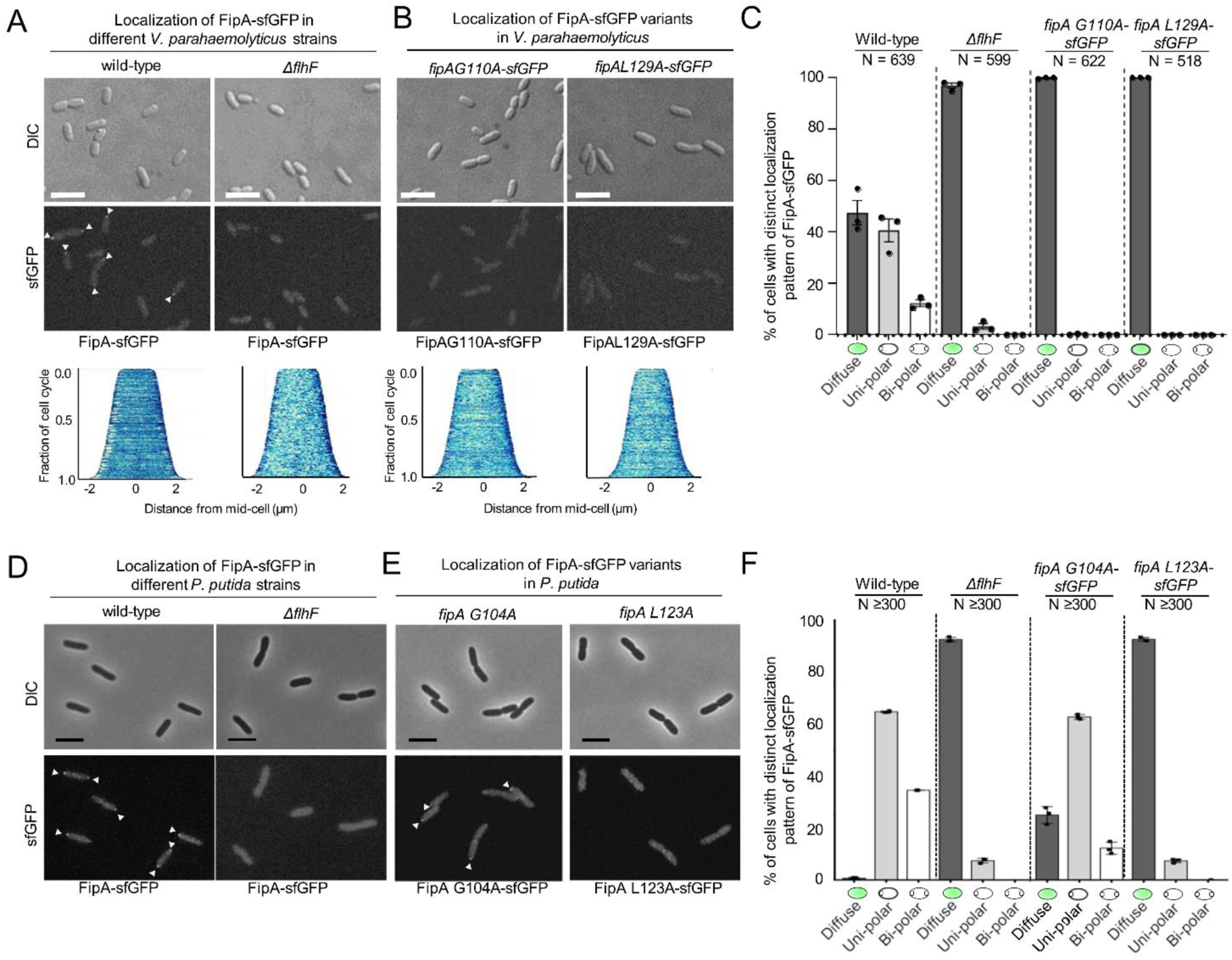
Normal localization of FipA depends on interaction with FlhF. **(A, B)** Localization pattern of *V. parahaemolyticus* FipA-sfGFP in the indicated wild-type and mutant strains. The upper panels display the DIC micrographs, the middle panel the corresponding fluorescence imaging (scale bar equals 5 µm), and the lower panel the corresponding demograph showing the fluorescence of FipA-sfGFP along the cell length. **(C)** Quantification of the cell localization pattern from the experiment shown in (A, B) as combined from three biological replicates. **(D, E, F)** The same analysis for the corresponding *P. putida* strains as indicated. The scale bar equals 5 µm. The data for *S. putrefaciens* is displayed in **Supplementary Figure 15**.

Altogether, these results suggest that, in *V. parahaemolyticus*, a direct interaction with FlhF, mediated by the residues in the DUF2802, is responsible for recruiting FipA to the pole.

### Functional conservation of FipA in *S. putrefaciens* and *P. putida*

Our FipA homology analyses (**Fig. 1F**) indicated that the protein is widely conserved in species also possessing FlhF and FlhG. Accordingly, as in *V. parahaemolyticus*, FipA interacted with FlhF in the polarly flagellated gammaproteobacteria *S. putrefaciens*, which is monopolarly flagellated and *P. putida*, which is lophotrichously flagellated (**Supplementary Fig. 2**). We therefore asked to what extent the role of FipA is conserved for flagellation in these two species. To avoid interference with the secondary non-polar flagellar system of *S. putrefaciens*, we used a strain that is unable to form the secondary flagella (*ΔflaAB_2_*). Determination of the flagellation state was carried out by fluorescence labeling of the flagellar filament(s) in both species.

#### Spreading

As observed for *V. parahaemolyticus*, loss of FlhF and FipA negatively affected flagella-mediated spreading in soft agar and flagellation of *P. putida* and *S. putrefaciens*. However, for both species, the phenotype was not as severe as in *V. parahaemolyticus*: We observed about 40% and 50% in soft-agar spreading of *ΔflhF* mutants and about 25% and 90% spreading of *ΔfipA* mutants in *P. putida* and *S. putrefaciens*, respectively (**Fig. 2A, 2C**; **Supplementary Fig. 3**). Accordingly, the mutations of *flhF* or *fipA* also negatively affected flagella number and localization in both species, with the exception of a *fipA* deletion in *S. putrefaciens*, where a dislocalization could not be observed (**Fig. 2F, G; Supplementary Fig. S3**). Altogether, these results indicate that FipA is necessary for normal flagellation and swimming motility in all three model species, albeit at a different extent reaching from almost complete loss of swimming in *V. parahaemolyticus* to only a small effect in *S. putrefaciens*.

#### Localization of FlhF

We then determined the localization of FlhF in mutants of FipA and HubP or FimV (the HubP homolog in *P. putida*) by using translational fusions of FlhF to fluorescent proteins expressed from the native site on the chromosome in *P. putida* and *S. putrefaciens* (**Supplementary Figs. 4 and 5**; Rossmann et al. 2015). As observed for *V. parahaemolyticus*, FipA and FimV were required for normal FlhF-mCherry localization in *P. putida* (**Fig. 3E, F; Supplementary Fig. 7**), and FlhF was completely delocalized in the double mutant *ΔfipA ΔfimV*. In contrast, FlhF localization was not affected in a *ΔhubP* background in *S. putrefaciens* (**Supplementary Fig. S8**; Rossmann et al, 2015). However, in a *S. putrefaciens* mutant lacking *fipA*, monopolar localization of FlhF-mVenus decreased significantly (**Supplementary Fig. S8**). When both *hubP* and *fipA* were deleted in this species, only weak polar localization remained in a minority of cells. Frequently, polar fluorescence was completely lost and FlhF-mVenus was distributed in the cytoplasm. The differences in localization were not due to differences in abundance of FlhF (**Supplementary Fig. 5**)

The results showed that the function of FipA for flagellation and FlhF localization is generally conserved within the three different species. However, the polar landmark HubP appears to be involved in localizing FlhF, in particular in *S. putrefaciens*, where both HubP and FipA need to be deleted to delocalize FlhF from the cell pole.

#### FipA localization pattern

As the next step, we determined the conservation of FipA domains’ function and localization. We found that, as observed in *V. parahaemolyticus*, loss of the FipA transmembrane region (*fipA ΔTM*) phenocopied a *ΔfipA* mutant in *P. putida* and *S. putrefaciens* (**Fig. 4C**, **Supplementary Figs. S3A)**. Furthermore, substitution of the conserved residues within the FipA DUF domain (G104A and L123A in *P. putida* FipA; G106A and L118A in *S. putrefaciens* FipA) decreased or abolished the interaction with FlhF but not FipA self-interaction (**Fig. 5C, G; Supplementary Fig. S10**).

In *P. putida*, similar to *V. parahaemolyticus*, FipA formed distinct foci at the cell pole, either uni- or bi-polarly (**Figs. 6A, B; Supplementary Fig. 12**). A remarkable difference between *P. putida* and *V. parahaemolyticus* is that in *P. putida,* FipA occurred as foci in virtually all cells (**Fig. 6B**), whereas in *V. parahaemolyticus*, FipA remained diffuse in half of the population (**Fig. 6A**). The ratio of bi-polar to uni-polar foci was also greater in *P. putida* (∼1:2) than in *V. parahaemolyticus* (∼1:5). Furthermore, in *P. putida*, the FipA foci were more stable, and foci at both poles often persisted until cell division. In fact, appearance of the second focus at the new pole frequently occurred right after cell division (**Fig. 6D; 50’**), explaining the greater bi-polar to uni-polar ratio of FipA foci in this organism.

Finally, in *V. parahaemolyticus*, the localization but not the stability of FipA is dependent on the interaction with FlhF (**Fig. 7, Supplementary Fig. 11**). This was similarly observed for *P. putida*, as the FipA L123A variant, which is unable to interact with FlhF (**Fig. S2A**), was also mostly distributed in the cytoplasm, whereas the variant FipA G104A did exhibit some polar localization, although reduced (**Fig. 7E, F; Supplementary Fig. 14**). This is the variant that did show some interaction with *Pp*FlhF (**Fig. 5C**). These results suggest that, in *V. parahaemolyticus* and *P. putida*, a direct interaction with FlhF, mediated by the residues in the DUF2802, is responsible for recruiting FipA to the pole.

Surprisingly, in *S. putrefaciens* mutants deleted in *flhF*, FipA displayed only a slight difference in polar localization (**Supplementary Fig. 15**). However, a *Sp*FipA G106A substitution reduced the protein’s localization (to ∼50% cells with foci). It is likely that in *S. putrefaciens*, HubP is (co-)mediating the localization of FipA, as in the absence of HubP, FipA-sfGFP localization at the cell pole drastically decreases (**Supplementary Fig. 15**). As observed for *V. parahaemolyticus* and *P. putida*, the stability of FipA was not dependent on the presence of FlhF or FipA in *S. putrefaciens* **(Supplementary Fig. S5).**

## Discussion

In bacteria, numerous processes are localized to specific cellular compartments. This is particularly evident in polarly flagellated bacteria, where the intricate flagellar multicomplex needs to be synthesized in a spatiotemporally regulated fashion. Flagellar synthesis is initiated by the assembly of the first flagellar building blocks within the cytoplasmic membrane, which in polar flagellates is targeted to the designated cell pole. In a wide range of bacterial species, the SRP-type GTPase FlhF and its antagonistic counterpart, the MinD-like ATPase FlhG, regulate flagellar positioning and number (Kazmierczak and Hendrixson 2013, Schuhmacher, Thormann, and Bange 2015). In particular, FlhF has been shown to function as a positive regulator and a major localization factor for the initial flagellar building blocks of polar flagella. Upon production, GTP-bound dimeric FlhF localizes to the cell pole, but the mechanism by which the protein assumes its designated position at the cell pole remained elusive. One candidate protein, the polar landmark HubP (FimV in *Pseudomonas* sp.) had been identified earlier and was shown to interact with FlhF in *V. parahaemolyticus* (Yamaichi et al, 2012; Dornes et al, 2024). Accordingly, polar localization of FlhF is decreased in mutants lacking *hubP* (this study; Rossmann, Brenzinger et al, 2015), however, in a number of cells FlhF localized normally and the cells showed normal flagellation. This was indicative that HubP does play a role in functionally recruiting FlhF, but that another factor was still missing. In this study we identified the protein FipA as a second polarity factor for FlhF.

FipA consists of an N-terminal transmembrane domain and a cytoplasmic region with a conserved domain of unknown function (DUF2802). The corresponding gene is located immediately downstream of the motility operon that includes *flhF* and *flhG*. Notably, FipA is highly conserved among bacteria that use the FlhF/FlhG system to position the flagella, strongly suggesting that FipA has evolved together with FlhF and FlhG to regulate the flagellation pattern. In this study, we therefore investigated three distantly related γ-proteobacteria including one lophotrichous species, *V. parahaemolyticus*, *P. putida* and *S. putrefaciens*, in more detail. We found that FipA is conserved as a regulatory factor of FlhF and that its functions rely on similar features in all three species.

FipA is essential for normal flagellum synthesis, however, the severity of a *fipA* deletion differed among the three species studied. In *V. parahaemolyticus*, a *fipA* deletion phenocopies the loss of *flhF*, and cells no longer synthesize flagella. Notably, in the absence of FipA in *V. parahaemolyticus,* FlhF is still localized at the pole in about a third of the cells. Despite this, these cells were still unable to form flagella, strongly suggesting that occurrence of FlhF at the pole alone is not sufficient to trigger flagellum synthesis in the absence of FipA.

This strict requirement for FlhF has been reported in the closely related *V. alginolyticus* (Kusumoto et al, 2009). On the other hand, in *V. cholerae,* the requirement for FlhF on flagellation is less strict (Green et al, 2009), and in *P. putida* and *S. putrefaciens* even less so: mutants deleted in *flhF* or *fipA* of both species can generally still produce flagella and swim (this work; Rossmann et al, 2015). In *P. putida,* a Δ*fipA* mutant has a phenotype reminiscent of a Δ*flhF* mutant as both drastically decrease the number of flagella, which may also be delocalized in *flhF* mutants (Pandza et al, 2003). In *S. putrefaciens*, deletion of *fipA* decreases the number of flagella to a similar extent as a *flhF* mutation, however, while in the latter case the flagella are frequently delocalized, the flagella remain at a polar postion, when *fipA* is missing. Taken together, these results strongly suggest that FipA does not act as a general polarity factor for the full flagellar machinery, but rather it stimulates the activity of FlhF to initiate polar flagellation.

### FipA and HubP affect polarity of FlhF through different mechanisms

In all three species, FipA directly interacts with FlhF as demonstrated by reciprocal co-immuno precipitations and by bacterial two-hybrid assays. These experiments suggest that FipA is able to dimerize (or oligomerize) and that the FlhF-FipA interaction is dependent on conserved residues in the DUF2802 domain. Furthermore, these residues are not required for the ability of FipA to interact with itself, suggesting that the two processes occur independently and at different protein interfaces. Microscopic and physiological assays showed that direct interaction between FlhF and FipA is required for FlhF targeting and function. In addition, a FipA mutant lacking its N-terminal transmembrane domain was non-functional, indicating that interaction of FlhF and FipA has to take place at the membrane.

In the absence of FipA, polar localization of FlhF was significantly decreased in all three species and almost completely diminished when *hubP* or *fimV* was deleted together with *fipA*. Removing HubP/FimV alone had a similar effect as removing FipA on the localization of FlhF, although to a lesser extent, as more cells still had polar FlhF in the Δ*hubP/fimV* mutant than in the Δ*fipA* mutant. Furthermore, the unipolar to bipolar ratio of FlhF foci increased in the Δ*hubP* mutant compared to the wild-type or the Δ*fipA* background. While the deletion of *fipA* increases the number of cells with diffuse localization, it does not affect the time frame in the cell cycle at which FlhF shifts from unipolar to bipolar in *V. parahaemolyticus*. In contrast, deleting *hubP* delays the time frame in which FlhF assumes a bipolar position. This indicates that HubP and FipA represent two different pathways that stimulate polar localization of FlhF, and that both are required to bring sufficient (active) FlhF molecules to trigger MS-ring formation and start flagellum assembly.

### Localization of FipA

A notable feature of FipA is its dynamic localization pattern within the cell over the cell cycle in all three species studied, the nature of which is unclear so far. Foci of FipA may appear mono- or bipolarly and frequently, but not necessarily, disappear once the flagellar apparatus, or the flagellar bundle in the case of *P. putida*, is established. It remains to be shown whether the FipA protein is actively moving from one pole to another or if newly formed FipA is recruited to the opposite pole while being degraded at the old position. Notably, in *V. parahaemolyticus* and *P. putida* FipA does not localize polarly in the absence of interaction with FlhF, suggesting that both proteins are recruited as a complex.

In contrast, in *S. putrefaciens,* FipA still localizes to the cell pole also in the absence of FlhF as long as the polar landmark HubP is present. This is indicating a more important role for HubP in this species. Accordingly, it has been shown recently that in *S. putrefaciens* HubP directly interacts with HubP via its NG domain to recruit the initial flagellar building blocks (Dornes et al, 2024. However, also in *S. putrefaciens* active FlhF localizes to the cell pole and flagella are formed in the absence of HubP. The nature of the polar marker that recruits or guides FipA or a FipA/FlhF to the designated pole is still elusive. The discovery of FipA provides a new point where spatiotemporal organization is coordinated. However, it remains to be explored if the complex is anchored through yet another protein, such as the TonB_m_-PocA-PocB complex in *P. putida* (Ainsaar et al, 2019) or through an intrinsic feature of the cell envelope or cytoplasm during cell division.

### How does FipA upgrade the current model of polar flagellar synthesis?

Based on the findings in different species, our current model of FlhF/FlhG-mediated polar flagellar synthesis predicts, that upon production, GTP-bound dimeric FlhF is localizing or being recruited to the cell pole to where it recruits the first flagellar components to initiate the assembly. The monomeric form of the ATPase FlhG, the FlhF antagonist, is binding to the flagellar C-ring building block FliM and is transferred to the nascent flagellar structure, where it is released and dimerizes upon ATP binding (Schuhmacher et al, 2015; Blagotinsek et al, 2020). The ATP-bound FlhG dimer can associate with the membrane and interact with the GTP-bound FlhF dimer, thereby stimulating its GTPase activity. This leads to monomerization of FlhF and loss of polar localization (reviewed in Kojima, Terashima, and Homma 2020; Schuhmacher et al., 2015). Dimeric FlhG does also interact with the master regulator of flagella synthesis, FlrA (or FleQ in *Pseudomonas*) and prevents the synthesis of further early flagellar building blocks (Dasgupta & Ramphal, 2001; Blagotinsek et al, 2020; Chanchal et al, 2021). By this, FlhG links flagella synthesis with transcription regulation and effectively restricts to number of polar flagella that are formed. Based on this model, FipA may recruit FlhF to the membrane and stimulate or stabilize GTP binding and dimerization of FlhF to promote polar localization and initiation of flagella synthesis. Thus, the role of FipA is to shift the equilibrium to the active state of FlhF in order to start assembly of the MS ring. Accordingly, current studies address the structural basis of the FipA-FlhF interaction, the potential effect of FipA on GTP-binding and dimerization of FlhF, and the role of the polar marker HubP in this process. In addition, *fipA* expression is independent of the rest of the flagellar genes in other species of *Vibrio* (Moisi et al. 2009; Petersen et al. 2021) and *S. putrefaciens* (Schwan et al, 2022). Thus, *fipA* transcription could provide another regulatory point to lead to the synthesis of a flagellum. Thus, the discovery of FipA opens several novel open questions in the field of flagellum regulation.

## Materials and methods

### Growth conditions and media

In all experiments *V. parahaemolyticus*, and *E*. *coli* were grown in LB medium or on LB agar plates at 37°C containing antibiotics in the following concentrations: 50 μg/mL kanamycin, 100 μg/mL ampicillin and 20 μg/mL chloramphenicol for *E*. *coli* and 5 μg/mL for *V. parahaemolyticus*. *S. putrefaciens* CN-32 and *P. putida* KT 2440 were grown in LB medium or LB agar plates at 30 °C. When required, the media were supplemented with 50 µg/mL kanamycin, 300 µM 2,6-diaminopimelic acid and/or 10 % (w/v) sucrose.

### Strains and plasmids

The strains and plasmids used in this study are listed in Tables S3 and S4, respectively. Primers used are listed in Table S5. *E. coli* strain SM10λ*pir* was used to transfer DNA into *V. parahaemolyticus* by conjugation (Miller & Mekalanos, 1988). For DNA transfer into *S. putrefaciens* and *P. putida*, *E. coli* WM3064 was used. *E. coli* strains DH5αλ*pir* and SM10λ*pir* were used for cloning. Construction of *V. parahaemolyticus* deletion mutants was performed with standard allele exchange techniques using derivatives of plasmid pDM4 (Donnenberg & Kaper 1991). Chromosomal deletions and integrations in *S. putrefaciens* and *P. putida* were carried out by sequential crossover as previously described (Rossman et al. 2015) using derivatives of plasmid pNPTS138-R6K (Lassak et al. 2010).

### Construction of plasmids

#### Plasmid pSW022

The regions flanking *vp2224* (*fipA*) were cloned with primers VP2224-del-a/ b and VP2224-del-c/ d, using *V. parahaemolyticus* RIMD 2210633 chromosomal DNA as template. The resulting products were fused in a third PCR using primers VP2224-del-a/ VP2224-del-d. The end product was digested with XbaI and ligated in the equivalent site of vector pDM4, resulting in plasmid pSW022. The mutation in *V. parahaemolyticus* was confirmed with a PCR using primers VP2224-del-a/VP2224-check. *Plasmid pPM123:* The regions flanking aa 7-27 of *vp2224* (*fipA*) were cloned with primers VP2224-del-a/ del AA7-27 vp2224-b and del AA7-27 vp2224-c/ VP2224-del-d, using *V. parahaemolyticus* RIMD 2210633 chromosomal DNA as template. The resulting products were fused in a third PCR using primers VP2224-del-a/ VP2224-del-d. The end product was digested with XbaI and ligated in the equivalent site of vector pDM4, resulting in plasmid pPM123. The mutation in *V. parahaemolyticus* was confirmed with a PCR using primers VP2224-del-a/VP2224-check. *Plasmid pPM178:* The gene *vp2224* (*fipA*) was amplified from *V. parahaemolyticus* RIMD 2210633 chromosomal DNA with primers C-term sfGFP-vp2224-a/-b; the downstream region with primers C-term sfGFP-vp2224-e/f; the gene encoding sfGFP with C-term sfGFP-vp2224-c/d from plasmid pJH036. The three products were fused together in another PCR using primers C-term sfGFP-vp2224-a/f. The obtained product, encoding FipA fused in frame to sfGFP via a 5-residue linker, was digested with SpeI and SphI and ligated in vector pDM4 digested with XbaI and SphI, resulting in plasmid pPM178. The mutation in *V. parahaemolyticus* was confirmed with a PCR using primers C-term sfGFP-vp2224-f/VP2224-check. *Plasmid pPM179 & pPM180:* The region upstream of *vp2224* (*fipA*) were amplified as for pSW022. The downstream region was amplified with downstream vp2224-cw/VP2224-del-d from *V. parahaemolyticus* RIMD 2210633. The gene *vp2224* itself was amplified with primers vp2224 cw restore deletion/vp2224 ccw restore deletion, from plasmids pEP005 & pEP006, carrying the mutations G110A and L129A. The products were fused in a third PCR using primers VP2224-del-a/ VP2224-del-d. The end product was digested with XbaI and ligated in the equivalent site of vector pDM4, resulting in plasmids pPM179 & pPM180. The the re-insertion in *V. parahaemolyticus* SW01 was confirmed with a PCR using primers VP2224-del-a/VP2224-check. *Plasmid pPM187 & pPM191:* The insertion of sfGFP at the C-terminus of FipA was cloned with the same strategy for pPM178, but using pPM179 & pPM180 as templates for the *vp2224* sequence, resulting in plasmids pPM191 & pPM187, respectively. *Plasmid pPM146:* The gene *vp2224* (*fipA*) was amplified from *V. parahaemolyticus* RIMD 2210633 chromosomal DNA with primers vp2224-cw-pBAD/vp2224-ccw-pBAD. The product was digested with enzymes XbaI and SphI, and inserted in the corresponding site in plasmid pBAD33. *Plasmid pPM159:* The gene *vp2224* (*fipA*) was amplified from *V. parahaemolyticus* RIMD 2210633 chromosomal DNA with primers C-term sfGFP-vp2224-a/-b, and the gene encoding sfGFP with C-term sfGFP-vp2224-c/sfGFP-1-ccw from plasmid pJH036. The resulting products were fused in another reaction with primers C-term sfGFP-vp2224-a/sfGFP-1-ccw, and this final product was digested with XbaI and cloned in pBAD33. *Plasmid pPM194:* Same as pPM159, but *vp2224 ΔTM* was amplified from pPM123 using primers vp2224 AA1-6/28-end w/o Stop/sfGFP-1-ccw. *Plasmids pSW74 & pSW119:* The region of the gene *vp2224* (*fipA*) encoding the cytoplasmic part (residues 28-163) was amplified with primers tr2224 put18C cw/ pUT18C/pKT25-vp2224-ccw from *V. parahaemolyticus* chromosomal DNA. The product was digested with KpnI and XbaI and ligated in the corresponding site in pKT25, to generate pSW74, and in pUT18C, to generate pSW119. *Plasmid pPM118 & pPM119:* The cytoplasmic part of the gene *vp2224* (*fipA*) was amplified with primers pUT18/pKNT25-tr-vp2224-cw & pUT18/pKNT25-vp2224ccw from *V. parahaemolyticus* chromosomal DNA. The product was digested with KpnI and XbaI and ligated in the corresponding site in pUT18, to generate pPM118, and in pKNT25, to generate pPM119. *Plasmid pPM124 & pPM128:* The gene *vp2234* (*flhF*) was amplified with primers pUT18/pKNT25-vp2234-cw & pUT18/pKNT25-vp2234 -ccw from *V. parahaemolyticus* chromosomal DNA. The product was digested with KpnI and XbaI and ligated in the corresponding site in pUT18, to generate pPM124, and in pKNT25, to generate pPM128. *Plasmid pPM132 & pPM136:* The gene *vp2234* (*flhF*) was amplified with primers pUT18C/pKT25-vp2234-cw & pUT18C/pKT25-vp2234 -ccw from *V. parahaemolyticus* chromosomal DNA. The product was digested with KpnI and XbaI and ligated in the corresponding site in pUT18C, to generate pPM132, and in pKT25, to generate pPM136. *Plasmids pPM160, pPM161 & pPM162:* Site directed mutagenesis was performed on plasmid pSW74 by the QuickChange method (Zheng, Baumann, and Reymond 2004), using primers vp2224-Gly-110Ala-cw/ccw, vp2224-Glu 126Ala-cw/ccw or vp2224-Leu 129Ala-cw/ccw. After digesting the template with DpnI, the products were transformed into *E. coli* and the mutations were confirmed by sequencing. *Plasmid pPM106:* The gene vp2224 (fipA) was amplified with primers pUT18C/pKT25-vp2224-cw & pUT18C/pKT25-vp2224ccw from V. parahaemolyticus chromosomal DNA. The product was digested with KpnI and XbaI and ligated in the corresponding site in pKTop, to generate pPM106. *Plasmid pPM109 & pPM112:* The *phoA-lacZα* fragment of pKTop was amplified with primers vp2224 C-term PhoA-LacZ cw & end -LacZ w/o STOP ccw. The full length *vp2224(fipA)* gene was amplified from plasmid pPM146 using primers LacZ to vp2224 w/o ATG & end vp2224 ccw. The *vp2224* ΔTM allele was amplified from plasmid pPM123 using primers vp2224 AA1-6/28-end w/o Stop & end vp2224 ccw. The PCR products were fused in a second PCR using primers vp2224 C-term PhoA-LacZ cw & end vp2224 ccw. The fusion product was digested with XbaI and HindIII and cloned in the corresponding site of plasmid pBAD33.

#### Plasmids for genetic manipulation of S. putrefaciens and P. putida

The desired DNA fragments were generated by PCR using appropriate primer pairs (see Supplementary Table S5) that in addition create overhangs suitable for subsequent Gibson assembly into EcoV-digested vector pNTPS138-R6K (Gibson et al., 2009; Lassak et al, 2010). If necessary, primer overhangs were also used to generate base substitutions in the gene fragment to be cloned. *Plasmids for Bacterial Two Hybrid (BACTH) analysis* were similarly generated by amplification of the desired DNA-fragments, which were then introduced into the suitable vectors by Gibson assembly.

### Soft-agar swimming assays

*V. parahaemolyticus* soft-agar swimming assays were essentially performed as described in (Ringgaard, et al, 2013) with the following modifications. Late exponential cultures of the required strains were used to prick LB plates with 0.3% agar, and incubated for 30 hours at 30° C. For *S. putrefaciens* and *P. putida* soft-agar swimming assays, 2 µl of an exponentially growing culture of the appropriate strains were spotted onto 0.25 % LB agar plates and incubated at 30 °C (*S. putrefaciens*) or room temperature (*P. putida*) for about 18 hours. Strains to be directly compared were always spotted onto the same plate. For the swimming assays in soft agar, strains of *V. parahaemolyticus* and *S. putrefaciens*, which are lacking lateral flagellin gene(s) (*ΔlafA; ΔflaAB_2_*) were used.

### Bacterial-two-hybrid experiments, BACTH

BTH101 cells were made competent with calcium chloride. 15 μL aliquots were spotted in a 96 well plate, to which 2.5 μL of the corresponding pUT18(C) and pK(N)T25 derivative plasmids were added. After 30 minutes on ice, a heat shock at 42° C was applied for 30 s. The transformed cells were allowed to recover for 1 hour, after which the were grown in selective LB broth for another 3 hours. The resulting cultures were spotted on plates containing kanamycin, ampicillin, IPTG (0.25 mM) and X-Gal (80 μg/mL). The plates were photographed after 48-72 hours at 30° C.

### Video tracking of swimming cells

Video tracking of swimming cells was performed essentially as described previously (Ringgaard, et al, 2013). Swimming cells were recorded using the streaming acquisition function in the Metamorph software and the swimming paths of individual cells were tracked using the MTrackJ plug-in for ImageJ. The swimming speed, displacement, and number of reversals of individual cells were then measured and the average plotted with error bars indicating the standard deviation. Video tracking was performed using a Zeiss Axio Imager M1 fluorescence microscope. Images were collected with a Cascade:1K CCD camera (Photometrics), using a Zeiss αPlan-Fluar 40x/1.45 Oil phase contrast objective.

### Transmission-electron-microscopy (TEM) analysis

Cell cultures grown to an OD600 = 0.5–0.6 were spotted on a plasma-discharged carbon-coated copper grid (Plano, Cat#S162-3) and rinsed with 0.002% uranyl acetate. Afterwards they were rinsed with water and blotted dry with Whatman filter paper. TEM images were obtained with a JEOL JEM-1400 Plus 120 KV transmission electron microscope at 80 kV.

### Flagellum labeling

To fluorescently label flagellar filaments, maleimide-ligates dyes (Alexa Fluor 488 C5 maleimide fluorescent dye; Thermo Fisher Scientific) were coupled to surface-exposed cysteine residues, which were introduced into the flagellins (FlaA and FlaB) of the polar flagellar system in *S. putrefaciens* and *P. putida* as described (Kühn et al. 2017, Hintsche et al. 2017). Images were recorded as described in the microscopy section.

### Fluorescence microscopy

Florescence microscopy on *V. parahaemolyticus* was carried out essentially as previously described (Muraleedharan et al, 2018), using a Nikon eclipse Ti inverted Andor spinning-disc confocal microscope equipped with a 100x lens, an Andor Zyla sCMOS cooled camera, and an Andor FRAPPA system. Fluorescence microscopy on *S. putrefaciens* and *P. putida* was carried out as previously described (Kühn et al. 2017) using a custom microscope set-up (Visitron Systems, Puchheim, Germany) based on a Leica DMI 6000 B inverse microscope (Leica) equipped with a pco.edge sCMOS camera (PCO), a SPECTRA light engine (lumencor), and an HCPL APO 63×/1.4–0.6 objective (Leica) using a custom filter set (T495lpxr, ET525/50m; Chroma Technology) using the VisiView software (Visitron Systems, Puchheim, Germany). Microscopy images were analyzed using ImageJ imaging software (http://rsbweb.nih.gov/ij) and Metamorph Offline (version 7.7.5.0, Molecular Devices). DIC (*Vibrio*) and phase contrast (*Shewanella*, *Pseudomonas*) were used according to the preferred settings for the corresponding species in the labs. Demographs were generated as described by Cameron et al, 2014, and modified in Heering & Ringgaard 2016 and Heering, Alvarado and Ringgaard, 2017.

### Mapping interaction partners using co-immunoaffinity purification and mass spectrometry (co-IP-MS)

For sample preparation of the IP-MS experiments we were using a modified version of the protocol presented by Turriziani et al, 2014. Cells were centrifuged and cell pellets were washed with cold PBS. Cell pellets were resuspended in lysis buffer (50 mM HEPES (pH 7.5), 150 mM NaCl, 0.5 % NP50, 5 mM EDTA, compete mini protease inhibitors (Complete Mini (Roche)). Cell lysis was performed by repetitive sonication. After removing cell debris by centrifugation 10ul GFP-trap Sepharose (Chromotek) slurry was added to the lysate and incubation was carried out for 1.5 hours on a rotating shaker at 4 °C. Then the beads were pelleted, the supernatant removed and the beads washed 4x with 100 mM NH_4_CO_3_ to remove remaining protease inhibitors and detergents. 200 µl elution buffer (1 M urea, 100 mM NH_4_CO_3_, 1 µg Trypsin (Promega)) was added to the beads and incubated at 1000 rpm on a thermomixer at 27°C for 45min. Beads were centrifuged and supernatant was collected. In order to increase peptide recovery 2 washing steps with 100µl elution buffer 2 (1 M urea, 100 mM NH_4_CO_3_, 5 mM Tris(2-caboxyethyl)phosphine)) was performed and the individual bead supernatants were collected into one tube. Tryptic digestion was carried out overnight at 30 °C. After digest alkylation was performed with 10 mM iodoacetamide at 25 °C (in the dark). Then the samples were acidified (1 % trifluoroacetic acid (TFA)) and C18 Microspin column (Harvard Apparatus) was carried out according to the manufactureŕs instruction. The samples were dried and recovered in 0.1 % TFA and applied to liquid chromatography-mass spectrometry (LC-MS) analysis.

LC-MS analysis of digested lysates was performed on a Thermo QExactive Plus mass spectrometer (Thermo Scientific), which was connected to an electrospray ionsource (Thermo Scientific). Peptide separation was carried out using an Ultimate 3000 RSLCnano with Proflow upgrade (Thermo Scientific) equipped with a RP-HPLC column (75 μm x 42 cm) packed in-house with C18 resin (2.4 μm; Dr. Maisch) on an in-house designed column heater. The following separating gradient was used: 98 % solvent A (0.15 % formic acid) and 2 % solvent B (99.85 % acetonitrile, 0.15 % formic acid) to 35 % solvent B over 90 at a flow rate of 300 nl/min. The data acquisition mode was set to obtain one high resolution MS scan at a resolution of 70,000 full width at half maximum (at m/z 200) followed by MS/MS scans of the 10 most intense ions. To increase the efficiency of MS/MS attempts, the charged state screening modus was enabled to exclude unassigned and singly charged ions. The dynamic exclusion duration was set to 30 sec. The ion accumulation time was set to 50 ms (MS) and 50 ms at 17,500 resolution (MS/MS). The automatic gain control (AGC) was set to 3×10^6^ for MS survey scan and 1×10^5^ for MS/MS scans.

MS raw data was then analyzed with MaxQuant (Version 1.6.3.4) (https://www.nature.com/articles/nbt.1511) using a *V.parahaemolyticus* RIMD 2210633 uniprot database (www.uniprot.org). MaxQuant was executed in standard settings with activated “match between runs” option. The search criteria were set as follows: full tryptic specificity was required (cleavage after lysine or arginine residues); two missed cleavages were allowed; carbamidomethylation (C) was set as fixed modification; oxidation (M) and deamidation (N,Q) as variable modification. For further data analysis the MaxQuant LFQ values were loaded into Perseus (https://www.nature.com/articles/nmeth.3901) and a Student’s T-test was performed on LFQ values with false discovery rate 0.01 and S0: 0.1 as significance cut-off.

The mass spectrometry proteomics data have been deposited to the ProteomeXchange Consortium via the PRIDE (PubMed ID: 34723319) partner repository with the dataset identifier PXD045379 (http://www.ebi.ac.uk/pride).

### Membrane topology mapping of FipA

We experimentally determined the membrane orientation of FipA in the membrane using the dual pho-lac reporter system (Karimova et al, 2009), which consists of a translational fusion of the *E. coli* alkaline phosphatase fragment PhoA22-472 and the α-peptide of *E. coli* β-galactosidase, LacZ4-60. A periplasmic localization of the reporter leads to high alkaline phosphatase activity and low β-galactosidase activity, whereas a cytosolic location of the reporter results in high β-galactosidase activity and low alkaline phosphatase activity. Pho-Lac-FipA and FipA-Pho-Lac fusion proteins were ectopically expressed in *E. coli* DH5α grown on a dual-indicator LB medium containing a blue indicator for phosphatase activity (X-Phos) and red indicator for β-galactosidase activity (Red-Gal) (see **Supplementary Figure S1**).

### Bioinformatic analysis

Homologues of FipA were searched using BLAST against the KEGG database, with the sequence of the *V. parahaemolyticus* protein. All homologues containing a DUF2802 were included. The search was later expanded to species known to encode a FlhF homologue (defined as the highest scoring result from a BLAST search using FlhF of *V. parahaemolyticus* as a query, that was also encoded upstream of an FlhG homologue). The flagellation phenotype was later corroborated in the description registered at the List of Prokaryotic names with Standing in Nomenclature (Parte et al. 2020).

## Acknowledgements

We are grateful to Dr. Kathrin Schirner for comments on the manuscript and suggestions for experiments, and Manuel González-Vera for his help on the phylogenetic search. We would like to thank Ulrike Ruppert for great technical support and Jan Heering for construction of plasmid pJH036. This work was supported by the Ludwig-Maximilians-Universität München and the Max Planck Society (SR) and by a grant (TRR 174 P12) from the Deutsche Forschungsgemeinschaft DFG to KMT within the framework of the DFG priority program TRR 174.

## References

Ainsaar, K., H. Tamman, S. Kasvandik, T. Tenson, and R. Hõrak. 2019. “The TonBm-PocAB system is required for maintenance of membrane integrity and polar position of flagella in *Pseudomonas putida*.” J Bacteriol 201: e00303–19. 10.1128/JB.00303-19.

Arroyo-Pérez, E. E., and S. Ringgaard. 2021. “Interdependent polar localization of FlhF and FlhG and their importance for flagellum formation of *Vibrio parahaemolyticus*.” Front Microbiol 12. 10.3389/fmicb.2021.655239.

Bange, G., G. Petzold, K. Wild, R. O. Parlitz, and I. Sinning. 2007. “The crystal structure of the third signal-recognition particle GTPase FlhF reveals a homodimer with bound GTP.” Proc Natl Acad Sci U S A 104: 13621–25. 10.1073/pnas.0702570104.

Blagotinsek, V., M. Schwan, W. Steinchen, D. Mrusek, J. C. Hook, F. Rossmann, S. A. Freibert, et al. 2020. “An ATP-dependent partner switch links flagellar C-ring assembly with gene expression.” Proc Natl Acad Sci U S A 117: 20826–35. 10.1073/pnas.2006470117.

Cameron, T. A., J. Anderson-Furgeson, J. R. Zupan, J. J. Zik, and P. C. Zambryski (2014) Peptidoglycan synthesis machinery in *Agrobacterium tumefaciens* during unipolar growth and cell division. mBio 5: e01219–14.

Campos-García, J., R. Nájera, L. Camarena, and G. Soberón-Chávez. 2000. “The *Pseudomonas aeruginosa* MotR gene involved in regulation of bacterial motility.” FEMS Microbiol Lett 184: 57–62. 10.1111/j.1574-6968.2000.tb08990.x.

Chanchal, P. Banerjee, S. Raghav, H. N. Goswami, and D. Jain. 2021. “The antiactivator FleN uses an allosteric mechanism to regulate σ54-dependent expression of flagellar genes in *Pseudomonas aeruginosa*.” Sci Adv 7: eabj1792. 10.1126/sciadv.abj1792.

Chen, M., Z. Zhao, J. Yang, K. Peng, M. A.B. Baker, F. Bai, and C. J. Lo. 2017. “Length-dependent flagellar growth of *Vibrio alginolyticus* revealed by real time fluorescent imaging.” eLife 6: 1–16. 10.7554/eLife.22140.

Chevance, F. F. V., and K. T. Hughes. 2008. “Coordinating assembly of a bacterial macromolecular machine.” Nat Rev Microbiol 6: 455–65. 10.1038/nrmicro1887.

Coil, D. A., and J. Anné. 2010. “The role of FimV and the importance of its tandem repeat copy number in twitching motility, pigment production, and morphology in *Legionella pneumophila*.” Arch Microbiol 192: 625–31. 10.1007/s00203-010-0590-8.

Correa, N. E, F. Peng, and K. E. Klose. 2005. “Roles of the regulatory proteins FlhF and FlhG in the *Vibrio cholerae* flagellar transcription hierarchy.” J Bacteriol 187: 6324–32. 10.1128/JB.187.18.6324-6332.2005.

Dasgupta, N., and R. Ramphal. 2001. “Interaction of the antiactivator FleN with the transcriptional activator FleQ regulates flagellar number in *Pseudomonas aeruginosa*.” J Bacteriol 183: 6636–44. 10.1128/JB.183.22.6636-6644.2001.

Donnenberg, M. S. and J. B. Kaper. 1991. “Construction of an eae deletion mutant of enteropathogenic *Escherichia coli* by using a positive-selection suicide vector. J Bacteriol 59: 4310–4317.

Dornes, A., L. M. Schmidt, C.-N. Mais, J. C. Hook, J. Pané-Farré, D. Kressler, K. M. Thormann and G. Bange. 2024. „Polar confinment of a macromolecular machine by an SRP-type GTPase.” Nat Comm 15:5797 10.1038/s41467-024-50274-4

Fogel, M. A., and M.K. Waldor. 2006. “A dynamic, mitotic-like mechanism for bacterial chromosome segregation.” Genes Dev 20: 3269–82. 10.1101/gad.1496506.tion.

Francis, N. R., G. E. Sosinsky, D. Thomas, and D. J. DeRosier. 1994. “Isolation, characterization and structure of bacterial flagellar motors containing the switch complex.” J Mol Biol 235: 1261–70; 10.1006/jmbi.1994.1079.

Fredrickson, J. K., J. M. Zachara, D. W. Kennedy, H. Dong, T. C. Onstott, N. W. Hinman, and S.-M. Li. 1998. “Biogenic iron mineralization accompanying the dissimilatory reduction of hydrous ferric oxide by a groundwater bacterium.” Geochim Cosmochim Acta 62: 3239–57. 10.1016/S0016-7037(98)00243-9.

Gao, T., M. Shi, L. Ju, and H. Gao. 2015. “Investigation into FlhFG reveals distinct features of FlhF in regulating flagellum polarity in *Shewanella oneidensis*.” Mol Microbiol 98: 571–85. 10.1111/mmi.13141.

Gibson D. G., L. Young, R. Y. Chuang, J. C. Venter, C. A. Hutchison 3rd, H. O. Smith. “Enzymatic assembly of DNA molecules up to several hundred kilobases”. Nat Methods 6: 343–5. doi: 10.1038/nmeth.1318.

Green, J. C. D., C. Kahramanoglou, A. Rahman, A. M. C. Pender, N. Charbonnel, and G. M. Fraser. 2009. “Recruitment of the earliest component of the bacterial flagellum to the old cell division pole by a membrane-associated signal recognition particle family GTP-binding protein.” J Mol Biol 391: 679–90. 10.1016/j.jmb.2009.05.075.

Heering, J. and S. Ringgaard. 2016. Differential localization of chemotactic signaling arrays during the lifecycle of *Vibrio parahaemolyticus*. Front. Microbiol. 7: 1767.

Heering, J., A. Alvarado, and S. Ringgaard. 2017. Induction of cellular differentiation and single cell imaging of *Vibrio parahaemolyticus* swimmer and swarmer cells. J. Vis. Exp. e55842 (2017).

Hendrixson, D. R., and V. J. DiRita. 2003. “Transcription of σ54-dependent but not σ28-dependent flagellar genes in *Campylobacter jejuni* is associated with formation of the flagellar secretory apparatus.” Mol Microbiol 50: 687–702. 10.1046/j.1365-2958.2003.3731.x.

Hintsche, M., V. Waljor, R. Großmann, M. J. Kühn, K. M. Thormann, F. Peruani, and C. Beta. 2017. “A polar bundle of flagella can drive bacterial swimming by pushing, pulling, or coiling around the cell body.” Sci Rep 7: 16771. 10.1038/s41598-017-16428-9.

Homma, M., S. Aizawa, G. E. Dean, and R. M. Macnab. 1987. “Identification of the M-ring protein of the flagellar motor of *Salmonella typhimurium*.” Proc Natl Acad Sci U S A 84: 7483–7. 10.1073/pnas.84.21.7483.

Hook, J. C., V. Blagotinsek, J. Pané-Farré, D. Mrusek, F. Altegoer, A. Dornes, M. Schwan, L. Schier, K. M. Thormann, and G. Bange. 2020. “A proline-rich element in the type III secretion protein FlhB contributes to flagellar biogenesis in the beta- and gamma-proteobacteria.” Front Microbiol 11: 564161 https://www.frontiersin.org/articles/10.3389/fmicb.2020.564161.

Iyer, S. C., D. Casas-Pastor, D. Kraus, P. Mann, K. Schirner, T. Glatter, G. Fritz, and S. Ringgaard. 2020. “Transcriptional regulation by σ factor phosphorylation in bacteria.” Nat Microbiol 5: 395–406. 10.1038/s41564-019-0648-6.

Karimova, G., J. Pidoux, A. Ullmann, and D. Ladant. 1998. “A bacterial two-hybrid system based on a reconstituted signal transduction pathway.” Proc Natl Acad Sci U S A 95: 5752–56. 10.1073/pnas.95.10.5752.

Karimova, G., C. Robichon, D. Ladant, Characterization of YmgF, a 72-residue inner membrane protein that associates with the *Escherichia coli* cell division machinery. J. Bacteriol. 91: 333– 346 (2009).

Kazmierczak, B. I., and D. R. Hendrixson. 2013. “Spatial and numerical regulation of flagellar biosynthesis in polarly flagellated bacteria.” Mol Microbiol 88: 655–63. 10.1111/mmi.12221.

Kim, Y. K., and L. L. McCarter. 2000. “Analysis of the polar flagellar gene system of *Vibrio parahaemolyticus*.” J Bacteriol 182: 3693–3704. 10.1128/JB.182.13.3693-3704.2000.

Kojima, S., H. Terashima, and M. Homma. 2020. “Regulation of the single polar flagellar biogenesis.” Biomolecules 10: 533. 10.3390/biom10040533.

Kondo, S., M. Homma, and S. Kojima. 2017. “Analysis of the GTPase motif of FlhF in the control of the number and location of polar flagella in *Vibrio alginolyticus*.” Biophys Physicobiol 14: 173–81. 10.2142/biophysico.14.0_173.

Kondo, S. Y. Imura, A. Mizuno, M. Homma, and S. Kojima. 2018. “Biochemical analysis of GTPase FlhF which controls the number and position of flagellar formation in marine Vibiro.” Sci Rep 8: 1–12. 10.1038/s41598-018-30531-5.

Kusumoto, A., K. Kamisaka, T. Yakushi, H. Terashima, A. Shinohara, and M. Homma. 2006. “Regulation of polar flagellar number by the FlhF and FlhG genes in *Vibrio alginolyticus*.” J. Biochem 139: 113–21. 10.1093/jb/mvj010.

Kusumoto, A., N. Nishioka, S. Kojima, and M. Homma. 2009. “Mutational analysis of the GTP-Binding motif of FlhF which regulates the number and placement of the polar flagellum in *Vibrio alginolyticus*.” J Biochem 146: 643–50. 10.1093/jb/mvp109.

Kühn, M. J., F. K. Schmidt, B. Eckhardt, and K. M. Thormann. 2017. “Bacteria exploit a polymorphic instability of the flagellar filament to escape from traps.” Proc Natl Acad Sci 114: 6340–45. 10.1073/pnas.1701644114.

Lassak, J., A.-L. Henche, L. Binnenkade, and K. M. Thormann. 2010. “ArcS, the cognate sensor kinase in an atypical Arc system of *Shewanella oneidensis* MR-1.” Appl Environl Microbiol 76: 3263–74. 10.1128/AEM.00512-10.

Macnab, R M. 2003. “How bacteria assemble flagella.” Annu Rev Microbiol 57. 77–100. 10.1146/annurev.micro.57.030502.090832.

McCarter, L. L. 1995. “Genetic and molecular characterization of the polar flagellum of *Vibrio parahaemolyticus*.” J Bacteriol 177: 1595–1609. 10.1128/jb.177.6.1595-1609.1995.

Miller, V, L., J. J. Mekalanos. 1988. A novel suicide vector and its use in construction of insertion mutations: osmoregulation of outer membrane proteins and virulence determinants in *Vibrio cholerae* requires *toxR*. J Bacteriol 170: 2575–2583.

Milton, D. L., R. O’Toole, P. Hörstedt, and H. Wolf-Watz. 1996. “Flagellin A is essential for the virulence of *Vibrio anguillarum*.” J Bacteriol 178: 1310–19. 10.1128/jb.178.5.1310-1319.1996.

Minamino, T. 2014. “Protein export through the bacterial flagellar type III export pathway.” Biochim Biophys Acta 1843: 1642–48. 10.1016/j.bbamcr.2013.09.005.

Moisi, M., C. Jenul, S. M. Butler, A. New, S. Tutz, J. Reidl, K. E. Klose, A. Camilli, and S. Schild. 2009. “A novel regulatory protein involved in motility of *Vibrio cholerae*.” J Bacteriol 191: 7027–38. 10.1128/JB.00948-09.

Muraleedharan, S., C. Freitas, P. Mann, T. Glatter, S. Ringgaard. 2018 “A cell length-dependent transition in MinD-dynamics promotes a switch in division-site placement and preservation of proliferating elongated *Vibrio parahaemolyticus* swarmer cells.” Mol Microbiol 109.:365–384.

Murray, T. S., and B. I. Kazmierczak. 2006. “FlhF is required for swimming and swarming in *Pseudomonas aeruginosa*.” J Bacteriol 188: 6995–7004. 10.1128/JB.00790-06.

Navarrete, B., A. Leal-Morales, L. Serrano-Ron, M. Sarrió, A. Jiménez-Fernández, L. Jiménez-Díaz, A. López-Sánchez, and F. Govantes. 2019. “Transcriptional organization, regulation and functional analysis of FlhF and FleN in *Pseudomonas putida*.” PLoS ONE 14: e0214166. 10.1371/journal.pone.0214166.

Nelson, K. E., C. Weinel, I. T. Paulsen, R. J. Dodson, H. Hilbert, V. a. P. Martins dos Santos, D. E. Fouts, et al. 2002. “Complete genome sequence and comparative analysis of the metabolically versatile *Pseudomonas putida* KT2440.” Environ Microbiol 4: 799–808. 10.1046/j.1462-2920.2002.00366.x.

Pandza, S., M. Baetens, C. H. Park, T. Au, M. Keyhan, and A. Matin. 2000. “The G-protein FlhF has a role in polar flagellar placement and general stress response induction in *Pseudomonas putida*.” Mol Microbiol 36: 414–23. 10.1046/j.1365-2958.2000.01859.x.

Parte, Aidan C., J. Sardà Carbasse, J. P. Meier-Kolthoff, L. C. Reimer, and M. Göker. 2020. “List of prokaryotic names with standing in nomenclature (LPSN) moves to the DSMZ.” Intl J Syst Evol Microbiol 70: 5607–12. 10.1099/ijsem.0.004332.

Petersen, B. D., M. S. Liu, R. Podicheti, A. Ying-Po Yang, C. A. Simpson, C. Hemmerich, D. B. Rusch, and J. C. van Kessel. 2021. “The polar flagellar transcriptional regulatory network in *Vibrio campbellii* deviates from canonical *Vibrio* species.” J Bacteriol 203: e00276–21. 10.1128/JB.00276-21.

Ringgaard, S., K. Schirner, B. M. Davis, and M. K. Waldor. 2011. “A family of ParA-like ATPases promotes cell pole maturation by facilitating polar localization of chemotaxis proteins.” Genes Dev 25: 1544–55. 10.1101/gad.2061811.

Ringgaard, S., et al., ParP prevents dissociation of CheA from chemotactic signaling arrays and tethers them to a polar anchor. Proc Natl Acad Sci U S A 111: E255–E264 (2013).

Rossmann, F., S. Brenzinger, C. Knauer, A. K. Dörrich, S. Bubendorfer, U. Ruppert, G. Bange, and K. M. Thormann. 2015. “The Role of FlhF and HubP as polar landmark proteins in *Shewanella putrefaciens* CN-32.” Mol Microbiol 98: 727–42. 10.1111/mmi.13152.

Rossmann, F. M., T. Rick, D. Mrusek, L. Sprankel, A. K. Dörrich, T. Leonhard, S. Bubendorfer, V. Kaever, G. Bange, and K. M. Thormann. 2019. “The GGDEF Domain of the phosphodiesterase PdeB in Shewanella putrefaciens mediates recruitment by the polar landmark protein HubP.” J. Bacteriol 201: e00534–18. 10.1128/JB.00534-18.

Schuhmacher, J. S., F. Rossmann, F. Dempwolff, C. Knauer, F. Altegoer, W. Steinchen, A. K. Dörrich, et al. 2015. “MinD-like ATPase FlhG effects location and number of bacterial flagella during C-ring assembly.” Proc Natl Acad Sci U S A 112: 3092–97. 10.1073/pnas.1419388112.

Schuhmacher, J. S., K. M. Thormann, and G. Bange. 2015. “How bacteria maintain location and number of flagella?” FEMS Microbiol Rev 39: 812–22. 10.1093/femsre/fuv034.

Schwan, M., A. Khaledi, S. Willger, K. Papenfort, T. Glatter, S. Häußler, and K. M. Thormann. 2022. “FlrA-Independent production of flagellar proteins is required for proper flagellation in *Shewanella putrefaciens*.” Mol Microbiol 118: 670–82. 10.1111/mmi.14993.

Takekawa, N., S. Kwon, N. Nishioka, S. Kojima, and M. Homma. 2016. “HubP, a polar landmark protein, regulates flagellar number by assisting in the proper polar localization of FlhG in *Vibrio alginolyticus*.” J Bacteriol 198: JB.00462-16. 10.1128/JB.00462-16.

Turriziani, B., A. Garcia-Munoz, R. Pilkington, C. Raso, W. Kolch, and A. Von Kriegsheim. 2014. “On-beads digestion in conjunction with data-dependent mass spectrometry: A shortcut to quantitative and dynamic interaction proteomics.” Biology 3: 320–32. 10.3390/biology3020320.

Ueno, T., K. Oosawa, and S. Aizawa. 1992. “M ring, S sing and proximal rod of the flagellar basal body of *Salmonella typhimurium* are composed of subunits of a single protein, FliF.” J Mol Biol 27: 672. 10.1016/0022-2836(92)90216-7.

Wehbi, H., E. Portillo, H. Harvey, A. E. Shimkoff, E. M. Scheurwater, P. L. Howell, and L. L. Burrows. 2011. “The peptidoglycan-binding protein FimV promotes assembly of the *Pseudomonas aeruginosa* type IV pilus secretin.” J Bacteriol 193: 540–50. 10.1128/JB.01048-10.

Yamaichi, Y., R. Bruckner, S. Ringgaard, A. Möll, D. E. Cameron, A. Briegel, G. J. Jensen, B. M. Davis, and M. K. Waldor. 2012. “A multidomain hub anchors the chromosome segregation and chemotactic machinery to the bacterial pole.” Genes Dev 26: 2348–60. 10.1101/gad.199869.112.

Zhang, K., J. He, C. Catalano, Y. Guo, J. Liu, and C. Li. 2020. “FlhF regulates the number and configuration of periplasmic flagella in *Borrelia burgdorferi*.” Mol Microbiol 113: 1122–39. 10.1111/mmi.14482.

Zheng, L., U. Baumann, and J.-L. Reymond. 2004. “An efficient one-step site-directed and site-saturation mutagenesis protocol.” Nucleic Acids Res 32: e115. 10.1093/nar/gnh110.

