## Supplementary Figures 1 - 15; Supplementary Tables 1 - 5 for "A conserved cell-pole determinant organizes proper polar flagellum formation"

[1]Max Planck Institute for Terrestrial Microbiology, Department of Ecophysiology, Marburg 35043, Germany; [2]Department of Biology I, Microbiology, Ludwig-Maximilians-Universität München, Munich, Germany; [3]Department of Microbiology and Molecular Biology, Justus-Liebig-Universität Gießen, Giessen, Germany; [4]Interfaculty Institute of Microbiology and Infection Medicine Tübingen, Bacterial Metabolomics, University of Tübingen, Tübingen, Germany; [5]Institute for Biological Physics, University of Cologne, Köln, Germany; [6]Core Facility for Mass Spectrometry and Proteomics, Max Planck Institute for Terrestrial Microbiology, Marburg, Germany.

### Supplementary Material

+ These authors contributed equally to this work

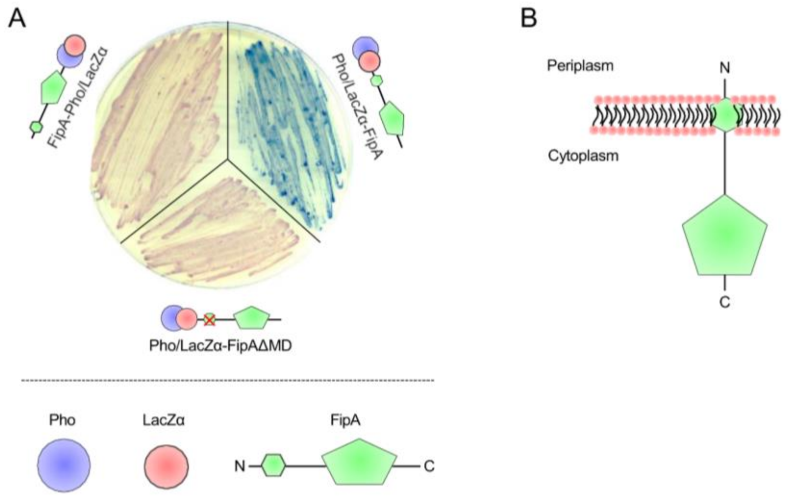


**Supplementary Figure 1: Membrane topology of FipA. A) Topology analysis and (B) the corresponding model.** A Pho/LacZα cassette was translationally fused to the N- or the C-terminus of FipA and the fusion is expressed in a suitable *E. coli* strain. Pho is only active in the periplasm, while LacZα can functionally complement β-galactosidase activity within the cytoplasm. The enzyme activities can then be tested using suitable color reactions. Expression of Pho-Lac-FipA resulted in blue colonies whereas expression of FipA-Pho-Lac resulted in red colonies, hence suggesting that the FipA N-terminus is positioned in the periplasm and the C-terminus in the cytoplasm **(B)**. Deletion of the predicted membrane spanning domain in Pho-Lac-FipA (aa 6-28, Pho-Lac-FipAΔMD) resulted in red *E. coli* colonies (A). This suggests that in the absence of the membrane spanning domain the N-terminus of FipA resides in the cytoplasm, and thus further supports that FipA is a transmembrane protein anchored in the inner membrane via the 6-28 aa N-terminal membrane spanning domain and is oriented with the 1-5 aa N-terminal in the periplasm and the 29-163 aa C-terminal in the cytoplasm (B).

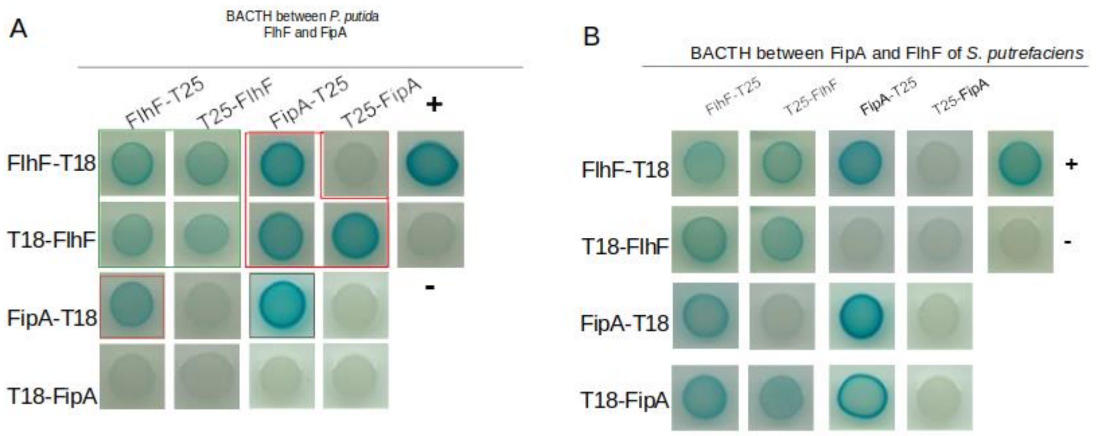


**Supplementary Figure 2: FlhF interacts with FipA in a Bacterial Two-Hybrid Analysis (BACTH) in *P. putida* and *S. putrefaciens*.** FlhF and FipA were produced as N-terminal or as C-terminal fusions to the T18 and T25 fragments of the *B. pertussis* adenylate cyclase in the given combinations within a single strain. *In vivo* protein-protein interactions result in active adenylate cyclase activity and high cAMP levels, indicated by blue coloring of the colonies on X-Gal-containing agar.

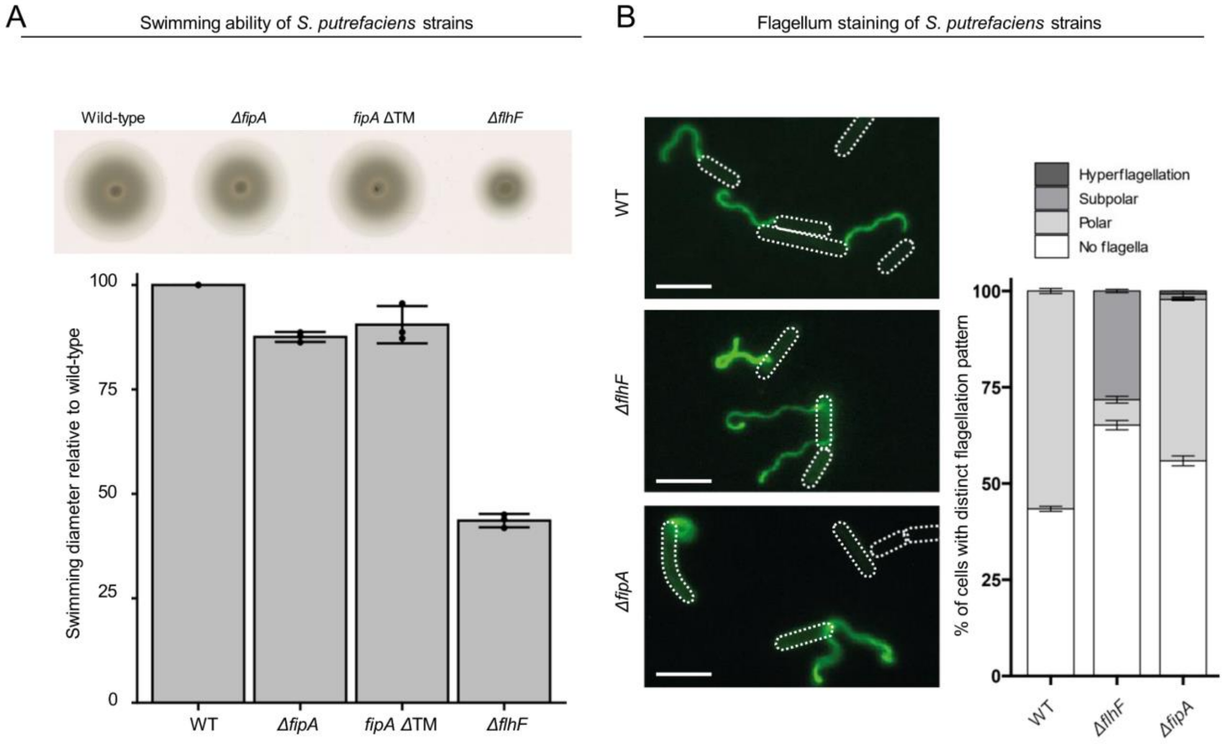


**Supplementary Figure 3: Deletions of or in FipA and FlhF affect *S. putrefaciens* flagellation. A) Upper panel:** Spreading of the indicated strains in soft agar. *fipA* ΔTM indicates that the transmembrane region of FipA was deleted. The images shown were compiled from a single plate. **Lower panel:** the corresponding quantification given as diameter; three independent experiments were conducted. The error bar marks the standard deviation. **B)** Left: Shown are fluorescent micrographs where the flagellar filament(s) of the indicated strains were fluorescently labeled by maleimide dye. The position of the cell bodies was outlined in the figure. The scale bare equals 5 µm. Right: The corresponding quantification of flagellation. The number of cells (n) for each experiment was > 300 from 3 indepenedent experiments. The error bar between the different populations shows the standard deviation.

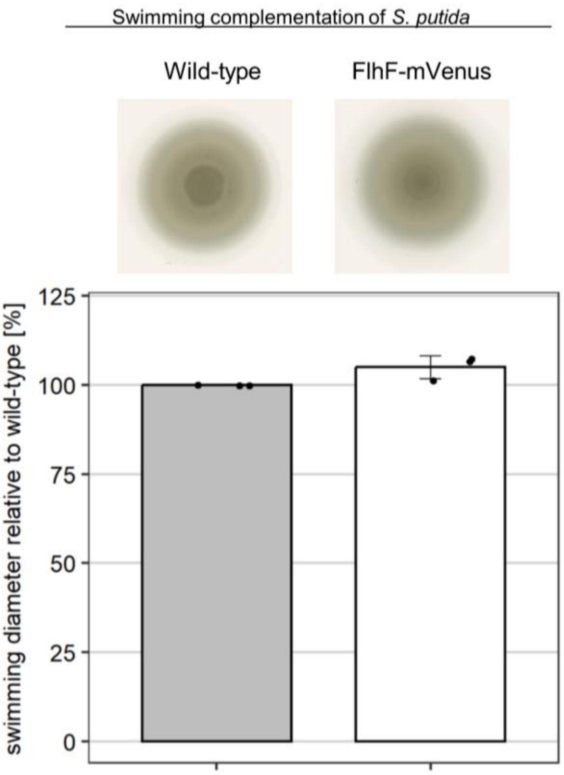


**Supplementary Figure 4: N-terminal tagging of *Pp*FlhF with mCherry does not negatively affect spreading motility in soft agar**. The upper panel shows the spreading in soft agar, the lower panel shows the corresponding quantification. The images shown are from the same plate, three independent experiments were conducted. The corresponding standard error bar is shown.

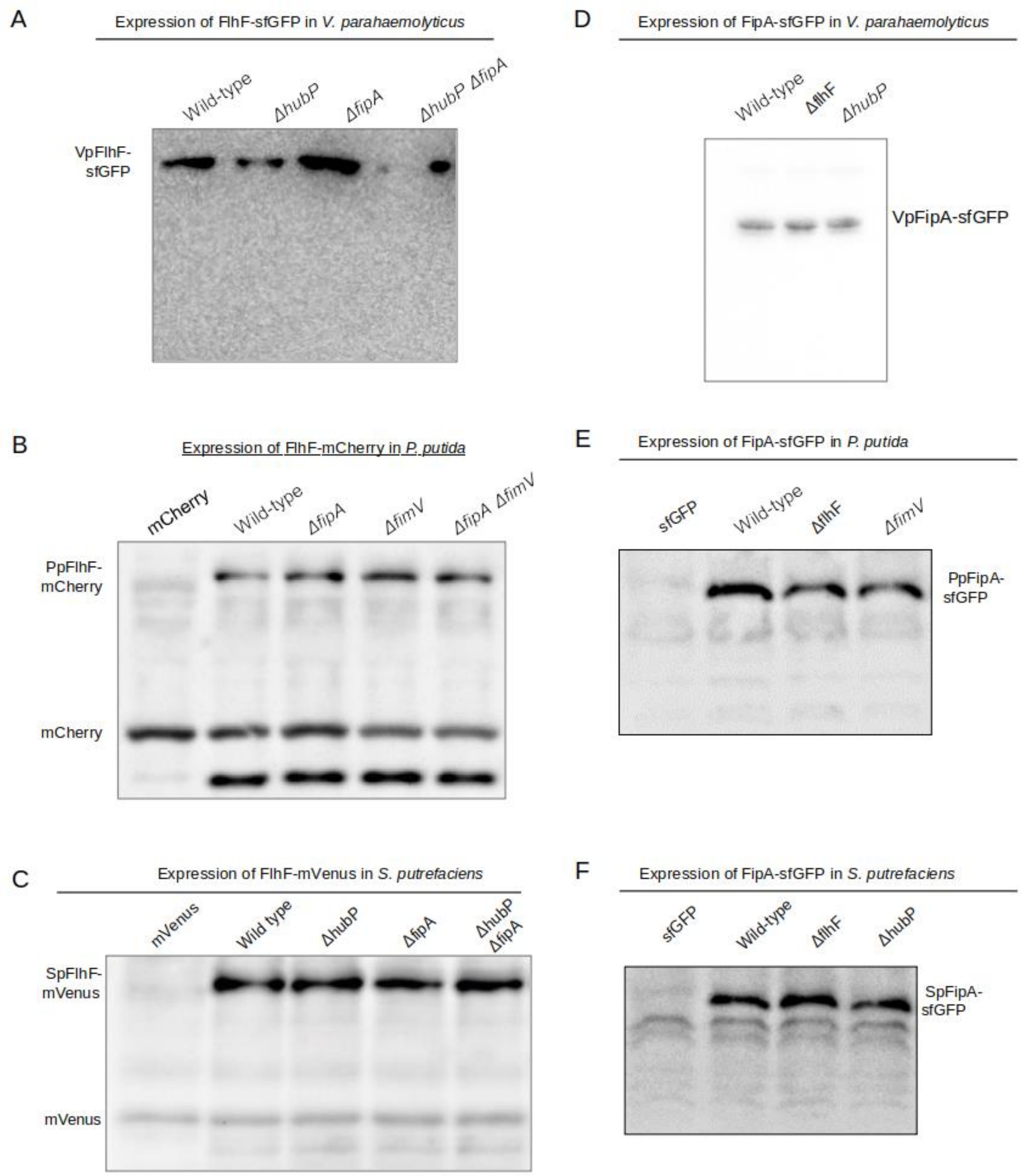


**Supplementary Figure 5: Western analysis on the protein levels of FlhF and FipA fusions to fluorescent proteins.** Crude protein extracts from the corresponding (mutant) strains of *V. parahaemolyticus* **(A D)**, *P. putida* **(B, E)** and *S. putrefaciens* **(C, F)** were separated by SDS-PAGE, transferred to membranes and visualized by western blotting using antibodies against GFP (**A, C-F**) and mCherry (**B**). The same OD units from the culture were loaded for each sample. The corresponding strain background is indicated above the lanes.

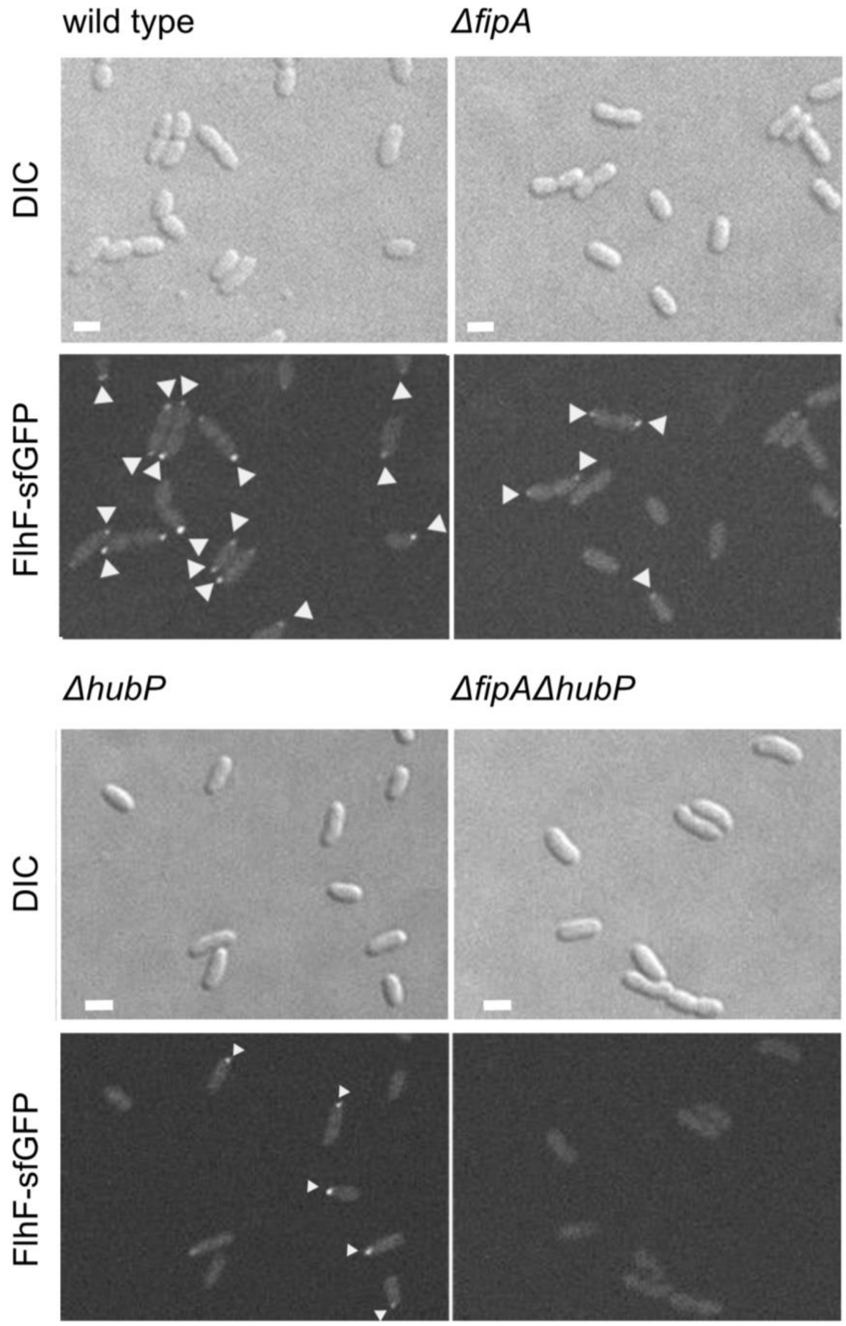


**Supplementary Figure 6: Localization of FlhF depends on FipA and HubP.** Shown are enlargments of Fig. 3A, representative micrographs of the indicated strains of *V. parahaemoloyticus* expressing FlhF-sfGFP from its native promoter. The upper panel shows the DIC image, the lower panel the corresponding fluorescence image. Fluorescent foci are highlighted by white arrows. Scale bar = 2 µm.

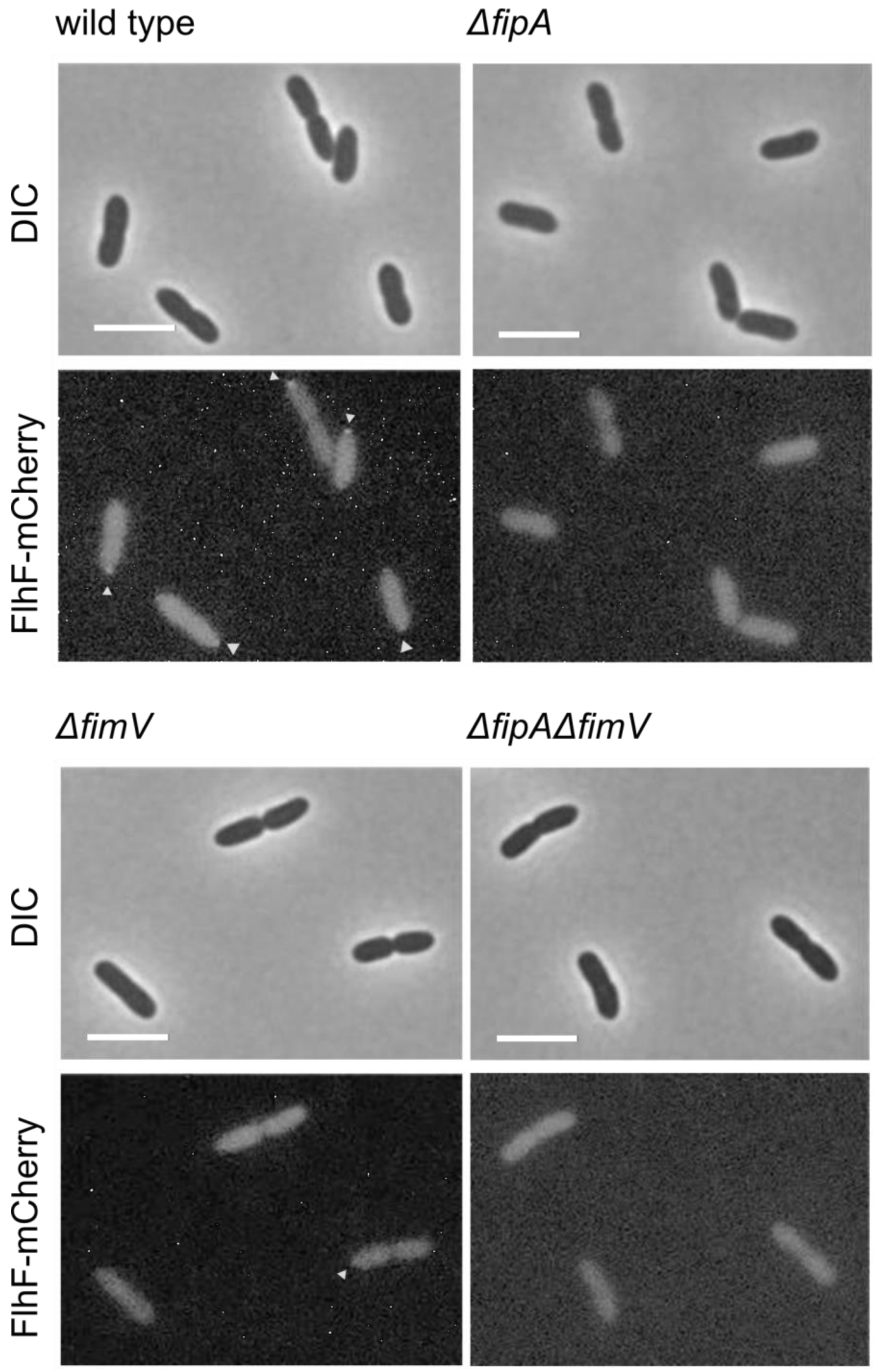


**Supplementary Figure 7: Localization of FlhF depends on FipA and HubP.** Shown are enlargments of Fig. 3E, representative micrographs of the indicated strains of *P. putida* expressing FlhF-sfGFP from its native promoter. The upper panel shows the DIC image, the lower panel the corresponding fluorescence image. Fluorescent foci are highlighted by white arrows. Scale bar = 5 µm.

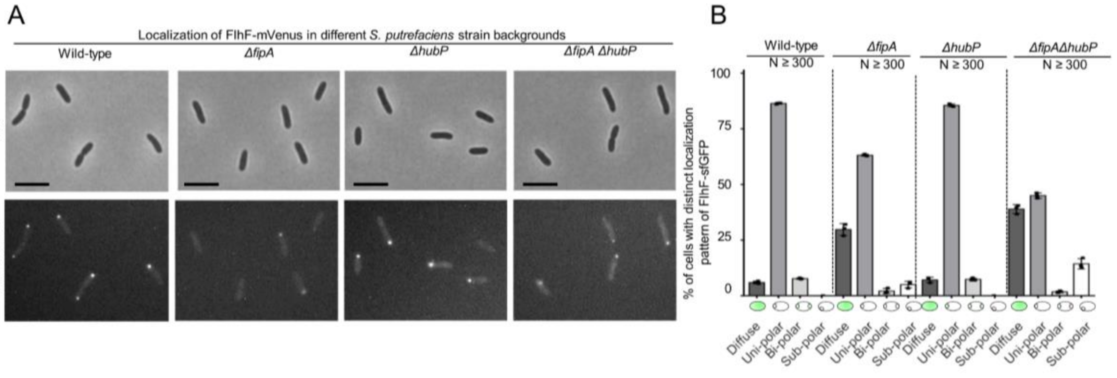


**Supplementary Figure 8: *Sp*FipA and *Sp*HubP concertedly affect polar *Sp*FlhF localization. A)** Fluorescence microscopy on the indicated (mutant) strains each bearing an *flhF-mvenus* hybrid gene replacing native *flhF* on the chromosome. The upper panel shows DIC micrographs, the lower panel the corresponding fluorescent images. The scale bar equals 5 µm. **B)** Quantification of the FlhF-mVenus localization patterns, using the data from > 300 cells (n) from three independent experiments, the corresponding standard error is shown as error bar.

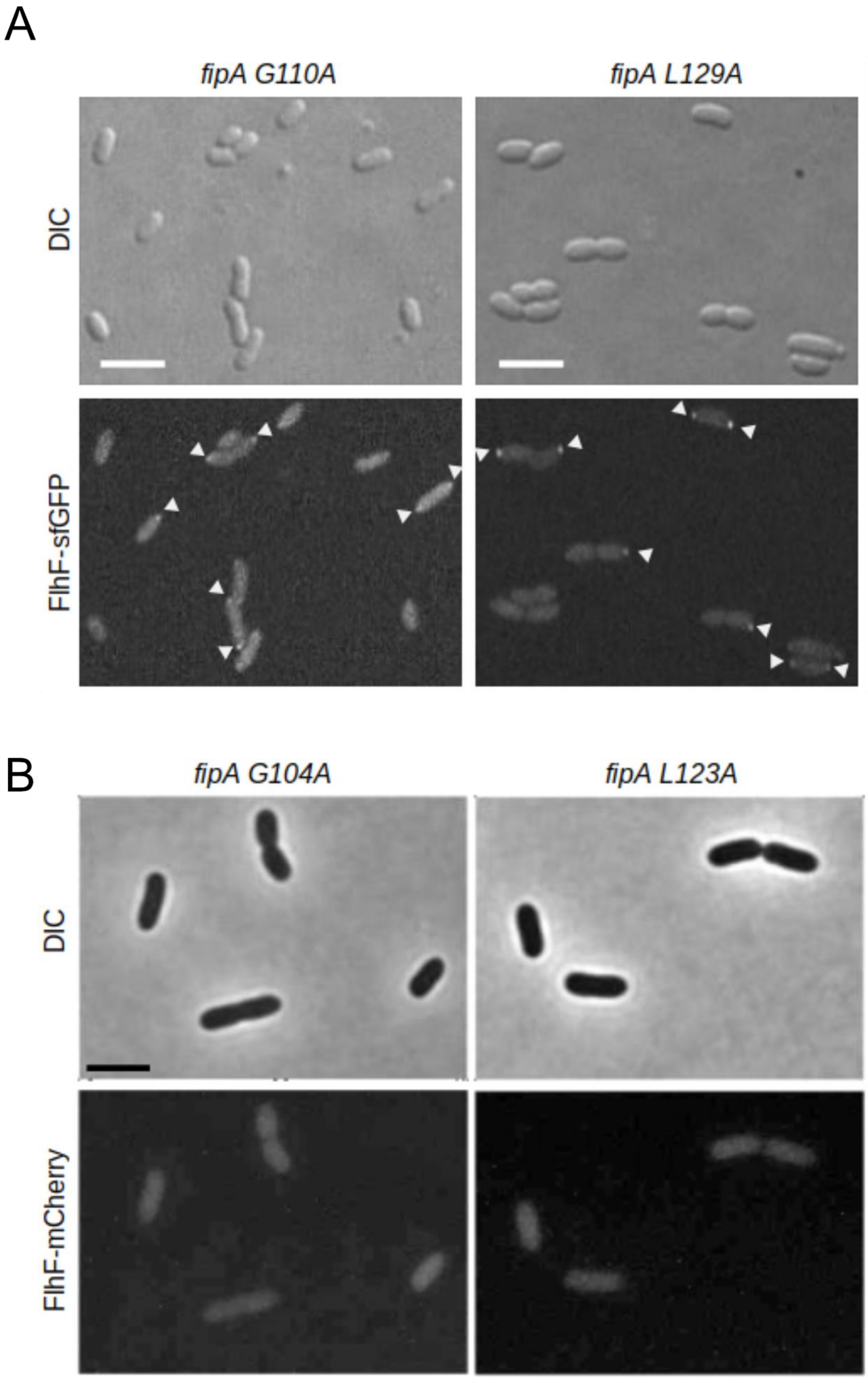


**Supplementary Figure 9: Conserved residues in the domain of unknown function of FipA are essential for interaction with FlhF. A)** Localization of *Vp*FlhF in cells with substitutions in the DUF2802 domain. Micrographs are showing the localization of FipA-sfGFP in the indicated strains; the upper panels display the DIC and the lower panels the corresponding fluorescence images. The scale bar equals 5 µm. **B)** The same analysis for *P. putida* FlhF-mCherry. The images are enlargements of **Fig. 5F (A)** and **Fig. 5G (B)**.

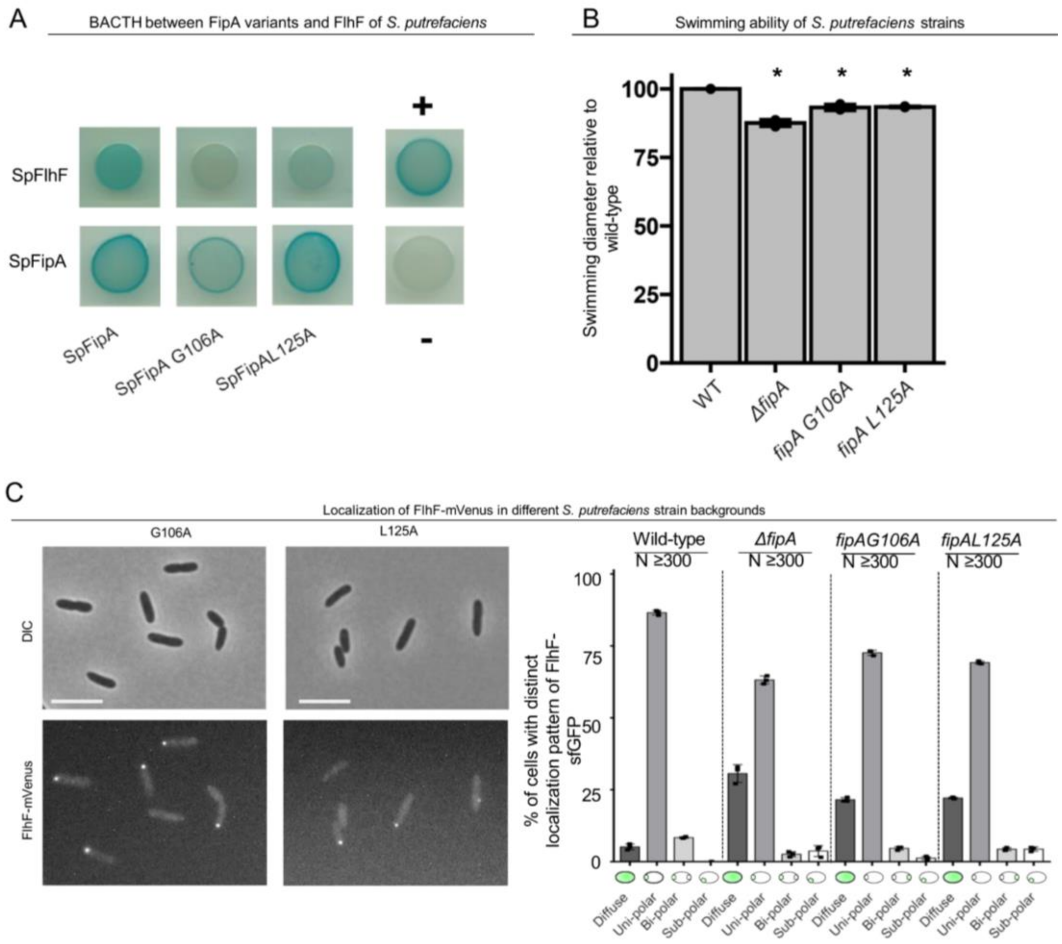


**Supplementary Figure 10: Targeted residue substitution in FipA affects FlhF positioning and function.** **A)** BACTH analysis (see, e.g., Figure S2) suggests that G106A and L125A in *Sp*FipA does not disrupt but maybe weakens (G106A) the interaction between *Sp*FipA and *Sp*FlhF. **B)** Spreading of the indicated *S. putrefaciens* strains in soft agar reveals a small but sigifnificant (asterisk = $p < 0.05$) and based on three independent experiments. **C) Left:** micrographs showing DIC (upper panels) and fluorescent imaging on mVenus-tagged FlhF cells in mutant backgrounds bearing the given substutions in *Sp*FipA with the corresponding quantification (n > 300 cells, **right**). The *in vivo* analysis suggests that the residue substitutions in *Sp*FipA give rise to phenotypes that are similar to that of a *fipA* deletion.

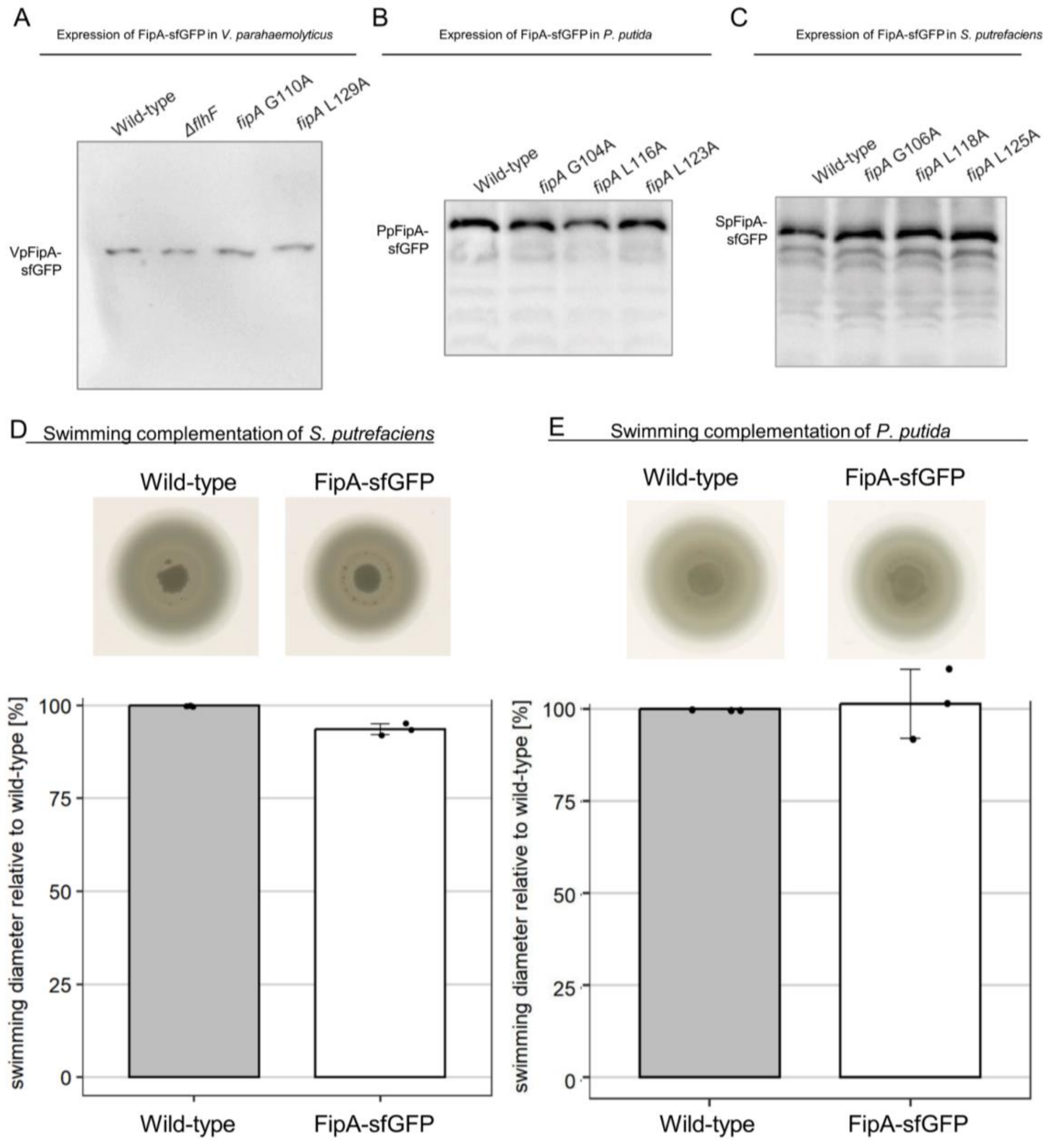


**Supplementary Figure 11: FipA-sfGFP is stably produced despited amino acid substitutions.** The three panels show western blotting analysis of PAGE protein separations from *V. parahaemolyticus* (**A**), *P. putida* (**B**) and *S. putrefaciens* (**C**) bearing FipA-sfGFP with substitutions in conserved residues as indicated, using antibodies raised against sfGFP. V. parahaemolyticus also includes a mutant deleted in *flhF*. The loaded samples are normalized by OD units. **D)** The C-terminal fusion of sfGFP has only minor effects on FipA function in *S. putrefaciens* and *P. putida*. The upper panels show the spreading in soft agar (taken from the same soft agar plate), the lower the corresponding quantification. The error bars are the standard deviation from three independent experiments.

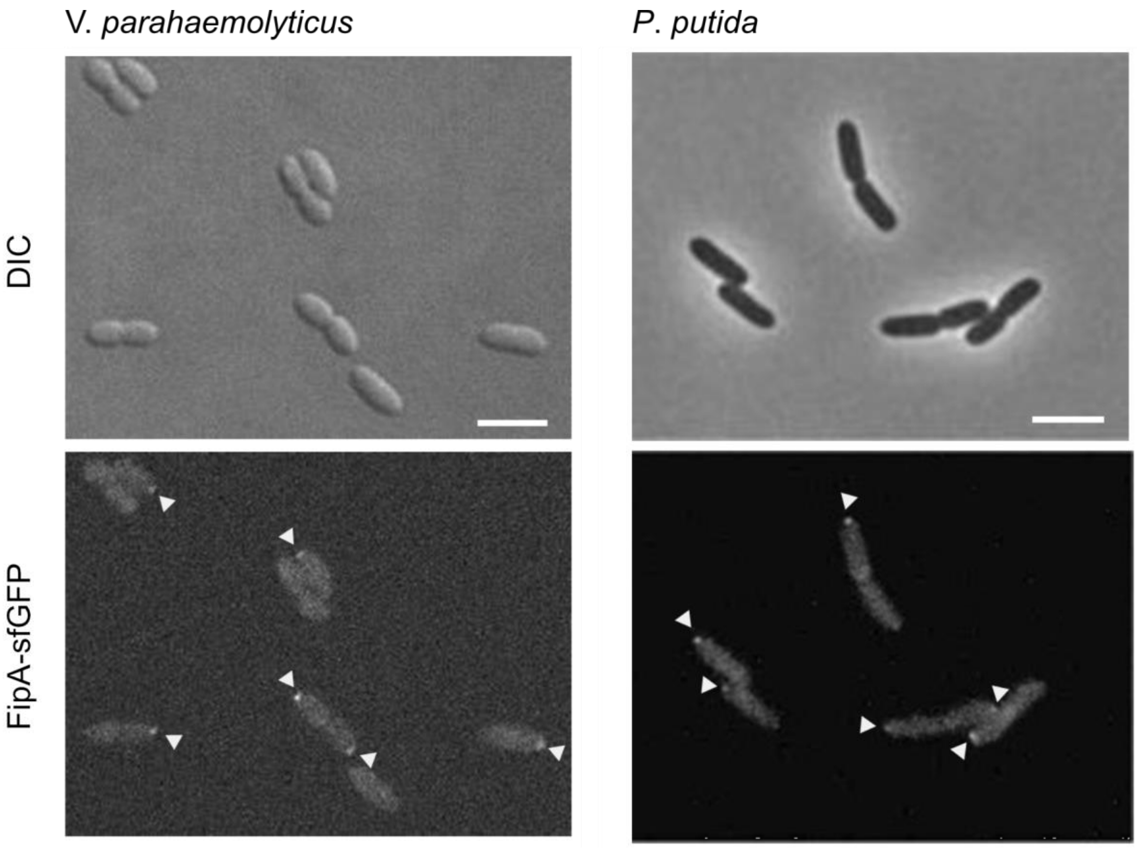


**Supplementary Figure 12: The localization pattern of FipA.** Localization pattern of fluorescently labeled FipA in *V. parahaemolyticus* and *P. putida*. **Left:** Representative micrographs of *V. parahaemolyticus* expressing FipA-sfGFP from its native promoter. Scale bar = 2 µm. The upper panel shows the DIC and the lower panel the corresponding fluorescence channel. **Right:** The same analysis for *P. putida*. The images are enlargements of the micrographs in **Fig. 6A and B**.

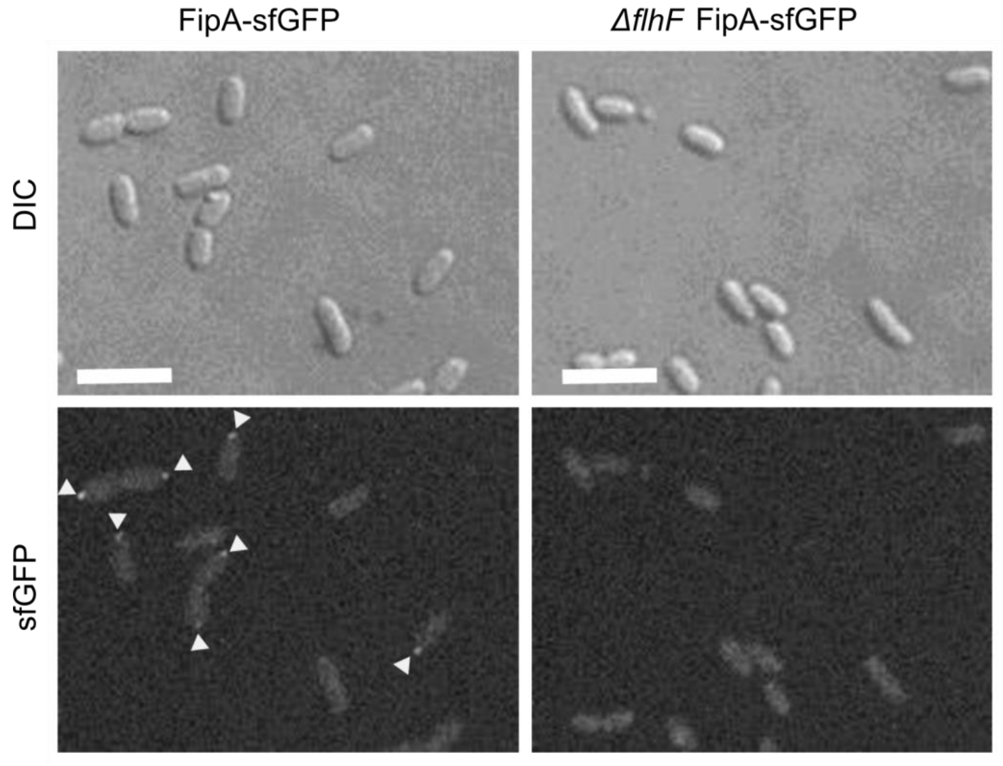


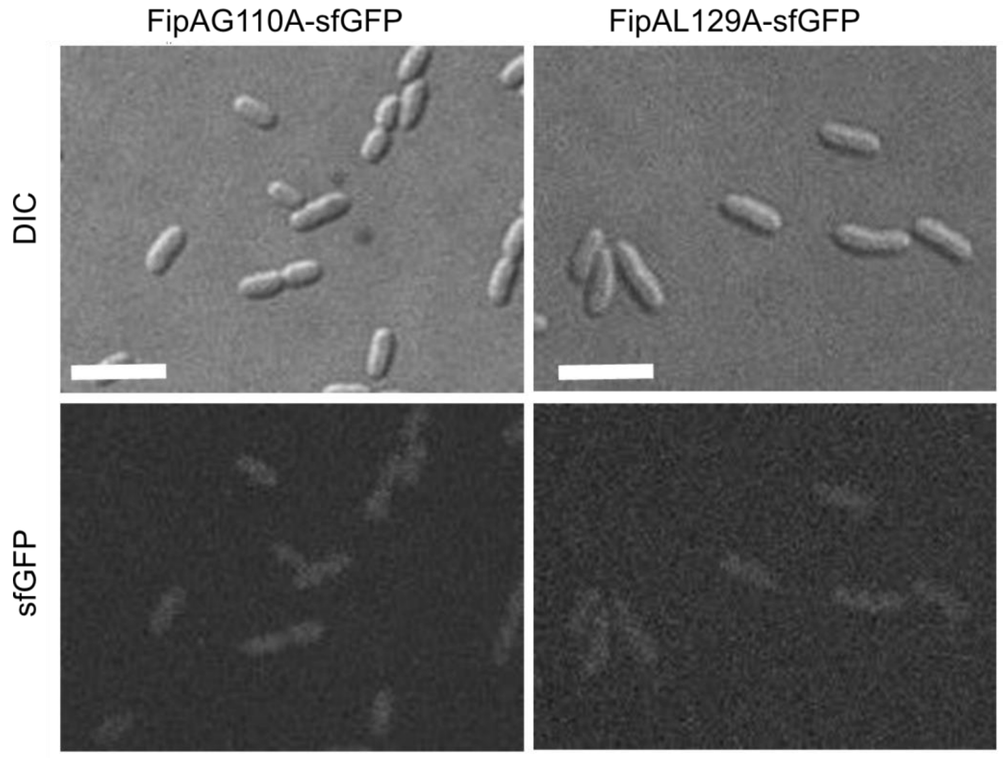


**Supplementary Figure 13: Normal localization of FipA depends on interaction with FlhF.** Localization pattern of *V. parahaemolitycus* FipA-sfGFP in the indicated wild-type and mutant strains. The upper panels display the DIC micrographs, the lower panel the corresponding fluorescence imaging (scale bar equals 5 µm). The images are enlargements of the micrographs in Fig. 7 A (upper) and 7B (lower).

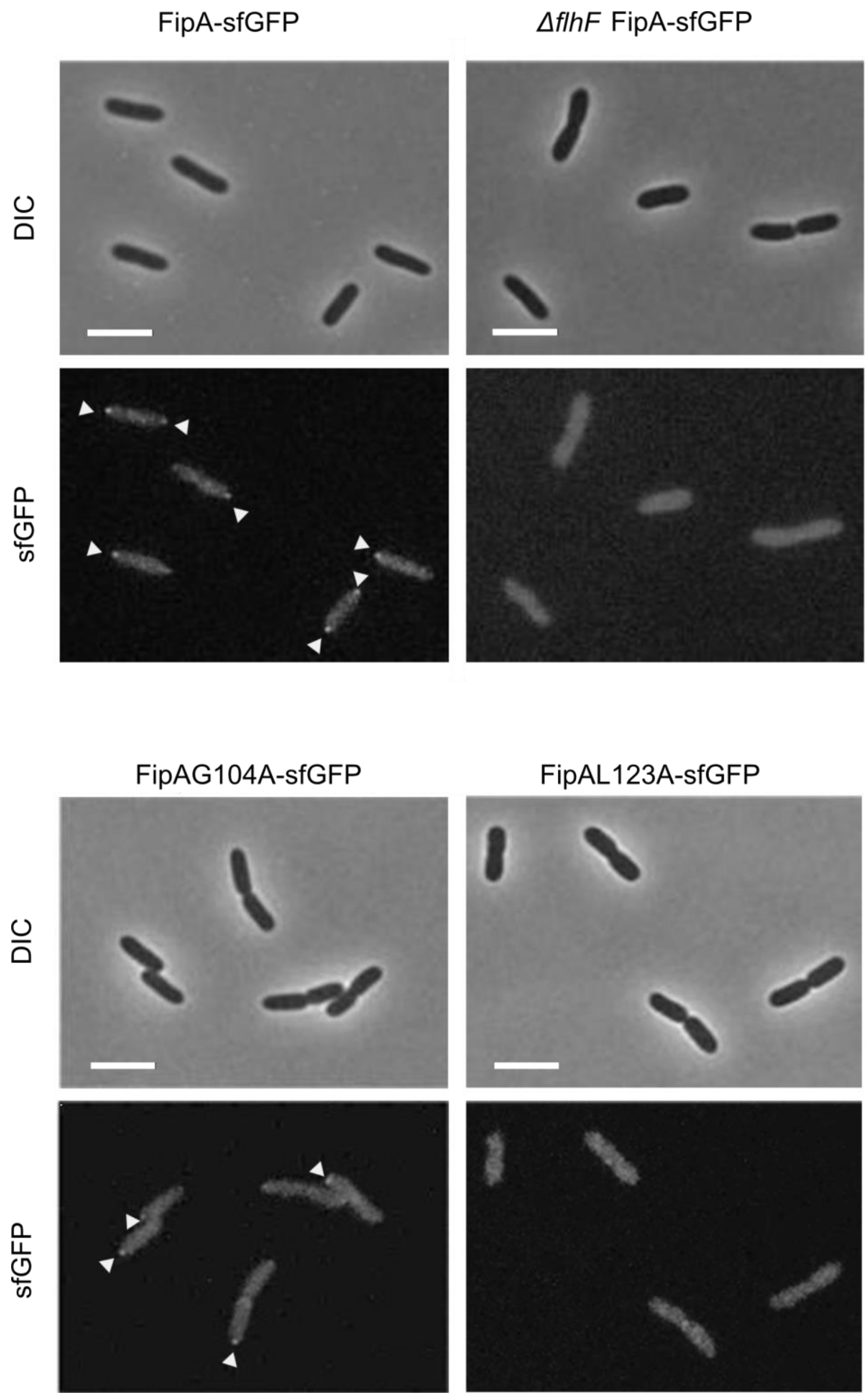


**Supplementary Figure 14: Normal localization of FipA depends on interaction with FlhF.** Localization pattern of *P. putida* FipA-sfGFP in the indicated wild-type and mutant strains. The upper panels display the DIC micrographs, the lower panel the corresponding fluorescence imaging (scale bar equals 5 µm). The images are enlargements of the micrographs in Fig. 7 D (upper) and 7E (lower).

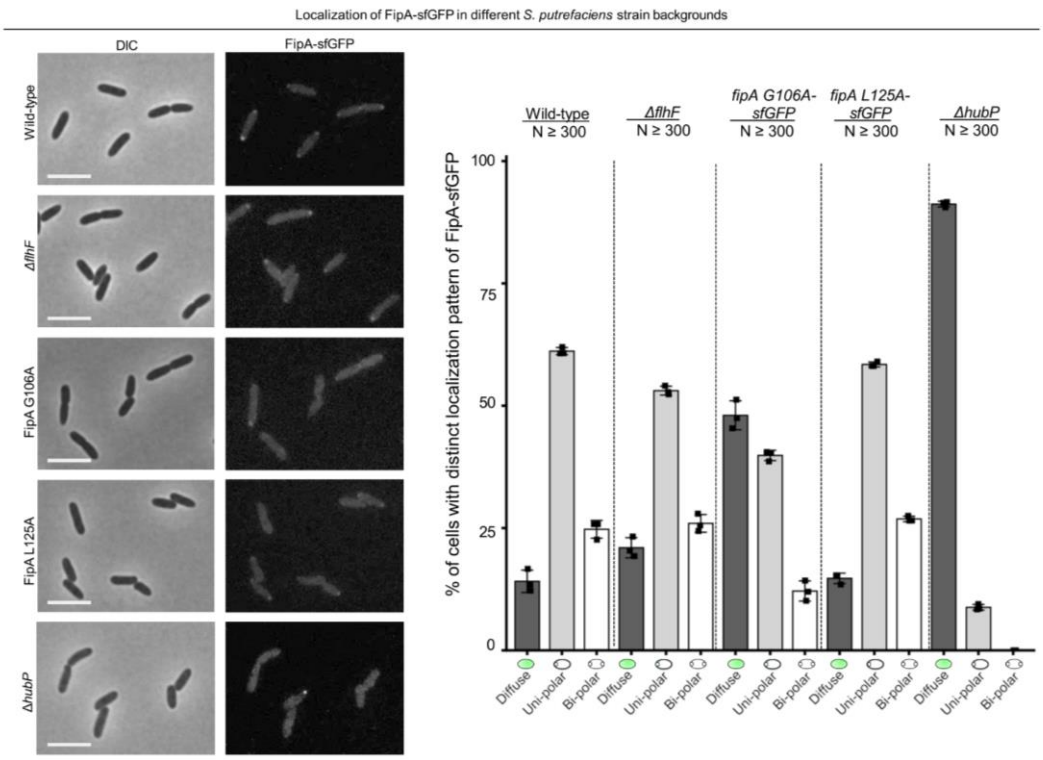


**Supplementary Figure 15: Localization of *Sp*FipA in different strain backgrounds.** Left: Micrographs showing DIC (left) and fluorescence (right) microscopy on cells with FipA-sfGFP in different background strains as indicated. The scale bar equals 5 µm. The corresponding quantification (N > 300, three independent experiments) is shown to the right. FipA loses its polar localization in a mutant lacking HubP and in a mutant with the L125A substitution in FipA.

#### Supplementary Tables

##### Supplementary Table S1: Enriched proteins in Co-IP FlhF-sfGFP vs. sfGFP

| Protein names | Gene name | log2-Fold change (FlhF - GFP) |
|---|---|---|
| **FlhF** | VP2234 | 11.33 |
| | VP0127 | 8.78 |
| | VPA0809 | 8.59 |
| **FliF** | VP2249 | 8.43 |
| | VPA0808 | 8.40 |
| | VPA0807 | 7.85 |
| | VP0700 | 7.42 |
| **FliG** | VP2248 | 7.39 |
| **FliE** | fliE | 6.72 |
| **FlgB** | VP0775 | 6.18 |
| | VP1353 | 5.91 |
| **FlhA** | VP2235 | 5.87 |
| | VP0944 | 5.65 |
| | VPA0337 | 5.32 |
| **FipA** | VP2224 | 4.85 |
| **FtsI** | VP0454 | 4.77 |
| **FlgC** | VP0776 | 4.74 |
| | VP0974 | 4.31 |
| **DedD** | VP2187 | 4.17 |
| | VPA1077 | 4.00 |
| **FliL** | VP2243 | 3.97 |
| | VP1100 | 3.24 |
| | VP0646 | 3.17 |
| **IbpA** | VP0018 | 2.65 |
| **SecD** | secD | 2.61 |
| **TolC** | VP0425 | 2.51 |
| **FlgF** | VP0780 | 2.50 |
| **DnaJ** | dnaJ | 2.47 |

**Supplementary Table S2: Flagellation pattern and pressence of FipA and FlhF in bacteria**

| Name | flhF | fip A | Flagellation pattern | Reference |
|---|---|---|---|---|
| **Buchnera aphidicola JF99** | No | No | non motile | https://doi.org/10.1099/00207713-41-4-566 |
| **Citrobacter freundii** | No | No | peritrichous | https://doi.org/10.1128%2Fjb.23.2.167-182.1932 |
| **Citrobacter rodentium** | No | No | non motile | https://doi.org/10.1128%2Fjcm.33.8.2064-2068.1995 |
| **Escherichia coli K12 MG1655** | No | No | peritrichous | |
| **Pantoea agglomerans C410P1** | No | No | peritrichous | https://doi.org/10.1099/00207713-39-3-337 |
| **Proteus vulgaris** | No | No | peritrichous | https://doi.org/10.1128/jb.90.5.1337-1354.1965 |
| **Serratia marcescens WW4** | No | No | subpolar | https://dx.doi.org/10.1016%2Fj.resmic.2008.07.003 |
| **Sodalis glossinidius** | No | No | non motile | https://doi.org/10.1099/00207713-49-1-267 |
| **Xenorhabdus nematophila** | No | No | peritrichous | https://doi.org/10.1099/00207713-29-4-352 |
| **Yersinia enterocolitica WA** | No | No | many | |
| **Aliivibrio fischeri ES114** | Yes | Yes | lophotrichous | https://doi.org/10.1128%2FJB.187.6.2058-2065.2005 |
| **Grimontia hollisae** | Yes | Yes | monotrichous | https://doi.org/10.1099/ijs.0.02660-0 |
| **Photobacterium profundum** | Yes | Yes | monotrichous | https://doi.org/10.1007/s007920050036 |
| **vibrio cholerae O1 El Tor N16961** | Yes | Yes | monotrichous | |
| **Vibrio parahaemolyticus** | Yes | Yes | monotrichous | |
| **Alteromonas australica H 17** | Yes | Yes | monotrichous | doi.org/10.1099/00207713-45-4-755 |

| | | | | |
|---|---|---|---|---|
| **Idiomarina loihensis L2TR** | Yes | Yes | monotrichous | https://doi.org/10.1099/ijs.0.02701-0 |
| **Pseudoalteromonas atlantica** | Yes | Yes | monotrichous | doi.org/10.1099/00207713-45-4-755 |
| **Pseudoalteromonas haloplanktis** | Yes | Yes | monotrichous | doi.org/10.1099/00207713-45-4-755 |
| **Pseudoalteromonas luteoviolacea** | Yes | Yes | monotrichous | doi.org/10.1099/00207713-45-4-755 |
| **Pseudoalteromonas rubra** | Yes | Yes | monotrichous | doi.org/10.1099/00207713-45-4-755 |
| **Catenovulum sp. CCB-QB4** | Yes | Yes | peritrichous | https://doi.org/10.1099/ijs.0.027565-0 |
| **Salinimonas sp. HMF8227** | Yes | Yes | monotrichous | https://doi.org/10.1099/ijs.0.63279-0 |
| **Shewanella putrefaciens** | Yes | Yes | monotrichous | |
| **Moritella viscosa** | Yes | Yes | monotrichous | https://doi.org/10.1099/00207713-50-2-479 |
| **Psychromonas ingrahamii** | No | No | non motile | https://doi.org/10.1099/ijs.0.64068-0 |
| **Aeromonas salmonicida** | Yes | Yes | monotrichous | https://doi.org/10.1099/00207713-17-3-273 |
| **Tolumonas auensis** | No | No | non motile | https://doi.org/10.1099/00207713-46-1-183 |
| **Hahella chejuensis** | Yes | Yes | monotrichous | https://doi.org/10.1099/00207713-51-2-661 |
| **Marinomonas sp. MWYL1** | Yes | Yes | monotrichous or amphitrichous | https://doi.org/10.1128/jb.110.1.402-429.1972 |
| **Cobetia marina** | No | No | subpolar | https://doi.org/10.1099/00221287-62-2-159 |
| **Halomonas elongata** | No | No | lophotrichous or peritrichous | https://doi.org/10.1099/00207713-30-2-485 |
| **Azotobacter vinelandii DJ** | Yes | No | peritrichous | https://dx.doi.org/10.1099%2Fmic.0.2008%2F017665-0 |
| **Pseudomonas aeruginosa PAO1** | Yes | Yes | lophotrichous | |

| | | | | |
|---|---|---|---|---|
| **Pseudomonas fluorescens** | Yes | Yes | lophotrichous | https://doi.org/10.1099/mic.0.27362-0 |
| **Pseudomonas putida F1** | Yes | Yes | lophotrichous | |
| **cellvibrio japonicus** | Yes | Yes | monotrichous | https://doi.org/10.1099/ijs.0.02271-0 |
| **Microbulbifer aggregans** | No | No | non motile | https://doi.org/10.1099/ijsem.0.002258 |
| **Saccharophagus degradans** | Yes | Yes | monotrichous | https://doi.org/10.1099/ijs.0.63627-0 |
| **Teredinibacter turnerae** | Yes | Yes | monotrichous | https://doi.org/10.1099/00207713-52-6-2261 |
| **Dasania marina (taxid:471499)** | Yes | No | monotrichous | https://pubmed.ncbi.nlm.nih.gov/18176532/ |
| **Acinetobacter baumanii** | No | No | non motile | https://doi.org/10.1007/978-1-4939-9118-1_17 |
| **Alkanindiges illinoisensis** | No | No | non motile | https://doi.org/10.1099/ijs.0.02568-0 |
| **Moraxella catarrhalis BBH18** | No | No | non motile | https://doi.org/10.1099/00221287-51-3-387 |
| **Perlucidibaca piscinae** | No | No | monotrichous | https://doi.org/10.1099/ijs.0.65039-0 |
| **Psychrobacter cryohalolentis** | No | No | non motile | https://doi.org/10.1099/ijs.0.64043-0 |
| **Thioalkalivibrio sp. K90 mix** | Yes | Yes | monotrichous | https://doi.org/10.1099/00207713-51-2-565 |
| **Allochromatium vinosum** | Yes | Yes | monotrichous | https://doi.org/10.1099/00207713-48-4-1129 |
| **Solimonas sp. K1W22B-7** | Yes | No | non motile | https://doi.org/10.1099/ijs.0.64938-0 |
| **Stenotrophomonas maltophilia R5513** | Yes | No | lophotrichous | https://doi.org/10.1099/00221287-26-1-123 |
| **Bordetella bronchiseptica RB50** | Yes | No | peritrichous | |
| **Burkholderia mallei ATCC 23344** | Yes | No | degenerate flagellum | |

| | | | | |
|---|---|---|---|---|
| **Burkholderia pseudomallei** | Yes | No | monotrichous | https://doi.org/10.1128/IAI.71.4.1622-1629.2003 |
| **Caulobacter crescentus** | No | No | monotrichous | |
| **Hyphomonas neptunium** | Yes | No | monotrichous | |
| **Magnetospirillum magneticum** | Yes | No | amphitrichous | https://doi.org/10.1099/00207713-31-4-452 |
| **Thalassospira xiamenensis** | Yes | No | monotrichous | https://doi.org/10.1099/ijs.0.64544-0 |
| **Rhodospirillum rubrum** | Yes | No | amphilophotrichous | |
| **Enhydrobacter aerosaccus** | No | No | non flagellated | https://doi.org/10.1099/00207713-37-3-289 |
| **Phycisphaera mikurensis** | Yes | No | monotrichous | https://doi.org/10.2323/jgam.55.267 |
| **Gimesia maris** | Yes | No | polar to subpolar | https://doi.org/10.1186/1944-3277-9-10 |
| **Planctopirus ephydatiae** | Yes | No | monotrichous | https://doi.org/10.1016/j.syapm.2019.126022 |
| **Rubinisphaera brasiliensis** | Yes | No | monotrichous | https://doi.org/10.1016/S0723-2020(89)80008-6 |
| **schlesneria paludicola** | Yes | No | subpolar | https://doi.org/10.1186/1944-3277-9-10 |
| **Thermogutta terrifontis** | Yes | No | monotrichous | https://doi.org/10.1099/ijs.0.000009 |
| **Brachyspira hyodysenteriae WA1** | Yes | No | spirochaete | |
| **Leptospira interrogans** | Yes | No | spirochaete | |
| **Borreliella bavariensis** | Yes | No | spirochaete | |
| **Spirochaeta africana** | Yes | No | spirochaete | https://doi.org/10.1099/00207713-46-1-305 |
| **Salinispira pacifica** | Yes | No | spirochaete | |
| **Oceanispirochaeta sp. K2** | Yes | No | spirochaete | https://doi.org/10.1099/ijsem.0.002130 |
| **Lentibacillus amyloliquefaciens** | No | No | non motile | https://doi.org/10.1007/s10482-015-0618-9 |

| | | | | |
|---|---|---|---|---|
| **Virgibacillus halodenitrificans** | Yes | No | polar and lateral | https://doi.org/10.1099/00207713-39-2-145 |
| **Bacillus subtilis** | Yes | No | peritrichous | |
| **Halobacillus halophilus** | Yes | No | peritrichous | https://doi.org/10.1016/s0723-2020(83)80007-1 |
| **Psychrobacillus sp. AK 1817** | Yes | No | peritrichous | https://doi.org/10.1128/jb.94.4.889-895.1967 |
| **Clostridioides difficile 630** | Yes | No | peritrichous | https://doi.org/10.1128/iai.69.12.7937-7940.2001 |
| **Selenomonas sputigena** | No | No | mid cell | https://doi.org/10.1128%2FMMBR.18.3.165-169.1954 |
| **Selenomonas ruminantium** | Yes | No | mid cell | https://doi.org/10.1128/aem.00286-11 |

#### Supplementary Table S3: Bacterial strains used in this study

##### *Escherichia coli*

| Strain | Genotype | Reference |
|---|---|---|
| ***SM10λpir*** | KmR, thi-1, thr, leu, tonA, lacY, supE, recA::RP4-2-Tc::Mu, pir | Simon et al, 1983 |
| ***DH5pir*** | *sup E44, ΔlacU169 (ΦlacZΔM15), recA1, endA1, hsdR17, thi-1, gyrA96, relA1, λpir phage lysogen* | Miller & Mekanalos, 1988 |
| ***BTH101*** | *F-, cya-99, araD139, galE15, galK16, rpsL1 (StrR), hsdR2, mcrA1, mcrB1, relA1* | Euromedex |
| **WM3064** | *thrB1004 pro thi rpsL hsdS lacZ ΔM15 RP4-1360 Δ(araBAD) 567ΔdapA 1341: [erm pir(wt)]* | W. Metcalf, University of Illinois, Urbana-Champaign |

##### *Vibrio parahaemolyticus*

| Strain | Genotype | Reference |
|---|---|---|
| **RIMD 2210633** | Clinical isolate, wild type | Makino et al, 2003 |
| **EP12** | *Δvp2234 (ΔflhF), Δvpa1548 (ΔlafA)* | Arroyo Pérez et al., 2021 |
| **SR58** | *Δvp2225 (ΔcheW)* | Ringgaard et al., 2014 |
| **JH2** | *Δvpa1548 (ΔlafA)* | Arroyo Pérez et al., 2021 |
| **EP15** | *Δvp2224 (ΔfipA), Δvpa1548 (ΔlafA)* | this study |
| **PM60** | *Δvp2234 (ΔflhF)* | Arroyo Pérez et al., 2021 |
| **SW01** | *Δvp2224 (ΔfipA)* | this study |
| **JH4** | *Δvp2191 (ΔhubP)* | Arroyo Pérez et al., 2021 |
| **PM69** | *Δvp2234::vp2234-sfgfp (ΔflhF::flhF-sfgfp)* | Arroyo Pérez et al., 2021 |
| **PM77** | *Δvp2234::vp2234-sfgfp (ΔflhF::flhF-sfgfp), Δvp2224 (fipA)* | this study |
| **EP11** | *Δvp2234::vp2234-sfgfp (ΔflhF::flhF-sfgfp), Δvp2191 (hubP)* | Arroyo Pérez et al., 2021 |
| **EP09** | *Δvp2234::vp2234-sfgfp (ΔflhF::flhF-sfgfp), Δvp2224 (fipA), Δvp2191 (hubP)* | this study |
| **PM65** | *vp2224 (fipA) L129A* | this study |
| **PM66** | *vp2224 (fipA) G110A* | this study |
| **EP16** | *Δvp2234::vp2234-sfgfp (ΔflhF::flhF-sfgfp), vp2224 (fipA) L129A* | this study |
| **EP17** | *Δvp2234::vp2234-sfgfp (ΔflhF::flhF-sfgfp), vp2224 (fipA) G110A* | this study |
| **EP13** | *vp2224 L129A, Δvpa1548(lafA)* | this study |
| **EP14** | *vp2224 G110A, Δvpa1548(lafA)* | this study |

| | | |
|---|---|---|
| **PM64** | *Δvp2224::vp2224-sfgfp (ΔfipA::fipA-sfgfp)* | this study |
| **PM68** | *Δvp2224::vp2224-sfgfp (ΔfipA::fipA-sfgfp), Δvp2234 (flhF)* | this study |
| **PM71** | *Δvp2224::vp2224 L129A-sfgfp (ΔfipA L129A ::fipA-sfgfp)* | this study |
| **PM72** | *Δvp2224::vp2224 G110A-sfgfp (ΔfipA G110A ::fipA-sfgfp)* | this study |

***Pseudomonas putida***

| **Strain** | **Genotype** | **Reference** |
|---|---|---|
| **Wild type** | Wild-type strain of *P. putida* KT2440 | Nelson et al, 2002 |
| **FliC S267C** | markerless in-frame substitution of Ser267 to Cys in the flagellin protein FliC (PP_4378) | Hintsche et al, 2017 |
| **FliC S267C Δ*flhF*** | markerless in-frame substitution of Ser267 to Cys in the flagellin protein FliC (PP_4378) and deletion of the gene *flhF* (*PP_4343*) | this study |
| **Δ*fipA*** | deletion of the gene *fipA* (*PP_4331*) | this study |
| **FliC S267C Δ*fipA*** | markerless in-frame substitution of Ser267 to Cys in the flagellin protein FliC (PP_4378) and deletion of the gene *fipA* (*PP_4331*) | this study |
| **FipA -DILEL-sfGFP** | C-terminal sfGFP tag of FipA (PP_4331) linked with Asp, Ile, Leu, Glu and Leu | this study |
| **FliC S267C Δ*flhF* FipA-DILEL-sfGFP** | markerless in-frame substitution of Ser267 to Cys in the flagellin protein FliC (PP_4378), deletion of the gene *flhF* (*PP_4343*) and C-terminal sfGFP tag of FipA (PP_4331) linked with Asp, Ile, Leu, Glu and Leu | this study |
| **FlhF-GS-mCherry** | C-terminal mCherry tag of FlhF (PP_4343) linked with Gly and Ser | this study |
| **FliC S267C Δ*fimV* FipA -DILEL-sfGFP** | markerless in-frame substitution of Ser267 to Cys in the flagellin protein FliC (PP_4378), deletion of the gene *fimV* (*PP_1992*) and C-terminal sfGFP tag of FipA (PP_4331) linked with Asp, Ile, Leu, Glu and Leu | this study |
| **ΔfipA FlhF-GS-mCherry** | deletion of the gene *fipA* (*PP_4331*) and C-terminal mCherry tag of FlhF (PP_4343) linked with Gly and Ser | this study |
| **FliC S267C FipA G104A** | markerless in-frame substitution of Ser267 to Cys in the flagellin protein FliC (PP_4378) and in-frame substitution of Gly104 to Ala in the protein FipA (Sputcn32_4331) | this study |
| **FipA L123A** | markerless in-frame substitution of Leu123 to Ala in the protein FipA (Sputcn32_4331) | this study |
| **FliC S267C FipA L123A** | markerless in-frame substitution of Ser267 to Cys in the flagellin protein FliC (PP_4378) and in-frame substitution of Leu123 to Ala in the protein FipA (Sputcn32_4331) | this study |
| **FliC S267C FipA-DILEL-sfGFP** | markerless in-frame substitution of Ser267 to Cys in the flagellin protein FliC (PP_4378) and C-terminal sfGFP tag of FipA (PP_4331) linked with Asp, Ile, Leu, Glu and Leu | this study |
| **FliC S267C FipA L116A** | markerless in-frame substitution of Ser267 to Cys in the flagellin protein FliC (PP_4378) and in-frame substitution of Leu116 to Ala in the protein FipA (Sputcn32_4331) | this study |
| **FliC S267C FipA G104A-DILEL-sfGFP** | markerless in-frame substitution of Ser267 to Cys in the flagellin protein FliC (PP_4378) and in-frame substitution of Gly104 to Ala in the protein FipA (Sputcn32_4331) with C-terminal sfGFP tag linked | this study |

| | | |
|---|---|---|
| | with Asp, Ile, Leu, Glu and Leu | |
| **FliC S267C FipA L123A-DILEL-sfGFP** | markerless in-frame substitution of Ser267 to Cys in the flagellin protein FliC (PP_4378) and in-frame substitution of Leu123 to Ala in the protein FipA (Sputcn32_4331) with C-terminal sfGFP tag linked with Asp, Ile, Leu, Glu and Leu | this study |
| **FliC S267C FipA L116A-DILEL-sfGFP** | markerless in-frame substitution of Ser267 to Cys in the flagellin protein FliC (PP_4378) and in-frame substitution of Leu116 to Ala in the protein FipA (Sputcn32_4331) with C-terminal sfGFP tag linked with Asp, Ile, Leu, Glu and Leu | this study |
| **FlhF-GS-mCherry ΔfimV** | C-terminal mCherry tag of FlhF (PP_4343) linked with Gly and Ser and deletion of the gene *fimV* (*PP_1992*) | this study |
| **FliC S267C ΔfipA FipA KI** | markerless in-frame substitution of Ser267 to Cys in the flagellin protein FliC (PP_4378), deletion of the gene *fipA* (*Sputcn32_4331*) and reconstitution of the gene *fipA* (*Sputcn32_4331*) | this study |
| **FlhF-GS-mCherry ΔfimV ΔfipA** | C-terminal mCherry tag of FlhF (PP_4343) linked with Gly and Ser and deletion of the genes *fimV* (*PP_1992*) and *fipA* (*Sputcn32_4331*) | this study |
| **FlhF-GS-mCherry FipA L123A** | C-terminal mCherry tag of FlhF (PP_4343) linked with Gly and Ser and markerless in-frame substitution of Leu123 to Ala in the protein FipA (Sputcn32_4331) | this study |
| **FlhF-GS-mCherry FipA L116A** | C-terminal mCherry tag of FlhF (PP_4343) linked with Gly and Ser and markerless in-frame substitution of Leu116 to Ala in the protein FipA (Sputcn32_4331) | this study |
| **FlhF-GS-mCherry FipA G104A** | C-terminal mCherry tag of FlhF (PP_4343) linked with Gly and Ser and markerless in-frame substitution of Gly104 to Ala in the protein FipA (Sputcn32_4331) | this study |
| **FliC S267C FlhF D362A-GS-mCherry** | markerless in-frame substitution of Ser267 to Cys in the flagellin protein FliC (PP_4378) in-frame substitution of Asp362 to Ala in FlhF (PP_4343) with C-terminal mCherry tag linked with Gly and Ser | this study |
| **FliC S267C FlhF-GS-mCherry FipA ΔTMD** | markerless in-frame substitution of Ser267 to Cys in the flagellin protein FliC (PP_4378) and in-frame deletion of the N-terminal transmembrane domain in the protein FipA (Sputcn32_4331) | this study |
| **FliC S267C FlhF-GS-mCherry FipA ΔTMD** | markerless in-frame substitution of Ser267 to Cys in the flagellin protein FliC (PP_4378), C-terminal mCherry tag of FlhF (PP_4343) linked with Gly and Ser and in-frame deletion of the N-terminal transmembrane domain in the protein FipA (Sputcn32_4331) | this study |
| **FliC S267C FipA ΔTMD-DILEL-sfGFP** | markerless in-frame substitution of Ser267 to Cys in the flagellin protein FliC (PP_4378) and in-frame deletion of the N-terminal transmembrane domain in the protein FipA (Sputcn32_4331) with C-terminal sfGFP tag linked with Asp, Ile, Leu, Glu and Leu | this study |

***Shewanella putrefaciens***

| **Strain** | **Genotype** | **Reference** |
|---|---|---|
| **wild type** | wild type strain of *S. putrefaciens* CN-32 | Fredrickson et al, 1998 |
| ***ΔflhF*** | deletion of the gene *flhF* (*Sputcn32_2561*) | Rossmann et |

| | | |
|---|---|---|
| | | al, 2015 |
| **CheA-mCherry** | C-terminal mCherry tag of CheA (Sputcn32_2556) | this study |
| ***$flgE_1$* T183C** | markerless in-frame substitution of Thr183 to Cys in the polar hook protein $FlgE_1$ (Sputcn32_2594), fully functional and suitable for maleimide staining | Rossmann et al, 2019 |
| ***$flaB_1$* T166C *$flaA_1$* T174C Δ*flagL*** | markerless in-frame substitution of Thr166 to Cys in the polar major flagellin protein $FlaB_1$ (Sputcn32_2585) and Thr174 to Cys in the polar minor flagellin protein $FlaA_1$ (Sputcn32_2586), fully functional and suitable for maleimide staining and deletion of the lateral gene cluster (Sputcn32_*3444*-Sputcn32_*3485*) | Kühn et al, 2017 |
| **Δ*fipA*** | deletion of the gene *fipA* (*Sputcn32_2550*) | this study |
| **CheA-mCherry Δ*fipA*** | C-terminal mCherry tag of CheA (Sputcn32_2556) and deletion of the gene *fipA* (*Sputcn32_2550*) | this study |
| ***$flaB_1$* T166C *$flaA_1$* T174C Δ*flagL* Δ*fipA*** | markerless in-frame substitution of Thr166 to Cys in the polar major flagellin protein $FlaB_1$ (Sputcn32_2585) and Thr174 to Cys in the polar minor flagellin protein $FlaA_1$ (Sputcn32_2586), fully functional and suitable for maleimide staining, deletion of the lateral gene cluster (Sputcn32_*3444*-Sputcn32_*3485*) and deletion of the gene *fipA* (*Sputcn32_2550*) | this study |
| **FlhF-GS-mVenus** | C-terminal mVenus tag of FlhF (Sputcn32_2561) linked with Gly and Ser | this study |
| **FipA-DILEL-sfGFP** | C-terminal sfGFP tag of FipA (Sputcn32_2550) linked with Asp, Ile, Leu, Glu and Leu | this study |
| **FlhF-GS-mVenus Δ*fipA*** | C-terminal mVenus tag of FlhF (Sputcn32_2561) linked with Gly and Ser and deletion of the gene *fipA* (*Sputcn32_2550*) | this study |
| **FlhF-GS-mVenus Δ*flhG*** | C-terminal mVenus tag of FlhF (Sputcn32_2561) linked with Gly and Ser and deletion of the gene *flhG* (*Sputcn32_2560*) | this study |
| **FipA-DILEL-sfGFP Δ*flhF*** | C-terminal sfGFP tag of FipA (Sputcn32_2550) linked with Asp, Ile, Leu, Glu and Leu and deletion of the gene *flhF* (*Sputcn32_2561*) | this study |
| **FlhF-GS-mVenus Δ*hubP*** | C-terminal mVenus tag of FlhF (Sputcn32_2561) linked with Gly and Ser and deletion of the gene *hubP* (*Sputcn32_2442*) | this study |
| **FlhF-GS-mVenus Δ*fipA* Δ*hubP*** | C-terminal mVenus tag of FlhF (Sputcn32_2561) linked with Gly and Ser and deletion of the genes *fipA* (*Sputcn32_2550*) and *hubP* (*Sputcn32_2442*) | this study |
| ***$flgE_1$* T183C Δ*flagL*** | markerless in-frame substitution of Thr183 to Cys in the polar hook protein $FlgE_1$ (Sputcn32_2594), fully functional and suitable for maleimide staining and deletion of the lateral gene cluster (Sputcn32_*3444-Sputcn32_3485*) | Hook et al, 2020 |
| **FipA-DILEL-sfGFP Δ*hubP*** | C-terminal sfGFP tag of FipA (Sputcn32_2550) linked with Asp, Ile, Leu, Glu and Leu and deletion of the gene *hubP* (*Sputcn32_2442*) | this study |
| **HubP-mCherry FlhF-GS-Venus** | C-terminal mCherry tag of HubP (Sputcn32_2442) and C-terminal mVenus tag of FlhF (Sputcn32_2561) linked with Gly and Ser | this study |
| **FlhF-GS-mVenus Δ*hubP*** | C-terminal mVenus tag of FlhF (Sputcn32_2561) linked with Gly and Ser and deletion of the genes hubP (*Sputcn32_2442*) and *Sputcn32_3157* | this study |

| | | |
|---|---|---|
| ***ΔSputcn32_3157*** | | |
| ***$flaB_1$* T166C *$flaA_1$* T174C *ΔflagL* FipA-DILEL-sfGFP** | markerless in-frame substitution of Thr166 to Cys in the polar major flagellin protein $FlaB_1$ (Sputcn32_2585) and Thr174 to Cys in the polar minor flagellin protein $FlaA_1$ (Sputcn32_2586), fully functional and suitable for maleimide staining, deletion of the lateral gene cluster (Sputcn32_*3444*-Sputcn32_*3485*) and C-terminal sfGFP tag of FipA (Sputcn32_2550) linked with Asp, Ile, Leu, Glu and Leu | this study |
| ***$flaB_1$* T166C *$flaA_1$* T174C *ΔflagL ΔflhF*** | markerless in-frame substitution of Thr166 to Cys in the polar major flagellin protein $FlaB_1$ (Sputcn32_2585) and Thr174 to Cys in the polar minor flagellin protein $FlaA_1$ (Sputcn32_2586), fully functional and suitable for maleimide staining, deletion of the lateral gene cluster (Sputcn32_*3444*-Sputcn32_*3485*) and deletion of the gene *flhF* (*Sputcn32_2561*) | this study |
| ***ΔfipA fipA* (*Sputcn32_2550*) KI** | deletion of the gene *fipA* (*Sputcn32_2550*) and reconstitution of the gene *fipA* (*Sputcn32_2550*) | this study |
| **FipA L118A** | markerless in-frame substitution of Leu118 to Ala in the protein FipA (Sputcn32_2550) | this study |
| **FipA G106A** | markerless in-frame substitution of Gly106 to Ala in the protein FipA (Sputcn32_2550) | this study |
| **FipA L118-DILEL-sfGFP** | markerless in-frame substitution of Leu118 to Ala in the protein FipA (Sputcn32_2550) with a C-terminal sfGFP tag linked with Asp, Ile, Leu, Glu and Leu | this study |
| **FipA G106A-DILEL-sfGFP** | markerless in-frame substitution of Gly106 to Ala in the protein FipA (Sputcn32_2550) with a C-terminal sfGFP tag linked with Asp, Ile, Leu, Glu and Leu | this study |
| ***$flgE_1$* T183C *ΔflagL* $FliM_1$-GS-sfGFP** | markerless in-frame substitution of Thr183 to Cys in the polar hook protein $FlgE_1$ (Sputcn32_2594), fully functional and suitable for maleimide staining, deletion of the lateral gene cluster (Sputcn32_*3444-Sputcn32_3485*) and C-terminal sfGFP tag of $FliM_1$ (Sputcn32_2569) linked with Gly and Ser | Hook et al, 2020 |
| **FipA L125A** | markerless in-frame substitution of Leu125 to Ala in the protein FipA (Sputcn32_2550) | this study |
| ***$flaB_1$* T166C *$flaA_1$* T174C *ΔflagL ΔfipA ΔhubP*** | markerless in-frame substitution of Thr166 to Cys in the polar major flagellin protein $FlaB_1$ (Sputcn32_2585) and Thr174 to Cys in the polar minor flagellin protein $FlaA_1$ (Sputcn32_2586), fully functional and suitable for maleimide staining and deletion of the lateral gene cluster (Sputcn32_*3444*-Sputcn32_*3485*) and the genes *fipA* (*Sputcn32_2550*) and *hubP* (*Sputcn32_2442*) | this study |
| **FipA L125A-DILEL-sfGFP** | markerless in-frame substitution of Leu125 to Ala in the protein FipA (Sputcn32_2550) with a C-terminal sfGFP tag linked with Asp, Ile, Leu, Glu and Leu | this study |
| **FlhF-mCherry FipA-DILEL-sfGFP** | C-terminal mCherry tag of FlhF (Sputcn32_2561) and C-terminal sfGFP tag of FipA (Sputcn32_2550) linked with Asp, Ile, Leu, Glu and Leu | this study |
| **FipA L118A FlhF-GS-mVenus** | markerless in-frame substitution of Leu118 to Ala in the protein FipA (Sputcn32_2550) and C-terminal mVenus tag of FlhF (Sputcn32_2561) linked with Gly and Ser | this study |
| **FipA G106A** | markerless in-frame substitution of Gly106 to Ala in the protein | this study |

| | | |
|---|---|---|
| **FlhF-GS-mVenus** | FipA (Sputcn32_2550) and C-terminal mVenus tag of FlhF (Sputcn32_2561) linked with Gly and Ser | |
| **FipA L125A FlhF-GS-mVenus** | markerless in-frame substitution of Leu125 to Ala in the protein FipA (Sputcn32_2550) and C-terminal mVenus tag of FlhF (Sputcn32_2561) linked with Gly and Ser | this study |
| **FipA ΔTMD** | in-frame deletion of the N-terminal transmembrane domain in the protein FipA (Sputcn32_2550) | this study |
| **FlhF-GS-mVenus FipA ΔTMD** | markerless in-frame deletion of the N-terminal transmembrane domain in the protein FipA (Sputcn32_2550) and C-terminal mVenus tag of FlhF (Sputcn32_2561) linked with Gly and Ser | this study |
| **FipA ΔTMD-DILEL-sfGFP** | in-frame deletion of the N-terminal transmembrane domain in the protein FipA (Sputcn32_2550) and C-terminal sfGFP tag of FipA (Sputcn32_2550) linked with Asp, Ile, Leu, Glu and Leu | this study |

#### Additional References for the Strains

Arroyo-Pérez EE, Ringgaard S. Interdependent Polar Localization of FlhF and FlhG and Their Importance for Flagellum Formation of *Vibrio parahaemolyticus*. Front Microbiol. 2021 Mar 17;12:655239. doi: 10.3389/fmicb.2021.655239. PMID: 33815347; PMCID: PMC8009987.

Fredrickson, J. K. et al. Biogenic iron mineralization accompanying the dissimilatory reduction of hydrous ferric oxide by a groundwater bacterium. Geochim. Cosmochim. Acta 62, 3239–3257 (1998)

Hintsche M, Waljor V, Großmann R, Kühn MJ, Thormann KM, et al. A polar bundle of flagella can drive bacterial swimming by pushing, pulling, or coiling around the cell body. Sci Rep. 2017 Dec 1;7(1):16771. doi: 10.1038/s41598-017-16428-9. Erratum in: Sci Rep. 2022 Aug 12;12(1):13732. PMID: 29196650; PMCID: PMC5711944.

Hook JC, Blagotinsek V, Pané-Farré J, Mrusek D, Altegoer F, et al. A Proline-Rich Element in the Type III Secretion Protein FlhB Contributes to Flagellar Biogenesis in the Beta- and Gamma-Proteobacteria. Front Microbiol. 2020 Dec 15;11:564161. doi: 10.3389/fmicb.2020.564161. PMID: 33384667; PMCID: PMC7771051.

Kühn MJ, Schmidt FK, Eckhardt B, Thormann KM. Bacteria exploit a polymorphic instability of the flagellar filament to escape from traps. Proc Natl Acad Sci U S A. 2017 Jun 13;114(24):6340-6345. doi: 10.1073/pnas.1701644114. Epub 2017 May 30. PMID: 28559324; PMCID: PMC5474801.

Makino K, Oshima K, Kurokawa K, Yokoyama K, Uda T, et al. Genome sequence of *Vibrio parahaemolyticus*: a pathogenic mechanism distinct from that of *V cholerae*. Lancet. 2003 Mar 1;361(9359):743-9. doi: 10.1016/S0140-6736(03)12659-1. PMID: 12620739.

Miller, V. L. & Mekalanos, J. J. (1988) A novel suicide vector and its use in construction of insertion mutations: Osmoregulation of outer membrane proteins and virulence determinants in *Vibrio cholera* requires *toxR*. J. Bacteriol.170, 2575–2583

Nelson KE, Weinel C, Paulsen IT, Dodson RJ, Hilbert H, et al. Complete genome sequence and comparative analysis of the metabolically versatile *Pseudomonas putida* KT2440. Environ Microbiol. 2002 Dec;4(12):799-808. doi: 10.1046/j.1462-2920.2002.00366.x. Erratum in: Environ Microbiol. 2003 Jul;5(7):630. PMID: 12534463.

Ringgaard S, Zepeda-Rivera M, Wu X, Schirner K, Davis BM, Waldor MK. ParP prevents dissociation of CheA from chemotactic signaling arrays and tethers them to a polar anchor. Proc Natl Acad Sci U S A. 2014 Jan 14;111(2):E255-64. doi: 10.1073/pnas.1315722111. Epub 2013 Dec 30. PMID: 24379357; PMCID: PMC3896188.

Rossmann, F. et al. The role of FlhF and HubP as polar landmark proteins in *Shewanella putrefaciens* CN-32. Mol. Microbiol. 98, 727–742 (2015)

Rossmann, F.M. et al. The GGDEF domain of the phosphodiesterase PdeB in *Shewanella putrefaciens* mediates recruitment by the polar landmark protein HubP. J Bacteriol. 201, 7 e00534-18. (2019)

Simon R, Priefer U, Pühler A (1983) A broad host range mobilization system for *in vivo* genetic-engineering - transposon mutagenesis in Gram-negative bacteria. Bio-Technol 1: 784–791

#### Supplementary Table S4: Plasmids used in this study

##### General constructions

| Plasmid | Description | Reference |
|---|---|---|
| **pDM4** | Suicide vector for gene deletions in *Vibrio* sp. | Milton et al., 1996 |
| **pJH036** | pBAD33 derivative for sfGFP C-terminal fusion | Iyer et al, 2020 |
| **pNPTS138-R6KT** | mobRP4+ ori-R6K *sacB;* β-galactosidase fragment alpha; suicide vector for in-frame deletions or integrations in *P. putida* and *S. putrefaciens*; $Kan^r$ | Lassak et al, 2010 |
| **pKT25** | *plac* ori p15A vector for protein-protein interaction analysis; MCS downstream from T25 fragment encoding region; $Kan^r$ | Karimova et al, 1998 |
| **pKNT25** | *plac* ori p15A vector for protein-protein interaction analysis; MCS upstream from T25 fragment encoding region; $Kan^r$ | Karimova et al, 1998 |
| **pUT18** | *plac* ori Col E1 vector for protein-protein interaction analysis; MCS downstream from T18 fragment encoding region; $Amp^r$ | Karimova et al, 1998 |
| **pUT18C** | *plac* ori Col E1 vector for protein-protein interaction analysis; MCS upstream from T18 fragment encoding region; $Amp^r$ | Karimova et al, 1998 |

##### Plasmids for *Vibrio*

| Plasmids | Description | Reference |
|---|---|---|
| **pJH003** | For deletion of vpa1548(lafA) | Heering & Ringgaard, 2016 |
| **pSW022** | For deletion of vp2224(fipA) | This work |
| **pPM188fip** | For insertion of vp2234-sfgfp (flhF-sfgfp), replacing native flhF | Arroyo Pérez et al., 2021 |
| **pPM178** | For insertion of vp2224-sfgfp (fipA-sfgfp), replacing native fipA | this work |
| **pPM179** | For insertion of vp2224 (fipA) G110A point mutation in the chromosome in the native locus | this work |
| **pPM180** | For insertion of vp2224 (fipA) L129A point mutation in the chromosome in the native locus | this work |
| **pPM191** | For insertion of vp2224 (fipA) G110A fused to sfGFP in the chromosome in the native locus | this work |
| **pPM187** | For insertion of vp2224 (fipA) L129A fused to sfGFP in the chromosome in the native locus | this work |
| **pPM039** | For deletion of vp2191 (hubP) | Arroyo Pérez et al., 2021 |
| **pPM194** | For overexpression of VP2224(FipA)Δ7-27 -sfGFP | this work |
| **pPM146** | For overexpression of VP2224(FipA) | this work |
| **pPM159** | For overexpression of VP2224(FipA)-sfGFP | this work |

##### Plasmids for *Pseudomonas putida*

| Plasmid | Description | Reference |
|---|---|---|
| **pNPTS138-R6KT *flhF* KO (*PP_4343*)** | plasmid for deletion of the *flhF* gene (*PP_4343*) in *P. putida* KT2440; Kan$^r$ | this study |
| **pNPTS138-R6KT FlhF K235A (PP_4343)** | plasmid for in frame complementation of *flhF* (*PP_4343*) with FlhF K235A mutant in *P. putida* KT2440; Kan$^r$ | this study |
| **pNPTS138-R6KT FlhF-GS-mCherry (PP_4343)** | plasmid for in frame complementation of *flhF* (*PP_4343*) with FlhF-GS-mCherry in *P. putida* KT2440; Kan$^r$ | this study |
| **pNPTS138-R6KT FlhF K235A-GS-mCherry (PP_4343)** | plasmid for in frame complementation of *flhF* (*PP_4343*) with FlhF K235A-GS-mCherry mutant in *P. putida* KT2440; Kan$^r$ | this study |
| **pNPTS138-R6KT FlhF D301A-GS-mCherry (PP_4343)** | plasmid for in frame complementation of *flhF* (*PP_4343*) with FlhF D301A-GS-mCherry mutant in *P. putida* KT2440; Kan$^r$ | this study |
| **pNPTS138-R6KT FlhF D362A-GS-mCherry (PP_4343)** | plasmid for in frame complementation of *flhF* (*PP_4343*) with FlhF D362A-GS-mCherry mutant in *P. putida* KT2440; Kan$^r$ | this study |
| **pNPTS138-R6KT *fipA* KO (*PP_4331*)** | plasmid for deletion of the *fipA* gene (*PP_4331*) in *P. putida* KT2440; Kan$^r$ | this study |
| **pNPTS138-R6KT *fipA* KI (*PP_4331*)** | plasmid for in frame complementation of *fipA* (*PP_4331*) with wild type *fipA* in *P. putida* KT2440; Kan$^r$ | this study |
| **pNPTS138-R6KT FipA ΔTMD (AS5-22) (PP_4331)** | plasmid for in frame complementation of *fipA* (*PP_4331*) with FipA ΔTMD mutant in *P. putida* KT2440; Kan$^r$ | this study |
| **pNPTS138-R6KT FipA G104A (PP_4331)** | plasmid for in frame complementation of *fipA* (*PP_4331*) with FipA G104A mutant in *P. putida* KT2440; Kan$^r$ | this study |
| **pNPTS138-R6KT FipA L116A (PP_4331)** | plasmid for in frame complementation of *fipA* (*PP_4331*) with FipA L116A mutant in *P. putida* KT2440; Kan$^r$ | this study |
| **pNPTS138-R6KT FipA L123A (PP_4331)** | plasmid for in frame complementation of *fipA* (*PP_4331*) with FipA L123A mutant in *P. putida* KT2440; Kan$^r$ | this study |
| **pNPTS138-R6KT FipA-DILEL-sfGFP (PP_4331)** | plasmid for in frame complementation of *fipA* (*PP_4331*) with FipA-DILEL-sfGFP in *P. putida* KT2440; Kan$^r$ | this study |
| **pNPTS138-R6KT FipA ΔTMD-DILEL-sfGFP (AS5-22) (PP_4331)** | plasmid for in frame complementation of *fipA* (*PP_4331*) with FipA ΔTMD-DILEL-sfGFP mutant in *P. putida* KT2440; Kan$^r$ | this study |

| | | |
|---|---|---|
| **pNPTS138-R6KT FipA G104A-DILEL-sfGFP (PP_4331)** | plasmid for in frame complementation of *fipA* (*PP_4331*) with FipA G104A-DILEL-sfGFP mutant in *P. putida* KT2440; Kan$^r$ | this study |
| **pNPTS138-R6KT FipA L116A-DILEL-sfGFP (PP_4331)** | plasmid for in frame complementation of *fipA* (*PP_4331*) with FipA L116A-DILEL-sfGFP mutant in *P. putida* KT2440; Kan$^r$ | this study |
| **pNPTS138-R6KT FipA L123A-DILEL-sfGFP (PP_4331)** | plasmid for in frame complementation of *fipA* (*PP_4331*) with FipA L123A-DILEL-sfGFP mutant in *P. putida* KT2440; Kan$^r$ | this study |

**Plasmids for *Shewanella putrefaciens***

| **Plasmid** | **Description** | **Reference** |
|---|---|---|
| **pNPTS138-R6KT polar flagellar cluster KO (*Sputcn32_2548-2608*)** | plasmid for deletion of the polar flagellar gene cluster (*Sputcn32_2548-2608*) in *S. putrefaciens* CN-32; Kan$^r$ | this study |
| **pNPTS138-R6KT lateral flagellar cluster KO (*Sputcn32_3444-3485*)** | plasmid for deletion of the lateral flagellar gene cluster (*Sputcn32_3444-3485*) in *S. putrefaciens* CN-32; Kan$^r$ | Lassak et al, 2010 |
| **pNPTS138-R6KT *flagL* KO (*Sputcn32_3455*, *Sputcn32_3456*)** | plasmid for deletion of the lateral flagellin genes (*Sputcn32_3455*, *Sputcn32_3456*) in *S. putrefaciens* CN-32; Kan$^r$ | Rossmann et al, 2015 |
| **pNPTS138-R6KT *hubP* KO (*Sputcn32_2442*)** | plasmid for deletion of the *hubP* gene (*Sputcn32_2442*) in *S. putrefaciens* CN-32; Kan$^r$ | Rossmann et al, 2015 |
| **pNPTS138-R6KT *flhF* KO (*Sputcn32_2561*)** | plasmid for deletion of the *flhF* gene (*Sputcn32_2561*) in *S. putrefaciens* CN-32; Kan$^r$ | Rossmann et al, 2015 |
| **pNPTS138-R6KT *flhG* KO (*Sputcn32_2560*)** | plasmid for deletion of the *flhG* gene (*Sputcn32_2560*) in *S. putrefaciens* CN-32; Kan$^r$ | Schuhmacher et al, 2015 |
| **pNPTS138-R6KT FlhF-GS-Venus (Sputcn32_2561)** | plasmid for in frame complementation of *flhF* (*Sputcn32_2561*) with FlhF-GS-mVenus in *S. putrefaciens* CN-32; Kan$^r$ | this study |
| **pNPTS138-R6KT *fipA* KO (*Sputcn32_2550*)** | plasmid for deletion of the *fipA* gene (*Sputcn32_2550*) in *S. putrefaciens* CN-32; Kan$^r$ | this study |
| **pNPTS138-R6KT *fipA* KI (*Sputcn32_2550*)** | plasmid for in frame complementation of *fipA* (*Sputcn32_2550*) with wild type *fipA* in *S. putrefaciens* CN-32; Kan$^r$ | this study |
| **pNPTS138-R6KT FipA ΔTMD (AS5-23) (Sputcn32_2550)** | plasmid for in frame complementation of *fipA* (*Sputcn32_2550*) with FipA ΔTMD mutant in *S. putrefaciens* CN-32; Kan$^r$ | this study |
| **pNPTS138-R6KT FipA G106A (Sputcn32_2550)** | plasmid for in frame complementation of *fipA* (*Sputcn32_2550*) with FipA G106A mutant in *S. putrefaciens* CN-32; Kan$^r$ | this study |
| **pNPTS138-R6KT FipA L118A (Sputcn32_2550)** | plasmid for in frame complementation of *fipA* (*Sputcn32_2550*) with FipA L118A mutant in *S. putrefaciens* CN-32; Kan$^r$ | this study |

| | | |
|---|---|---|
| **pNPTS138-R6KT FipA L125A (Sputcn32_2550)** | plasmid for in frame complementation of *fipA* (*Sputcn32_2550*) with FipA L125 mutant in *S. putrefaciens* CN-32; Kan$^r$ | this study |
| **pNPTS138-R6KT FipA-DILEL-sfGFP (Sputcn32_2550)** | plasmid for in frame complementation of *fipA* (*Sputcn32_2550*) with FipA-DILEL-sfGFP in *S. putrefaciens* CN-32; Kan$^r$ | this study |
| **pNPTS138-R6KT FipA ΔTMD-DILEL-sfGFP (AS5-23) (Sputcn32_2550)** | plasmid for in frame complementation of *fipA* (*Sputcn32_2550*) with FipA ΔTMD-DILEL-sfGFP mutant in *S. putrefaciens* CN-32; Kan$^r$ | this study |
| **pNPTS138-R6KT FipA G106A-DILEL-sfGFP (Sputcn32_2550)** | plasmid for in frame complementation of *fipA* (*Sputcn32_2550*) with FipA G106A-DILEL-sfGFP mutant in *S. putrefaciens* CN-32; Kan$^r$ | this study |
| **pNPTS138-R6KT FipA L116A-DILEL-sfGFP (Sputcn32_2550)** | plasmid for in frame complementation of *fipA* (*Sputcn32_2550*) with FipA L116A-DILEL-sfGFP mutant in *S. putrefaciens* CN-32; Kan$^r$ | this study |
| **pNPTS138-R6KT FipA L125A-DILEL-sfGFP (Sputcn32_2550)** | plasmid for in frame complementation of *fipA* (*Sputcn32_2550*) with FipA L125A mutant in *S. putrefaciens* CN-32; Kan$^r$ | this study |
| **pNPTS138-R6KT FliM$_1$-GS-sfGFP (Sputcn32_2569)** | plasmid for in frame complementation of *fliM$_1$* (*Sputcn32_2569*) with FliM$_1$-GS-sfGFP in *S. putrefaciens* CN-32; Kan$^r$ | Hook et al, (2020) |

**BACTH plasmids for *V. parahaemolyticus***

| **Plasmid** | **Description** | **Reference** |
|---|---|---|
| **pSW74** | T25-vp2224(fipA)Δ1-27 | this study |
| **pSW119** | T18-vp2224(fipA)Δ1-27 | this study |
| **pPM118** | vp2224(fipA)Δ1-27-T18 | this study |
| **pPM119** | vp2224(fipA)Δ1-27-T25 | this study |
| **pPM124** | vp2234(flhF)-T18 | this study |
| **pPM128** | vp2234(flhF)-T25 | this study |
| **pPM132** | T18-vp2234(flhF) | this study |
| **pPM136** | T25-vp2234(flhF) | this study |
| **pPM160** | T18-vp2224(fipA)Δ1-27 G110A | this study |
| **pPM161** | T18-vp2224(fipA)Δ1-27 E126A | this study |
| pPM162 | T18-vp2224(fipA)Δ1-27 L129A | this study |

**BACTH plasmids for *P. putida***

| Plasmid | Description | reference |
|---|---|---|
| **pKT25 FlhF (PP_4343)** | plasmid for BACTH assay carrying T25-FlhF (PP_4343); $Kan^r$ | this study |
| **pKNT25 FlhF (PP_4343)** | plasmid for BACTH assay carrying FlhF-T25 (PP_4343); $Kan^r$ | this study |
| **pUT18 FlhF (PP_4343)** | plasmid for BACTH assay carrying FlhF-T18 (PP_4343); $Amp^r$ | this study |
| **pUT18C FlhF (PP_4343)** | plasmid for BACTH assay carrying T18-FlhF (PP_4343); $Amp^r$ | this study |
| **pKT25 FlhF K235A (PP_4343)** | plasmid for BACTH assay carrying T25-FlhF K235A (PP_4343); $Kan^r$ | this study |
| **pKNT25 FlhF K235A (PP_4343)** | plasmid for BACTH assay carrying FlhF K235A - T25 (PP_4343); $Kan^r$ | this study |
| **pUT18 FlhF K235A (PP_4343)** | plasmid for BACTH assay carrying FlhF K235A - T18 (PP_4343); $Amp^r$ | this study |
| **pUT18C FlhF K235A (PP_4343)** | plasmid for BACTH assay carrying T18-FlhF K235A (PP_4343); $Amp^r$ | this study |
| **pKT25 FipA (PP_4331)** | plasmid for BACTH assay carrying T25-FipA (PP_4331); $Kan^r$ | this study |
| **pKNT25 FipA (PP_4331)** | plasmid for BACTH assay carrying FipA-T25 (PP_4331); $Kan^r$ | this study |
| **pUT18 FipA (PP_4331)** | plasmid for BACTH assay carrying FipA-T18 (PP_4331); $Amp^r$ | this study |
| **pUT18C FipA (PP_4331)** | plasmid for BACTH assay carrying T18-FipA (PP_4331); $Amp^r$ | this study |
| **pKT25 FipA G104A (PP_4331)** | plasmid for BACTH assay carrying T25-FipA G104A (PP_4331); $Kan^r$ | this study |
| **pKNT25 FipA G104A (PP_4331)** | plasmid for BACTH assay carrying FipA G104A - T25 (PP_4331); $Kan^r$ | this study |
| **pUT18 FipA G104A (PP_4331)** | plasmid for BACTH assay carrying FipA G104A - T18 (PP_4331); $Amp^r$ | this study |
| **pUT18C FipA G104A (PP_4331)** | plasmid for BACTH assay carrying T18-FipA G104A (PP_4331); $Amp^r$ | this study |
| **pKT25 FipA L116A (PP_4331)** | plasmid for BACTH assay carrying T25-FipA L116A (PP_4331); $Kan^r$ | this study |
| **pKNT25 FipA L116A (PP_4331)** | plasmid for BACTH assay carrying FipA L116A - T25 (PP_4331); $Kan^r$ | this study |
| **pUT18 FipA L116A (PP_4331)** | plasmid for BACTH assay carrying FipA L116A - T18 (PP_4331); $Amp^r$ | this study |
| **pUT18C FipA L116A (PP_4331)** | plasmid for BACTH assay carrying T18-FipA L116A (PP_4331); $Amp^r$ | this study |
| **pKT25 FipA L125A (PP_4331)** | plasmid for BACTH assay carrying T25-FipA L123A (PP_4331); $Kan^r$ | this study |

| | | |
|---|---|---|
| **pKNT25 FipA L125A (PP_4331)** | plasmid for BACTH assay carrying FipA L123A - T25 (PP_4331); Kan$^r$ | this study |
| **pUT18 FipA L125A (PP_4331)** | plasmid for BACTH assay carrying FipA L123A - T18 (PP_4331); Amp$^r$ | this study |
| **pUT18C FipA L125A (PP_4331)** | plasmid for BACTH assay carrying T18-FipA L123A (PP_4331); Amp$^r$ | this study |

**BACTH plasmids for *S. putrefaciens***

| **Plasmid** | **purpose/description** | **reference** |
|---|---|---|
| **pKT25 FlhF (Sputcn32_2561)** | plasmid for BACTH assay carrying T25-FlhF (Sputcn32_2561); Kan$^r$ | this study |
| **pKNT25 FlhF (Sputcn32_2561)** | plasmid for BACTH assay carrying FlhF-T25 (Sputcn32_2561); Kan$^r$ | this study |
| **pUT18 FlhF (Sputcn32_2561)** | plasmid for BACTH assay carrying FlhF-T18 (Sputcn32_2561); Amp$^r$ | this study |
| **pUT18C FlhF (Sputcn32_2561)** | plasmid for BACTH assay carrying T18-FlhF (Sputcn32_2561); Amp$^r$ | this study |
| **pKT25 FipA (Sputcn32_2550)** | plasmid for BACTH assay carrying T25-FipA (Sputcn32_2550); Kan$^r$ | this study |
| **pKNT25 FipA (Sputcn32_2550)** | plasmid for BACTH assay carrying FipA-T25 (Sputcn32_2550); Kan$^r$ | this study |
| **pUT18 FipA (Sputcn32_2550)** | plasmid for BACTH assay carrying FipA-T18 (Sputcn32_2550); Amp$^r$ | this study |
| **pUT18C FipA (Sputcn32_2550)** | plasmid for BACTH assay carrying T18-FipA (Sputcn32_2550); Amp$^r$ | this study |
| **pKT25 FipA G106A (Sputcn32_2550)** | plasmid for BACTH assay carrying T25-FipA G106A (Sputcn32_2550); Kan$^r$ | this study |
| **pKNT25 FipA G106A (Sputcn32_2550)** | plasmid for BACTH assay carrying FipA G106A - T25 (Sputcn32_2550); Kan$^r$ | this study |
| **pUT18 FipA G106A (Sputcn32_2550)** | plasmid for BACTH assay carrying FipA G106A - T18 (Sputcn32_2550); Amp$^r$ | this study |
| **pUT18C FipA G106A (Sputcn32_2550)** | plasmid for BACTH assay carrying T18-FipA G106A (Sputcn32_2550); Amp$^r$ | this study |
| **pKT25 FipA L116A (Sputcn32_2550)** | plasmid for BACTH assay carrying T25-FipA L116A (Sputcn32_2550); Kan$^r$ | this study |
| **pKNT25 FipA L116A (Sputcn32_2550)** | plasmid for BACTH assay carrying FipA L116A - T25 (Sputcn32_2550); Kan$^r$ | this study |
| **pUT18 FipA L116A (Sputcn32_2550)** | plasmid for BACTH assay carrying FipA L116A - T18 (Sputcn32_2550); Amp$^r$ | this study |
| **pUT18C FipA L116A (Sputcn32_2550)** | plasmid for BACTH assay carrying T18-FipA L116A (Sputcn32_2550); Amp$^r$ | this study |
| **pKT25 FipA L125A (Sputcn32_2550)** | plasmid for BACTH assay carrying T25-FipA L125A (Sputcn32_2550); Kan$^r$ | this study |

| | | |
|---|---|---|
| **pKNT25 FipA L125A (Sputcn32_2550)** | plasmid for BACTH assay carrying FipA L125A - T25 (Sputcn32_2550); $Kan^r$ | this study |
| **pUT18 FipA L125A (Sputcn32_2550)** | plasmid for BACTH assay carrying FipA L125A - T18 (Sputcn32_2550); $Amp^r$ | this study |
| **pUT18C FipA L125A (Sputcn32_2550)** | plasmid for BACTH assay carrying T18-FipA L125A (Sputcn32_2550); $Amp^r$ | this study |

#### References Plasmids


Arroyo-Pérez EE, Ringgaard S. Interdependent Polar Localization of FlhF and FlhG and Their Importance for Flagellum Formation of *Vibrio parahaemolyticus*. Front Microbiol. 2021 Mar 17;12:655239. doi: 10.3389/fmicb.2021.655239. PMID: 33815347; PMCID: PMC8009987.

Heering J, Ringgaard S. Differential Localization of Chemotactic Signaling Arrays during the Lifecycle of Vibrio parahaemolyticus. Front Microbiol. 2016 Nov 2;7:1767. doi: 10.3389/fmicb.2016.01767. PMID: 27853457; PMCID: PMC5090175.

Hook JC, Blagotinsek V, Pané-Farré J, Mrusek D, Altegoer F, et al. A Proline-Rich Element in the Type III Secretion Protein FlhB Contributes to Flagellar Biogenesis in the Beta- and Gamma-Proteobacteria. Front Microbiol. 2020 Dec 15;11:564161. doi: 10.3389/fmicb.2020.564161. PMID: 33384667; PMCID: PMC7771051.

Iyer, Shankar Chandrashekar, Delia Casas-Pastor, David Kraus, Petra Mann, Kathrin Schirner, Timo Glatter, Georg Fritz, and Simon Ringgaard. 2020. "Transcriptional Regulation by σ Factor Phosphorylation in Bacteria." *Nature Microbiology* 5 (3): 395–406. https://doi.org/10.1038/s41564-019-0648-6.

Karimova G, Pidoux J, Ullmann A, Ladant D. A bacterial two-hybrid system based on a reconstituted signal transduction pathway. Proc Natl Acad Sci U S A. 1998 May 12;95(10):5752-6. doi: 10.1073/pnas.95.10.5752. PMID: 9576956; PMCID: PMC20451.

Lassak J, Henche AL, Binnenkade L, Thormann KM. ArcS, the cognate sensor kinase in an atypical Arc system of Shewanella oneidensis MR-1. Appl Environ Microbiol. 2010 May;76(10):3263-74. doi: 10.1128/AEM.00512-10. Epub 2010 Mar 26. PMID: 20348304; PMCID: PMC2869118.

Milton, D. L., O'Toole, R., Horstedt, P., and Wolf-Watz, H. (1996). Flagellin a is essential for the virulence of *Vibrio anguillarum*. J. Bacteriol. 178, 1310–1319. doi: 10.1128/jb.178.5.1310-1319.1996

Rossmann, F. et al. The role of FlhF and HubP as polar landmark proteins in *Shewanella putrefaciens* CN-32. Mol. Microbiol. 98, 727–742 (2015)

Schuhmacher JS, Rossmann F, Dempwolff F, Knauer C, Altegoer F, et al. MinD-like ATPase FlhG effects location and number of bacterial flagella during C-ring assembly. Proc Natl Acad Sci U S A. 2015 Mar 10;112(10):3092-7. doi: 10.1073/pnas.1419388112. Epub 2015 Mar 2. PMID: 25733861; PMCID: PMC4364217.

**Table S5: Oligonucleotides used in this study**

**Oligos used for *V. parahaemolyticus* work**

| Name | **Sequence** | **Purpose** |
|---|---|---|
| VP2224-del-a | CCCCC tctaga ACGTTGTCATGCTTGGTGAAAGCA | KO |
| VP2224-del-b | AGTCTCTTCAGCCATCGTCATTC | KO |
| VP2224-del-c | gaatgacgatggctgaagagact cgacgataaagagaataaaaagaagc | KO |
| VP2224-del-d | CCCCC tctaga ACGCGACGCTGCTGACCCGCAGAA | KO |
| VP2224-check | acaaactccgtggggatgaatac | CP |
| vp2224 AA1-6/28-end w/o Stop | ccccc ctcaga atg gctgaagagacttttctgcgc | KI |
| pUT18C/pKT25-vp2234-cw | ccccc tctaga G aaaataaagcgatttttgccaaagac | BACTH |
| pUT18C/pKT25-vp2234-ccw | ccccc ggtacc ctagagtccttcgttgtcactg | BACTH |
| vpa1548-del-d | Ccccc ctcgag TTATGTGTTCCGCCTTCCTCTC | CP |
| vpa1548-del-chk | aagtagccacatcccaaacgc | CP |
| VP2191-del-d | ccccc tctaga GACAATGCGCTGCACGGAAT | CP |
| VP2191-del-chk | gatggaaaacggctacacca | CP |
| del vp2234(FlhF)-d | CCCCC tctaga GAATACATGCTACGAGCTCAAGG | CP |
| del vp2234(FlhF)-chk | GTTTACGGCATGATTGATGGCG | CP |
| vp2224-Gly110Ala-cw | gagcaaccaaaatggtgcagttaGCGgctgatatcaacgagctaatcg | KI |
| vp2224-Gly110Ala-ccw | CGATTAGCTCGTTGATATCAGCcgcTAACTGCACCATTTTGGTTGCT C | KI |
| vp2224-Glu126Ala-cw | agagtgtgaactgccaaaagcaGCAgcagagttgatgctctctttgc | KI |
| vp2224-Glu126Ala-ccw | GCAAAGAGAGCATCAACTCTGctgCTGCTTTTGGCAGTTCACACTC T | KI |
| vp2224-Leu129Ala-cw | tgaactgccaaaagcagaagcagag GC gatgctctctttgcagaaaaaactg | KI |
| vp2224-Leu129Ala-ccw | CAG TTT TTT CTG CAA AGA GAG CAT CGC CTC TGC TTC TGC TTT TGG CAG TTC A | KI |
| C-term sfGFP-vp2224-a | CCCCC actagt ATGGCTGAAGAGACTTTTTTATCTGTAC | KI |
| C-term sfGFP-vp2224-b | gagctcgaggatgtc TCGTCGACGCCCACGTGG | KI |
| C-term sfGFP-vp2224-c | gacatcctcgagctc atgagcaaaggagaagaacttttcac | KI |
| C-term sfGFP-vp2224-d | tta tttgtagagctcatccatgcc | KI |
| C-term sfGFP-vp2224-e | ggcatggatgagctctacaaa taa AGAGAATAAAAAGAAGCTTCGG | KI |

| | | |
|---|---|---|
| C-term sfGFP-vp2224-f | ccccc gcatgc TTTGTTTGTCGATTGCTGTTAGTGG | KI |
| del AA7-27 vp2224-b | AAAAGTCTCTTCAGCCATCGTCATTC | KO |
| del AA7-27 vp2224-c | GAATGACGATGGCTGAAGAGACTTTT CTGCGCATTCGTGCTAGTTTGC | KO |
| vp2224-cw-pBAD | CCCCC tctaga atggctgaagagactttttatctg | KI |
| vp2224-ccw-pBAD | CCCCC gcatgc ttatcgtcgacgcccacg | KI |
| vp2224 cw restore deletion | ACCTATAATTGGCTGAATGACG ATGGCTGAAGAGACTTTTTATCTGTAC | KI |
| downstream vp2224 cw | AGAGAATAAAAAGAAGCTTCGGC | CP |
| **pUT18/pKNT25-vp2224-cw** | ccccc TCTAGA atggctgaagagactttttatctgtac | BACTH |
| **pUT18/pKNT25- tr-vp2224-cw** | ccccc TCTAGA ATG cgcattcgtgctagtttgc | BACTH |
| **pUT18/pKNT25-vp2222 -ccw** | ccccc GGTACC CG tcgtcgacgcccacgtg | BACTH |
| **pUT18C/pKT25-vp2224-cw** | ccccc tctaga G gctgaagagactttttatctgtac | BACTH |
| **pUT18C/pKT25-vp2224-ccw** | ccccc ggtacc ttatcgtcgacgcccacgtg | BACTH |
| **tr2224 put18C cw** | ccccc tctaga G ATG cgcattcgtgctagtttgcaaaa | BACTH |
| **sfGFP-1-ccw** | ccccc tctaga tttgtagagctcatccatgccatg | BACTH |
| **vp2224 C-term PhoA-LacZ cw** | CCCCC tctaga g atggcccggacaccagaaatg | TOP |
| **end -LacZ w/o STOP ccw** | gcgccattcgccattcaggctgc | TOP |
| **LacZ to vp2224 w/o ATG** | CCT GAA TGG CGA ATG GCG C GCT GAA GAG ACT TTT TTA TCT GTA CC | TOP |
| **end vp2224 ccw** | CCCCC aagctt ttatcgtcgacgcccacgtgg | TOP |
| vp2224 ccw restore deletion | GCCGAAGCTTCTTTTTATTCTCT TTATCGTCGACGCCCACGTG | KI |

**Oligonucleotides used for *Pseudomonas* and *Shewanella* work**

| **Name** | **Sequence** | **Purpose** |
|---|---|---|
| **M13** | TGTAAAACGACGGCCAGTCC | CP/SP |
| **M13r** | CACACAGGAAACAGCTATGACC | CP/SP |
| **flhF1-flhG1 fwd** | GCGCTGAGTGTGTTGATCCAAA | CP |
| **EcoRV FliM1 N-term fwd** | GCGAATTCGTGGATCCAGATGCTCATTGAAGATGCTCTCCTG | KI |
| **EcoRV FliM1 N-term rev** | GCCAAGCTTCTCTGCAGGATAATAAAACTGCGGCCCACTTCC | KI |

| | | |
|---|---|---|
| **Check-GFP FliM1-fwd** | GCAGTTCAGATGAGTCATCCTC | CP |
| **Check-GFP FliM1 KO-rev** | GACATTTTGGCAGTTGATGCGAC | CP |
| **OL FliM1 GFP rev** | GAAAAGTTCTTCTCCTTTGCTGCTGCCTAATTCAGATATATCTCTAGCTTTGCCTTTGC | KI |
| **OL FliM1 GFP fwd** | GGATGAGCTCTACAAAGGATCCTAAGGTGAAGCAAGATGAGCACAGAAGATA | KI |
| **EcoRV FlhF C-term fwd** | GCGAATTCGTGGATCCAGATGCAAGAAATGGTTGGACAGCCT | KI |
| **EcoRV FlhF C-term rev** | GCCAAGCTTCTCTGCAGGATGCCACATCTAAAAATCGGTCGG | KI |
| **Check-FlhF-FLAG-fwd** | GCATCAGTCAATGCAAGCAACC | CP |
| **OL-FlhF-Venus rev** | CACGCTGCCCTCAAATGCACAGGCCATATTATCTG | KI |
| **OL_Venus fwd** | GCATTTGAGGGCAGCGTGAGCAAGGGCGAGGAGCTGTT | KI |
| **OL_Venus rev** | GTCATAACTTTACTTGTACAGCTCGTCCATGCC | KI |
| **OL-FlhF-Venus fwd** | TACAAGTAAAGTTATGACCCTGGATCAAGCAAG | KI |
| **FlhF-Ven Seq_Primer** | GCTGAGTTAGTACGAGCACTAC | SP |
| **FlhG-Ven Seq_Primer** | CGATATTATTGTCCGTGGGCCT | SP |
| **FlhF-Ven Seq_Primer fwd** | GCTGTTGTAGTTGTACTCCAGC | SP |
| **EcoRV FlhF C-term rev** | GCCAAGCTTCTCTGCAGGATGCCACATCTAAAAATCGGTCGG | KI |
| **EcoRV-2550-GFP-fwd** | GCGAATTCGTGGATCCAGATGCCATCAATAACGGAAAAGGGG | KI/ CP |
| **OL-2550-GFP-rev** | GAAAAGTTCTTCTCCTTTGCTCAGTTCCAGAATATCTTTACGATGTAACCGGATCAATAATTCAGC | KI/CP |
| **OL-2550-GFP-fwd** | GGATGAGCTCTACAAAGGATCCTAACGAAGTGTAGGGGCTAAGACG | KI |
| **EcoRV-2550-GFP-rev** | GCCAAGCTTCTCTGCAGGATGCCTTTGTTTATATGCTCGACGG | KI |
| **Check-2550-GFP-fwd** | CGATGAAGAATGGGCTGAACTC | KI/CP |
| **Check-2550-GFP-rev** | CGAAGGATGCGAGAATGACGAA | KI/CP |
| **OL-2069_FlhF-rev** | AATCTTCACTAGCATCCCCGTACATTGAACTC | KI |
| **OL-FlhF-Ven-fwd** | GGGATGCTAGTGAAGATTAAACGATTTTTTGCCAAAGAC | KI |
| **OL-FlhF-Ven-rev** | AACATTAGCTTACTTGTACAGCTCGTCCATGC | KI |
| **OL-2068-fwd** | TACAAGTAAGCTAATGTTTTAGGGTCTTACGCG | KI |
| **BACTH 2550 pkT25 fwd** | CAGGGTCGACTCTAGAGGGCGATGAATTTTTGATCGCGG | BACTH |
| **BACTH 2550 pkT25 rev** | TTAGTTACTTAGGTACCCGGGGTTTACGATGTAACCGGATCAATAATTCAGC | BACTH |

| | | |
|---|---|---|
| **BACTH 2550 fwd** | CTGCAGGTCGACTCTAGAGGGCGATGAATTTTTGATCGCGG | BACTH |
| **BACTH 2550 rev** | GAGCTCGGTACCCGGGGTTTACGATGTAACCGGATCAATAATTCAGC | BACTH |
| **OL_FliM1 mCh rev** | TTTGTATAACTCATCCATACCA | KI |
| **FlhF-Ven Seq_Primer rev** | GCTGGAGTACAACTACAACAGC | SP |
| **OL-GFP-fwd** | AGCAAAGGAGAAGAACTTTTC | KI |
| **OL-GFP-rev** | GGATCCTTTGTAGAGCTCATCC | KI |
| **OL -mCherry fwd** | GTTTCCAAAGGGGAAGAGGACA | KI |
| **pKT25-for** | CACTGACGGCGGATATCGACATGTT | CP/SP |
| **pKT25-rev** | CCGCCGGACATCAGCGCCATTC | CP/SP |
| **pUT18-for** | CCAGGCTTTACACTTTATGCTTCC | CP/SP |
| **pUT18-rev** | GACGCGCCTCGGTGCCCACTGC | CP/SP |
| **pKNT25-for** | CCCAGGCTTTACACTTTATGCTTCC | CP/SP |
| **pKNT25-rev** | GTTTTTTTCCTTCGCCACGGCCTTG | CP/SP |
| **pUT18C-for** | CGGCGTGCCGAGCGGACGTTCG | CP/SP |
| **pUT18C-rev** | TCAGCGGGTGTTGGCGGGTGTC | CP/SP |
| **FlhF Seq_Primer fwd** | GCCCACTTTGGATCAACACACT | SP |
| **FlhF Seq_Primer rev** | CGTGCTCACAAAACTCGATGAA | SP |
| **EcoRV FliFG1 KO fwd** | GCGAATTCGTGGATCCAGATGCCGAAAACTTGTGGCTGAAAA | KO |
| **OL- FliFG1 KO rev** | ATCGCCACCCCCGACAATCATTTCTGTGCTC | KO |
| **OL- FliFG1 KO fwd** | ATTGTCGGGGGTGGCGATGAGTTCCTCTAAT | KO |
| **EcoRV FliFG1 KO rev** | GCCAAGCTTCTCTGCAGGATGCAACCTAATAGTCACTGCTTG | KO |
| **OL-fipA L118A rev** | AGCTTCAGCTTTGGGCGCTTCACA | KI |
| **OL-fipA L118A fwd** | ATAAAAGAGTGTGAAGCGCCCAAA | KI |
| **OL-fipA G106A rev** | TTCATCGACTCCCGCGGCAAGTCC | KI |
| **OL-fipA G106A fwd** | AAAATGGTCGGACTTGCCGCGGGA | KI |
| **OL-PPfipA G104A rev** | CATCGATACTCGCAGCCATCCC | KI |
| **OL-PPfipA G104A fwd** | GCTGGTGGGGATGGCTGCGAGT | KI |
| **OL-PPfipA L123A rev** | ACACCTTGCTCATCGCCTCCGC | KI |
| **OL-PPfipA L123A fwd** | GGCCGAGGCGGAGGCGATGAGC | KI |
| **EcoRV-flhF KO-fwd** | GCCAAGCTTCTCTGCAGGATGCATAGGCGTCGGTGATTGAGG | KO |
| **OL-flhF KO-rev** | TAAGTGAAGGCATTTGAGTAGAGTTATGACCCTGG | KO |
| **OL-flhF KO-fwd** | CTCAAATGCCTTCACTTATGCGTCCTCTACTGG | KO |

| | | |
|---|---|---|
| **EcoRV-flhF KO-rev** | GCGAATTCGTGGATCCAGATGCTAAGCATTCTCCTAAGCTTGTTG | KO |
| **OL-fipA L125A rev** | TAACCGGATCAAGGCTTCAGCTTC | KI |
| **OL-fipA L125A fwd** | GCTGAAGCTGAAGCCTTGATCCGG | KI |
| **EcoRV FlhF sub rev** | GCCAAGCTTCTCTGCAGGATGCTCGTCACATACAACGACTAG | KI |
| **BACTH 2550 L125A pkT25 rev** | TTAGTTACTTAGGTACCCGGGGTTTACGATGTAACCGGATCAAGGCTTCAGC | BACTH |
| **BACTH 2550 L125A rev** | GAGCTCGGTACCCGGGGTTTACGATGTAACCGGATCAAGGCTTCAGC | BACTH |
| **OL-FipA L125A-GFP-rev** | GAAAAGTTCTTCTCCTTTGCTCAGTTCCAGAATATCTTTACGATGTAACCGGATCAAGGCTTCAGC | KI |
| **BACTH FlhF pkT25 fwd** | CAGGGTCGACTCTAGAGAAGATTAAACGATTTTTTGCCAAAGACA | BACTH |
| **BACTH FlhF pkT25 rev** | TTAGTTACTTAGGTACCCGGGGCTCAAATGCACAGGCCATATTATCT | BACTH |
| **BACTH FlhF fwd** | CTGCAGGTCGACTCTAGAGAAGATTAAACGATTTTTTGCCAAAGACA | BACTH |
| **BACTH FlhF rev** | GAGCTCGGTACCCGGGGCTCAAATGCACAGGCCATATTATCT | BACTH |
| **BACTH FlhF GTG fwd** | CTGCAGGTCGACTCTAGAGGTGAAGATTAAACGATTTTTTGCCAAAG | BACTH |
| **EcoRV_FipA KO fwd** | GCGAATTCGTGGATCCAGATTTTTAGGTATCATTAACTTACGTGGTAATGT | KO |
| **OL-FipA KO rev** | ACACTTCGCTATTTACGATGATCGCCCATTAAAAATCCTTATGCA | KO |
| **OL-FipA KO fwd** | AAGGATTTTTAATGGGCGATCATCGTAAATAGCGAAGTGTAGGG | KO |
| **EcoRV-FipA KO rev** | GCCAAGCTTCTCTGCAGGATGAACTGATCGCCTTTGTTTATATGC | KO |
| **Check-FipA KO fwd** | AAGAAATGTCGCAGCCGTAGC | CP |
| **Check-FipA KO rev** | CCAGTTGCGACAATCTTCGGAG | CP |
| **OL-PPfipA L116A rev** | CATCAACTCCGCCTCGGCCTGGGTCGCGCCGCAGCTCTGGGT | KI |
| **OL-PPfipA L1164A fwd** | GAGTTGACCCAGAGCTGCGGCGCGACCCAGGCCGAGGCG | KI |
| **Check-PP_4331 (FipA) fwd** | GCTTACGAACAGAACGCAAGGC | CP |
| **Check-PP_4331 (FipA) rev** | GCAATACGTGATTTCGGTGCAG | CP |
| **EcoRV-PP_4331 KO-fwd** | GCGAATTCGTGGATCCAGATGCAGATGCACGCCAAACAGAAA | KO |
| **PP_4331 KO-OL-rev** | TCAAGGAGCTAGGATCAACTCAGATGTTCTCCAGC | KO |
| **PP_4331KO-OL-fwd** | TTGATCCTAGCTCCTTGACGGGGTACCCTCG | KO |
| **EcoRV-PP_4331 KO-rev** | GCCAAGCTTCTCTGCAGGATGCATGAATTGCCTGTACAACACCA | KO |
| **Check-PP_4331KO-fwd** | GCGAAACGATCGATCAGGTCGA | CP |
| **Check-PP_4331KO-** | GCACCGTAATCGAACACATGTG | CP |

| | | |
|---|---|---|
| **rev** | | |
| **EcoRV-PP_4331-GFP-fwd** | GCGAATTCGTGGATCCAGATGCAGATGCACGCCAAACAGAAA | KI |
| **PP_4331-GFP-OL-rev** | GAAAAGTTCTTCTCCTTTGCTCAGTTCCAGAATATCAGGAGCCCGG TACACCTTGCTC | KI |
| **PP_43310-GFP-OL-fwd** | GGATGAGCTCTACAAAGGATCCTGACGGGGTACCCTCGGCAGCA | KI |
| **EcoRV-PP_4331-GFP-rev** | GCCAAGCTTCTCTGCAGGATGCATGAATTGCCTGTACAACACCA | KI |
| **EcoRV-FlhF-mCh-fwd** | GCGAATTCGTGGATCCAGATGCATGGACAGCTTCCGTATCGG | KI |
| **FlhF-mCh-OL-rev** | CTCTTCCCCTTTGGAAACGCTGCCACCCGCTCGCCGTGGGTTGTGA | KI |
| **FlhF-mCh-OL-fwd** | ATGGATGAGTTATACAAATGACCATGAAGCGTGTGCAAAG | KI |
| **EcoRV-FlhF-mCh-rev** | GCCAAGCTTCTCTGCAGGATGCCAACACACGGAAACGGTTCA | KI |
| **Check-PP_4343 KO-fwd** | GCCTGAAATCGAGCCGATCGAA | CP |
| **Check-PP_4343 KO-rev** | GCGTCGGTAATCGAGGTAGGTT | CP |
| **BACTH PP FipA pkT25 fwd** | CAGGGTCGACTCTAGAGATCCTAGAGGTTGCTGTCATCT | BACTH |
| **BACTH PP FipA pkT25 rev** | TTAGTTACTTAGGTACCCGGGGAGGAGCCCGGTACACCTTGCTC | BACTH |
| **BACTH PP FipA fwd** | CTGCAGGTCGACTCTAGAGATCCTAGAGGTTGCTGTCATCT | BACTH |
| **BACTH PP FipA rev** | GAGCTCGGTACCCGGGGAGGAGCCCGGTACACCTTGCTC | BACTH |
| **BACTH PP FlhF pkT25 fwd** | CAGGGTCGACTCTAGAGCAAGTTAAGCGATTTTTCGCCGC | BACTH |
| **BACTH PP FlhF pkT25 rev** | TTAGTTACTTAGGTACCCGGGGACCCGCTCGCCGTGGGTTGTGA | BACTH |
| **BACTH PP FlhF fwd** | CTGCAGGTCGACTCTAGAGCAAGTTAAGCGATTTTTCGCCGC | BACTH |
| **BACTH PP FlhF rev** | GAGCTCGGTACCCGGGGACCCGCTCGCCGTGGGTTGTGA | BACTH |
| **EcoRV FlhF sub fwd** | GCGAATTCGTGGATCCAGATGCATCAGTCAATGCAAGCAACC | KI |
| **Check-FlhF KI/O-fwd** | GCCACTGGGTAGTGTCGTAAAA | CP |
| **OL-FlhF D328A rev** | CCCCATACCAGCGGTGGCTATCAATAC | KI |
| **OL-FlhF D328A fwd** | AAGCTAGTATTGATAGCCACCGCTGGT | KI |
| **EcoRV PPFlhF sub fwd** | GCGAATTCGTGGATCCAGATGCATGTTCTGGCGTATCAGGAA | KI |
| **OL-FlhF K235A rev** | GCGCGCGGCCAGCGCGGCCAGGGT | KI |
| **OL-FlhF K235A fwd** | GGCAAGACCACCACCCTGGCCGCGCTGGCCGCG | KI |
| **EcoRV PPFlhF sub rev** | GCCAAGCTTCTCTGCAGGATGCATGCTACCCATGTCTGTTCT | KI |
| **Check-PP_4343 KI-fwd** | GCTACCAGTGATTACCCTGGAG | CP |

| | | |
|---|---|---|
| **EcoRV PPFlhF sub 1 rev** | GCCAAGCTTCTCTGCAGGATGCGTCGGTAATCGAGGTAGGTT | KI |
| **KT2440 FlhF Seq primer rev** | GCTGGTGAGCATGGACAGCTTC | SP |
| **EcoRV-FipA dTM fwd** | GCGAATTCGTGGATCCAGATGCCGTAGCTGCAAGTAAAGATG | KI |
| **OL-FipA dTM rev** | CTGCTTTTGTTCATCGCCCATTAAAAATCCTTATGC | KI |
| **OL-FipA dTM fwd** | GGCGATGAACAAAAGCAGTTGAGTAAATTACGTAATAAAGTTG | KI |
| **OL-PP_FipA dTM rev** | GCTGTAGTTCTCTAGGATCAACTCAGATGTTCTCC | KI |
| **OL-PP_FipA dTM fwd** | ATCCTAGAGAACTACAGCAAGCGCCAGCGCG | KI |

**Abbreviations**: fwd: forward; rev: reverse; KO: knock-out (deletion) primer; KI: knock-in (integration) primer, CP: check primer, SP: sequencing primer, BACTH: Bacterial adenylate cyclase two-hybrid system primer
